# *Arabidopsis* seed stored mRNAs and translation regulation during post-harvest ripening and imbibition

**DOI:** 10.1101/2025.01.30.635699

**Authors:** J. Balarynová, B. Klčová, R. Čegan, K. Raabe, P. Krejčí, P. Bednář, D. Potěšil, V. Pustka, D. Tarkowská, V. Turečková, Z. Zdráhal, D. Honys, P. Smýkal

## Abstract

Seed germination marks the critical transition from dormancy to active growth, driven by environmental cues and water availability. This study explores translational regulation during germination by comparing two *Arabidopsis thaliana* accessions with contrasting dormancy phenotypes: Columbia (Col), which germinates readily, and Cape Verde Islands (Cvi), which exhibits deep dormancy. Using sucrose gradient centrifugation, we isolated monosomal and polysomal fractions from freshly harvested (FH), after-ripened (AR), and imbibed (IM) seeds. RNA-seq analysis revealed stage- and genotype-specific gene expression, with Col IM seeds displaying the highest number of expressed genes. We identified ∼14,000 mRNAs in FH seeds, increasing to 19,000 in Col and 17,000 in Cvi upon imbibition. Of these, 9,000 were shared, while 3,000 were accession-specific in monosomes. Enrichment analysis highlighted molecular pathways associated with translation and dormancy release. Analysis of RNA modifications identified N1-methyladenosine (m1A) as the predominant modification, with Col seeds exhibiting higher m1A levels than Cvi, peaking at three months post-harvest. m6A sequencing revealed distinct modification patterns between accessions, with the highest abundance of m6A-modified transcripts in IM seeds. Positional analysis of m6A peaks suggested a link to differential gene expression between Col and Cvi. Proteomic analysis identified ∼15,000 proteins, with translation-related proteins enriched in IM seeds. Notable differences between Col and Cvi were observed in both monosomal and polysomal fractions. RNA-binding proteins exhibited similar profiles in FH and AR stages but diverged significantly in IM seeds. Col-specific proteins were enriched in 40S ribosomes, processing bodies, and RNA-binding complexes.

These findings provide new insights into the molecular and translational dynamics underlying seed germination, advancing our understanding of dormancy release and early seedling establishment.

## Introduction

Angiosperm plants evolved seeds as a novel dispersal unit, sparing spores, originally used for this task in seedless plants. This improvement enabled further protection of vulnerable embryo and was accompanied by the rapid co-evolution of hidden alternation of generations, double fertilization and the establishment of completely new structures, such as flowers and seeds (Magnani 2018). Seeds are unique in terms of potential longevity, as some are reported to be alive after hundreds of years (Baskin and Baskin 2014; Sano *et al*. 2015). Dry seeds are well equipped to survive extended periods of unfavourable conditions and contain all components required for germination and seedling establishment until the seedling reaches an autotrophic state. Mature dry seeds are dispersed in a dormant state and seed germination is activated by hydration in relation to the environment. Dormancy evolved to control the seasonal timing of seed germination (Bewley 1997). Seed germination is a massively demanding process in terms of reserve mobilization, protein synthesis and the creation of new structures. One mechanism enabling the fast and massive activation of protein synthesis is the regulation of translation at the level of mRNA storage and coordinated activation of their translation. In many model systems including *Arabidopsis*, translation was identified as an important regulatory checkpoint being under both environmental and developmental control (Browning, Bailey-Serres 2015; Sajeev, Bai and Bentsink 2019). Moreover, more examples are emerging for the translational regulation in response to abiotic stress in plants, for example, heat, drought, light and osmotic stress (Juntawong and Bailey-Serres 2012; Kawaguchi *et al*. 2004; Kage *et al*. 2020).

Seed-stored proteins have been studied for a long time, while seed-stored mRNAs have been studied in more detail only recently (Bai *et al*. 2017, 2020; Holdsworth *et al*. 2008; Sajeev *et al*. 2019). Similarly, the degradation of specific mRNAs has also long been implicated in breaking seed dormancy (Dure and Waters 1965); the degradation of a particular subset of mRNAs might be a prerequisite to germination (Howell *et al*. 2009). These stored, long-lived mRNAs have been found in many angiosperms and are believed to be crucial for protein synthesis during germination (Basbouss-Serhal *et al*. 2015; Galland *et al*. 2014; Layat *et al*. 2014). A mystery about long-lived mRNAs that remains to be solved is how these are stored and protected in seeds (Sajeev *et al*. 2019). It has been shown that ribonucleoprotein complexes (mRNPs) accumulate mRNA during seed maturation (Ajtkhozhin *et al*. 1976). During the imbibition, the monosomes containing stored mRNAs become translation-competent polysomes as revealed by polysome profiles from seeds of various plants, such as castor bean (Marre *et al*. 1965), wheat (Spiegel *et al*. 1975). Recently, modifications of stored mRNAs during after-ripening have been proposed to be involved in the regulation of mRNA translation and/or mRNA degradation in the early steps of seed imbibition and then to subsequently program cell functioning toward germination or dormancy maintenance (Sano *et al*. 2020). How mRNAs are protected for such a long period is unknown, but it is likely due to association with RNA-binding proteins (Dedow and Bailey-Serres 2019). Transcriptional changes observed in imbibed seeds during dormancy cycling also indicate that an important molecular event in dormancy loss is increased expression of the translation machinery (Gallant *et al*. 2014; Buijs *et al*. 2019). The role of translational control was shown in *Arabidopsis* pollen tube growth (Lin *et al*. 2014) and seeds (Bai *et al*. 2017, 2020; Basbouss-Serhal *et al*. 2015). *Arabidopsis* seeds display physiological dormancy type (Finch-Savage and Leubner-Metzger 2006) based on the endogenous block regulated by plant hormone interactions, the balance of abscisic acid (ABA) and gibberellins (GAs), which is usually broken by a relatively short period of cold stratification or dry seed storage and room temperature. However, there is a large natural variation in *Arabidopsis* seed dormancy regulated by *DELAY OF GERMINATION 1* (*DOG-1*) locus (Alonso-Blanco *et al*. 2003; Bentsink *et al*. 2010) and related to environmental conditions and displays a latitudinal gradient (Vidigal *et al*. 2016). Commonly used accessions, such as Columbia (Col) or Landsberg (Ler), are non-dormant, while others like Cape Verde Islands (Cvi) have strong dormancy (Ali-Rachedi *et al*. 2004; Cadman *et al*. 2006; Finch-Savage *et al*. 2007). Subsequently, in natural conditions, these genotypes differ in life strategies starting from germination through flowering and seeds (Martinez-Berdeja *et al*. 2020). *Arabidopsis* seeds display dormancy cycling related to the environmental conditions, resulting in the establishment of a soil seed bank (Donohue 2002). Interestingly, different maternal growth conditions can result in variable dormancy phenotypes of the same genotype (Finch-Savage and Footitt, 2019; Iwasaki, Penfield, Lopez-Molina 2022) with differences being in translation-related transcripts (Buijs *et al*. 2019). The most abundant modification of eucaryotic mRNA is m6A methylation (Shi *et al*. 2017), which was suggested to play a role in plant development (Zhong *et al*. 2008) and regulation of gene expression (Luo *et al*. 2014). Moreover, m6A modification can influence RNA stability, decay, transport, splicing or translation (Haussmann *et al*. 2016; Frye *et al*. 2018; Yang *et al*. 2018; Luo *et al*. 2020). Both, positive (Meyer *et al*. 2015; Wang *et al*. 2015; Shi *et al*. 2017) and negative (Choi *et al*. 2016; Qi *et al*. 2016) effects of m6A modification on RNA translation were reported. A writer, a protein complex of the N6-adenosine methyltransferase A (MTA), N6-adenosine methyltransferase B (MTB), and the FKBP Interacting protein 37 (FIP37), is responsible for the deposition of m6A modification (Reichel *et al*. 2019). Furthermore, the VIRILIZER (VIR) and HAKAI proteins were discovered to be a part of the writer complex and a decrease of their gene expression led to a reduction of a level of m6A modifications (Růžička *et al*. 2017). The effect of the modification is mediated by m6A-binding proteins called readers represented by YTH domain family 2 (YTHDF2). Finally, the modification-removing proteins, erasers, such as Alpha-ketoglutarate-dependent dioxygenase (AlkB) and AlkB-homology (AlkBH) family proteins, can remove m6A modification (Zaccara *et al*. 2019). The position of m6A modification is not random. The sites around start and stop codons, and 3’UTRs are the most preferred in mammalian cells (Dominissini *et al*. 2012; Meyer *et al*. 2012). The significant m6A modification enrichment in the 3’UTR, at stop, start codons, and lower amount in the CDS was shown in *Arabidopsis* (Schwartz *et al*. 2013; Luo *et al*. 2014; Wan *et al*. 2015; Luo *et al*. 2020).

In this study, we have compared the mRNAs and associated proteins stored in monosomes or polysomes at the stages of seed maturity, post-ripening and early germination using two *Arabidopsis* genotypes differing in dormancy level. Moreover, we isolated and investigated transcripts with m6A modification and identified differences between stages and genotypes.

## Material and methods

### Plant material

Plants of *Arabidopsis thaliana* Columbia-0 (Col) and Cape Verde Islands (Cvi, accession number N8580) were grown in 4×4cm pots irrigated with tap water and fertilized weekly with standard nutrient solution (Krystalon, CZ) in a growth chamber at 22/18°C (day/night) under a 16-h day/8-h night photoperiod of artificial light (150 μmolm^−2^s^−1^) and 70% relative humidity. Seeds were harvested upon maturation and sampled as dry freshly-harvested (FH) or 3 months (100 days) since harvest labelled as after-ripened (AR) being stored in the dark and at room temperature in paper bags. In addition, 3-month-old seeds were imbibed and germinated for 48 hours (IMB) on Whatman 1 filter paper in 90 mm Petri dishes at 12/12h light/dark regime in 23C. All experiments were conducted in the replicates.

### Polysome profiling

The polysome profiling method (Mustroph *et al*. 2009; Mašek *et al*. 2011; Bai *et al*. 2020) was modified. Seed material, dry or imbibed, was collected, weighted and immediately frozen in liquid nitrogen. Samples were either used right away or stored in -80 °C. Frozen samples were ground to powder using a mortar and pestle pre-cooled with liquid nitrogen. Polysome extraction buffer (400 mM Tris-HCl [pH 9.0], 200 mM KCl, 25 mM EGTA [pH 8.3], 36 mM MgCl_2_, 5 mM DTT, 5 mM PMSF, 25 µg/mL cycloheximide, 25 µg/mL chloramphenicol, 0.8 % mercaptoethanol) was added in 10:1 ratio (buffer volume: sample weight). The resulting powder was transferred to a cool 50 mL falcon tube and left to melt on ice. The melted sample was further homogenized by pushing through 21G needle (B. Braun, Germany) several times. The homogenate was centrifuged at 16,000 x *g* for 15 minutes in 2 mL Eppendorf Safe-lock tubes. The supernatant was transferred to a new tube and centrifuged again. The final supernatant was loaded on top of the continuous 10 to 45% sucrose gradient prepared according to Mustroph *et al*. (2009) and formed using the Gradient Master^TM^ 108 (BioComp Instruments, Canada) set for 10 to 45 % w/v short cap sucrose gradient program. Formed gradients were placed at 8 °C overnight. The appropriate amount of ultrapure sucrose was dissolved in warm DEPC-treated ddH2O up to 45 mL and mixed with 5 mL of autoclaved 10x salt solution (0.4 M Tris-HCl [pH 8.4], 0.2 M KCl, 0.1 M MgCl_2_) and cycloheximide (final concentration 50 µg/mL). Ice-cold sucrose solutions were loaded in a 13.2 mL open-top thin-wall polypropylene ultracentrifuge tube (Beckman, USA) where 45 % sucrose solution was placed below 10 % sucrose solution using the underlay approach with a blunt needle. Tubes were sealed with rubber caps. Before sample loading, rubber caps were removed from the gradient together with 200 µL of the top to prevent spillover. 500 µL of the sample was then carefully loaded on top of the gradient. Sample tubes were placed in SW41 Ti Beckman rotor tube bucket, and centrifuged at 190,000 × *g* (∼39,000 rpm) at 4 °C in the Optima XPN Ultracentrifuge (Beckman, USA) for 3 hours (maximum acceleration, no braking). For sample processing, Brandel Density Gradient Fractionation System SYN-202 Syringe pump (Brandel, MD, USA) was used together with Foxy R1 fraction collector (Teledyne ISCO, NE, USA). Absorbance signal output was recorded (flow speed 1.5 mL/min, sensitivity 0.5, baseline 10), using either UA-6 chart recorder system (Teledyne ISCO, NE, USA) or Clarity Chromatography Software (DataApex, Czech Republic) connected to the output of the UA-6 detector. Fractions were collected in 2 mL Eppendorf Safe-lock tubes, with the collector set for 30 seconds per tube, equal to 750 µL per fraction. Fractions collected from the sample gradients were used for protein or RNA extraction according to the protein/RNA preparation protocol (Mustroph *et al*. 2009). The polysome lysate input was treated identically to get the proteome/transcriptome reference.

### RNA isolation and analysis

For RNA extraction, 2 volumes of 8 M Guanidine-HCl and 3 volumes of 99.8 % ethanol were added to the samples, mixed well and left to precipitate RNA at -20°C overnight. Precipitate was then pelleted at 21 000 x *g* for 30 minutes at 4 °C with one 75 % ethanol wash. RNA was isolated using TRI Reagent (Sigma-Aldrich, CZ) followed by further purification using Norgen Plant/Fungi Total RNA Purification Kit (Norgen, Canada) from pooled (separately monosomal and polysomal fractions) samples and sent for RNA sequencing to Novogene Ltd. Cambridge, UK. The sequencing of the libraries was conducted on Illumina instrument with 150 bp paired-end reads (Illumina Inc. San Diego, CA, USA). mRNA read trimming on quality (Q30) and sequencing adaptor removal were done with Trimmomatic 0.32 (Bolger *et al*. 2014). Resulting high-quality reads from each library were mapped and quantified onto the *A.thaliana* reference genome GCF_000001735.4_TAIR10.1 with Araport11 annotation using RSEM (v1.3.3; Li and Dewey 2011) and with bowtie (v-1.0.0.) with default parameters. Quantified reads were used as input for differential expression analysis using the Bioconductor DESeq2 package (version 1.44.0; Love *et al*. 2014). Three replicates were used per each condition (Table S1). Transcripts were considered as differentially expressed when the adjusted P value was < 0.05 and log2 fold change was >±1. For the analysis of mono and polysome RNAseq data, the 90th percentile of expressed genes was used instead of differential expression analysis due to the lack of replicates for all samples. Upset plots were done with UpSetR (Gehlenborg 2019). The intersections and Venn diagrams were produced using the online tool available at https://bioinformatics.psb.ugent.be/webtools/Venn/.

### m6A RNA profiling and analysis

Frozen seeds were ground to a fine powder with liquid nitrogen using mortar and pestle. Total RNA was isolated using PureLink™ Plant RNA Reagent (Thermo Fisher Scientific, MA USA). Residual DNA was removed by DNase I (Top-Bio, Czech Republic) treatment followed by phenol/chloroform extraction. EpiQuick CUT and RUN m6A RNA Enrichment (MeRIP) kit (Epigentek, NY, USA) was used for m6A RNA profiling. As an input, we used 10,000 ng of total RNA and followed the manufactureŕs instructions with one modification. After binding beads with antibody and enzymatic digestion (step 2b), isolated RNA containing m6A was released using RNA Clean-Up and Concentration Micro-Elute Kit (Norgen, Canada). This procedure was used instead of RNA binding beads suggested in the manufacturer protocol of EpiQuick CUT and RUN m6A RNA Enrichment (MeRIP) kit. In the final step of RNA Clean-UP and Concentration Kit, RNA was eluted in 11 µl of Elution Solution A. The received fractions were submitted for RNA immunoprecipitation sequencing (RIP-seq) at Novogene Ltd., Munich, Germany (Table S1). Sequencing libraries were prepared at Novogene after sample quality check, RNA fragmentation, reverse transcription, dA-tailing, adapter ligation, and PCR amplification. Resulting sequencing libraries were sequenced on Illumina instrument with 150 bp paired-end reads (Illumina Inc., San Diego, CA, USA).

m6A-modified RNA read trimming and sequencing adaptor removal were done with Trimmomatic 0.32 (Bolger *et al*. 2014). Trimmed reads were aligned to *A. thaliana* reference genome similarly like mRNA reads and by STAR aligner (v2.7.7a; Dobin *et al*., 2013). m6A peaks were identified with Exomepeak2 package (https://github.com/ZW-xjtlu/exomePeak2) from bam files generating by RSEM mapping. Visualization and detailed analysis were done with R package CHIPseeker (Wang *et al*. 2022). Subsetting and combinations of dataset with DE results were done by bedtools v2.26.0 (Quinlan and Hall. 2010) and upset plots were done with UpSetR (Gehlenborg, 2019). 25% TSS and 15% TTS sequences for motif analysis were extracted from the reference genome according to CDS annotation file. The intersections and Venn diagrams were produced using the online tool available at https://bioinformatics.psb.ugent.be/webtools/Venn/. The enrichment analysis of the m6A-modified RNA fractions was performed using the ShinyGO 0.81 tool (Ge *et al*. 2020). The MEME Suite 5.5.7 (Bailey *et al*. 2015) was used to identify the motifs in the sequences of m6A-modified genes. The following MEME set-up was applied: the motif length: 6–10 nucleotides, single strand reading only.

### UHPLC-MS nucleoside analysis

RNA nucleosides (N-6-methyladenosine, N-1-methyladenosine, 5-methylcytidine and 8-oxoguanosine) were quantified using UHPLC-MS and protocol modified according to the method by Fleming *et al*. (2018). RNA was digested to nucleosides in 200 µL overnight reaction at 37°C with 100 U µL^-1^ of S1 Nuclease (Thermo Fisher Scientific) in reaction buffer (5x reaction buffer for S1 Nuclease). This was followed by the addition of 1 U µL^-1^ of FastAP Thermosensitive Alkaline Phosphatase (Thermo Fisher Scientific) in reaction buffer (10x FastAP Buffer) and the reaction incubated 1 h at 37 °C in the dark. Enzymes were removed by microfiltration (10 kDa, Amicon Ultra, Sigma-Aldrich). UHPLC-MS was performed using Waters (Milford, MA, USA) Acquity UHPLC system with mass spectrometry detection (Select Series Cyclic IMS, Waters). Chromatographic separations were performed on a Waters T3 column (1 mm×100 mm, 1.7 µM) and the column was operated at 45 °C. Mobile phases were water with 0.1 % formic acid (A) methanol with 0.1 % formic acid (B). The analytical gradient was: 0 min, 0.1 % B; 3.0 min, 0.1 % B; 22 min, 55 % B; 22.01 min, 97 % B; 23 min, 97 % B; 23.01 min, 0.1 % B. Flow rate was 0.1 mL min^-1^ and injection volume was 5 µL. The mass spectrometer was operated in positive ionization modes with capillary spray voltage at 3.5 kV. Source temperature was 150 °C and desolvation gas temperature 220 °C. Desolvation gas flow was 600 L h^-1^, cone gas flow 200 L h^-1^. Quantification used calibration curves generated with authentic standards for studied compounds, parameters of calibration curves are listed in Table S1.

### Proteomic analysis

To investigate the ribosome protein complexes, high-throughput liquid chromatography-mass spectrometry LC/MS was applied to both mono-/polysomal fractions (total of 36 samples, in triplicates) obtained by sucrose gradient fractionation (Col and Cvi genotypes, FH, AR and IM stages) identically as for RNAseq analysis. For protein extraction, 2 volumes of 99.8 % ethanol were added, mixed and left to precipitate proteins at 4 °C overnight. Precipitate was then pelleted at 12,000 x *g* for 30 minutes at 4 °C, twice washed with 80 % ethanol and air dried at room temperature. The pellets were stored at -80 °C. The pellets were solubilized by hot SDT buffer (4% SDS, 0.1 M DTT, 0.1 M Tris/HCl, pH 7.6) in thermomixer (Thermo Scientific). The protein mixture (ca 50 μg of total protein) was used for filter-aided sample preparation (FASP) described by Wisniewski *et al*. (2018) using 1 μg of trypsin (sequencing grade; Promega). The resulting peptides were analysed by LC-MS/MS using nanoELUTE system (Bruker, Germany) connected to timsTOF Pro mass spectrometer (Bruker, Germany). Before LC separation, tryptic digests were online concentrated and desalted using a trapping column (Acclaim PepMap 100 C18, dimensions 300 μm ID, 5 mm long, 5 μm particles, Thermo Fisher Scientific). After washing the trapping column with 0.1% formic acid (FA), the peptides were eluted (flow rate 300 nL/min) from the trapping column onto an analytical column (Aurora C18, 75μm ID, 250 mm long, 1.6 μm particles, Ion Opticks) by 90 min linear gradient program (3-30 % of mobile phase B; mobile phase A: 0.1% FA in water; mobile phase B: 0.1 % FA in 80 % ACN). The linear gradient program was followed by an intensive wash of the column by 80 % of the mobile phase B. The trapping and analytical column were equilibrated before sample injection into the sample loop. The analytical column was placed inside the Column Toaster (Bruker). According to the manufacturer’s instructions, its emitter side was installed inside the CaptiveSpray ion source (Bruker) with the column temperature set to 40 °C. MS/MS data were acquired in data-independent acquisition (DIA) mode with base method m/z range of 100-1700 and 1/k0 range of 0.6-1.6 V×s×cm^-2^. Enclosed DIAparameters.txt file defined m/z 400-1000 precursor range with equal windows size of 21 Th using two steps for each PASEF scan and cycle time of 100 ms locked to 100 % duty cycle.DIA data were processed in DIA-NN^2^ (version 1.8) in library-free mode against the modified cRAP database (based on https://www.thegpm.org/crap/; 111 sequences in total) and UniProtKB protein database for *Arabidopsis thaliana* (version 2021/11, number of protein sequences: 27,469).

No optional, but carbamidomethylation as fixed modification and trypsin/P enzyme with 1 allowed missed cleavage and peptide length 7-30 were set during the library preparation. False discovery rate (FDR) control was set to 1 % FDR. MS1 and MS2 accuracies as well as scan window parameters were set based on the initial test searches (median value from all samples ascertained parameter values). MBR was switched on. Protein MaxLFQ intensities reported in the DIA-NN main report file were further processed using the software container environment (https://github.com/OmicsWorkflows), version 4.6.3a. Protein groups were classified into total proteins, RNA-binding (Rbome) based on Zhang *et al*. (2023), RNA-binding (seed Rbome based on Sajeev *et al*. 2021), 40S and 60S ribosomal subunits (Scarpin *et al*. 2023), eIFs/eEFs/eRFs (Browning and Bailey-Serres, 2015), stress granules (Kosmacz *et al*. 2019), processing bodies (Xu and Chua, 2011) and PABP/ALBA/ECT (Belostotsky, 2003, Náprstková *et al*. 2021, Flores-Téllez *et al*. 2023) proteins.

### Isolation and analysis of ABA, GA and its metabolites

Analysis of ABA was performed according to the method described in (Turečková *et al*. 2009) with some modifications. Briefly, approximately 3 mg of plant tissue was extracted in 1 mL ice-cold methanol/water/acetic acid (10/89/1, v/v) containing 2 pmol of mixture of stable isotope-labeled internal standards ((-)-7′,7′,7′-^2^H_3_-phaseic acid; (-)-7′,7′,7′-^2^H_3_-dihydrophaseic acid; (-) - 8′,8′,8′-^2^H_3_-neophaseic acid; (+)-4,5,8′,8′,8′-^2^H_5_-ABAGE; (-)-5,8′,8′,8′-^2^H_4_-7′-OH-ABA (National Research Council, Saskatoon, Canada); (+)-3′,5′,5′,7′,7′,7′-^2^H_6_-ABA (Olchemim, Olomouc, Czech Republic). After 1 h of shaking in the dark at 4°C, the homogenates were centrifuged (36 670 x *g*, 10 min, 4°C), and the pellets were then re-extracted in 0.5 mL extraction solvent for 30 min. The combined extracts were purified by solid phase extraction (SPE) using Oasis^TM^ HLB columns (30 mg, 1 mL; Waters, Milford, CT, USA), then evaporated to dryness *in vacuo* and analysed by an Acquity UPLC® I-class system (Waters, Milford, MA, USA) combined with Xevo™ TQ-XS triple quadrupole mass spectrometer (Waters, Manchester, UK).

The sample preparation and analysis of gibberellins (GAs) was performed according to the method described in Urbanová *et al*. (2013) with some modifications. Briefly, tissue samples of about 5 mg FW were ground to a fine consistency using 2.7-mm zirconium oxide beads (Retsch GmbH, Haan, Germany) and MM 400 vibration mill at frequency of 27 Hz for 3 min (Retsch GmbH, Haan, Germany) with 1 mL of ice-cold 80 % acetonitrile containing 5 % formic acid as extraction solution. The samples were then extracted overnight at 4 °C using a benchtop laboratory rotator Stuart SB3 (Bibby Scientific Ltd., Staffordshire, UK) after adding internal gibberellins standards ([^2^H_2_]GA_1_, [^2^H_2_]GA_4_, [^2^H_2_]GA_9_, [^2^H_2_]GA_19_, [^2^H_2_]GA_20_, [^2^H_2_]GA_24_, [^2^H_2_]GA_29_, [^2^H_2_]GA_34_ and [^2^H_2_]GA_44_) purchased from OlChemIm, Czech Republic. The homogenates were centrifuged at 36 670 x *g* and 4 °C for 10 min, and corresponding supernatants were further purified using reversed-phase and mixed-mode SPE cartridges (Waters, Milford, MA, USA) and analysed by ultra-high performance liquid chromatography-tandem mass spectrometry (UHPLC-MS/MS; Micromass, Manchester, UK). GAs were detected using multiple-reaction monitoring mode of the transition of the ion [M–H]^-^ to the appropriate product ion. Masslynx 4.2 software (Waters, Milford, MA, USA) was used to analyze the data, and the standard isotope dilution method (Rittenberg and Foster, 1940) was used to quantify the GAs levels.

## Results

### ABA declines of during germination, whereas GA levels remain stable in both genotypes

Two *Arabidopsis* accessions differing in seed dormancy were analysed. Columbia (Col) is non-dormant and its seeds are capable of germination immediately after shedding, while Cape Verde Islands (Cvi) has deeply dormant seeds requiring after-ripening (minimum of 90-100 days) to be able to germinate. In our conditions, freshly harvested Col and Cvi seeds displayed 60% and 0% germination, respectively, while after 3 months (90 days) Cvi seeds germination was around 80%. The levels of ABA, phaseic acid (PA), dihydrophaseic acid (DPA), 7’-hydroxy-ABA (7′-OH-ABA) and neophaseic acid (neoPA) were the lowest in germinating seeds of both Col and Cvi. On the other hand, the ABA glycosyl ester (ABA-GE) was detected only in germinating seeds of both genotypes (Fig. 1). Dihydrophaseic acid (DPA), produced from PA by the 8’-hydroxylation pathway, was the major ABA catabolic product in both genotypes.

**Fig. 1:**
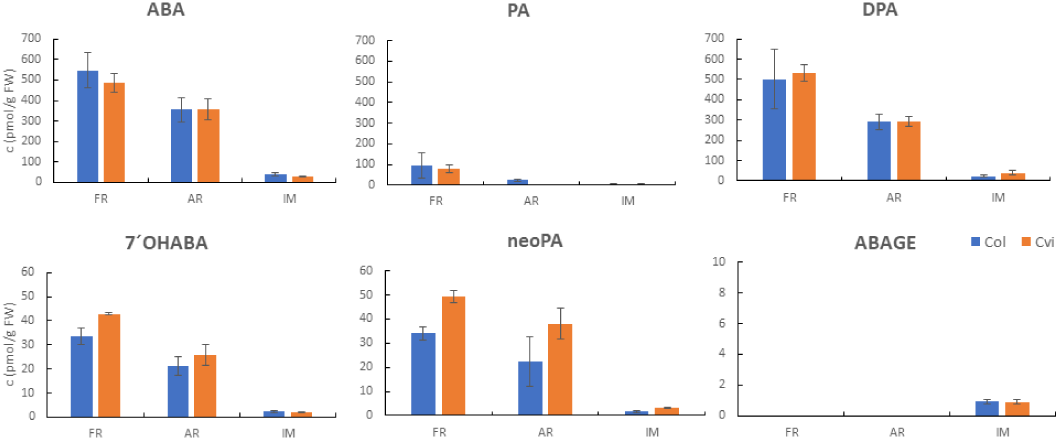
Quantification of ABA, phaseic acid (PA), dihydrophaseic acid (DPA), 7’-hydroxy-ABA (7′-OH-ABA), neophaseic acid (neoPA) and ABA glycosyl ester (ABA-GE) levels in dry freshly harvested (FH), dry after-ripened (AR) and imbibed (IM) Col and Cvi seeds. Data expressed mean ± SD of three measurements.

The levels of bioactive GAs, their biosynthetic precursors and catabolites were determined in dry (freshly harvested and after-ripened) and germinating Col and Cvi seeds. GA_20_ was the most abundant GA precursor belonging to the 13-hydroxylated biosynthetic pathway in dry and germinating seeds of both genotypes. GA_34_ was the only precursor of the 13-non-hydroxylated biosynthetic pathway determined. GA_1_ (the 13-hydroxylated pathway) and GA_4_ (the 13-non-hydroxylated pathway) were found to be the main bioactive GAs in the germinating Col and Cvi seeds (Fig. 2). In Cvi seeds, GA_5_ (the 13-hydroxylated pathway) was also abundant.

**Fig. 2:**
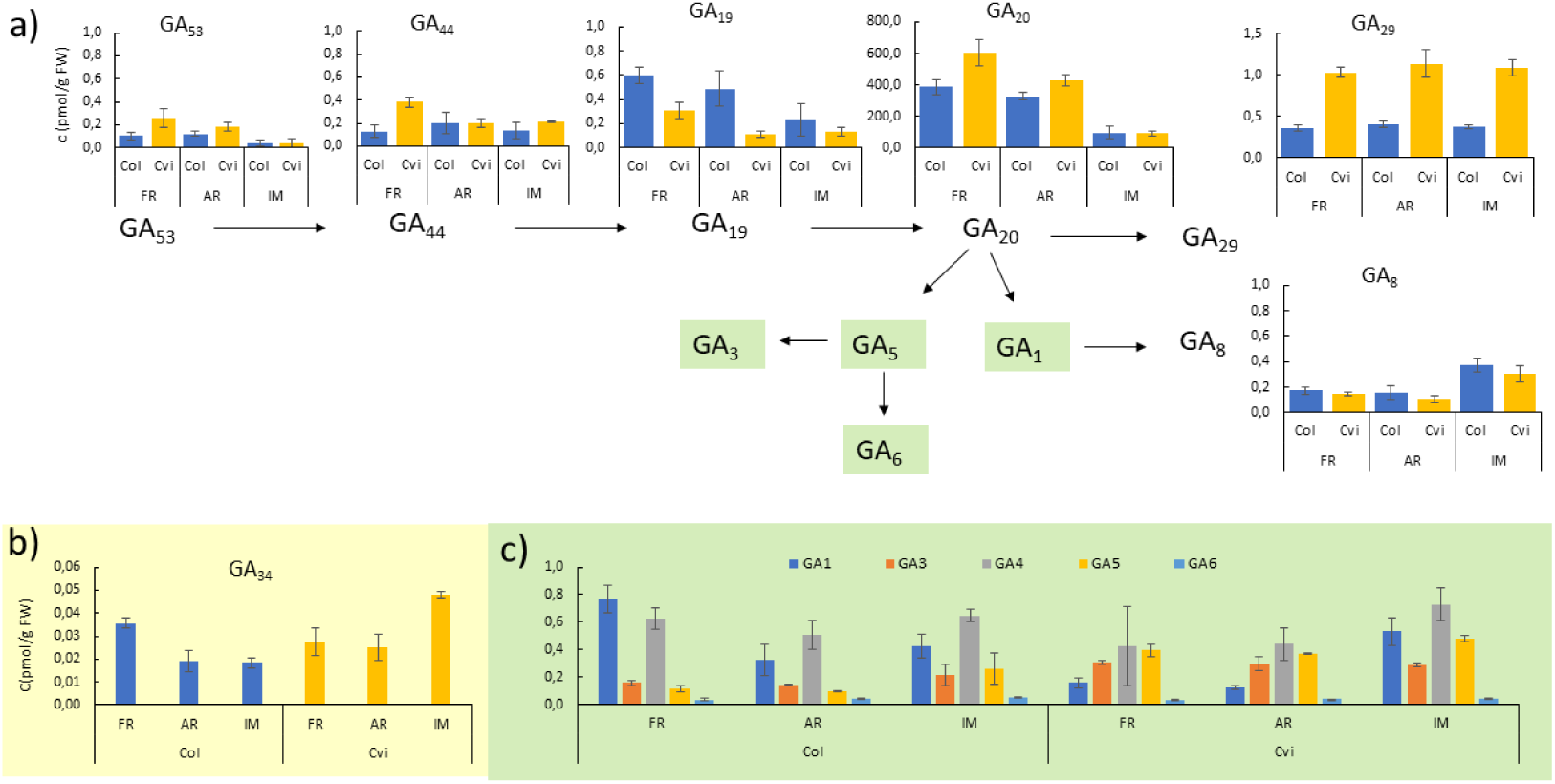
**The levels of gibberellins (GAs)** in dry freshly harvested (FH), dry after-ripened (AR) and imbibed (IM) Col and Cvi seeds, a) GAs **belonging to the 13-hydroxylation** (leading to the production of GA_1_, GA_3_, GA_5_, GA_6_) gibberellin metabolic pathway, v) GA_34_, the GA belonging to the 13-non-hydroxylated (leading to the production of GA_4_ and GA_7_) biosynthetic pathway, **c) the bioactive GAs belonging to the 13-hydroxylated (GA_1_, GA_3_, GA_5_, GA_6_) and 13-non-hydroxylated (GA_4_) pathway** Data expressed mean ± SD of three measurements.

#### ABA and GA metabolic gene expression showed a similar trend between Col and Cvi seeds

The RNA sequencing data were searched for levels of gene expression of genes encoding enzymes of ABA and GA biosynthesis and catabolism. The data showed that genes encoding enzymes involved in the formation of GA precursors (*ent*-kaurene synthase, GA 20-oxidase and GA 3-oxidase) were expressed especially in imbibed seeds. On the contrary, the expression of genes encoding GA 2-oxidase, ensuring the catabolism of bioactive GAs, decreased with the germination stage. These trends were similar in both genotypes. The gene encoding ABA biosynthetic enzymes tended to be expressed more in dry seeds (freshly harvested as well as after-ripened seeds) compared to imbibed ones. On the other hand, ABA 8′-hydroxylase (a key enzyme in ABA catabolism) genes were expressed especially in germinated seeds (Fig. S1).

### Transcriptional differences in seeds during post-harvest ripening and germination

To monitor and characterise ribosome association of mRNAs, we isolated mono- and polysomal fractions from freshly harvested (FH), after-ripened (3 months after harvest, AF), and 48-h imbibed (IM Col and Cvi seeds (Fig. 3a). In polysomal profiles of FH and AR seeds of both genotypes (Fig. 3b), monosomal fractions were detected, while polysomal peaks (referred to two or more ribosomes bound to mRNA) were not present. In these profiles, the fraction preceding monosomal peak was clearly visible. This peak/fraction was referred as a pre-monosomal peak (R), presumably containing lighter RNA-protein complexes such as 30S, 40S, 50S, 60S subunits of a eukaryotic pre-initiation complexes (43S a 48S.) Profile of 48h imbibed Col seeds contained both monosomal and polysomal peaks (Fig. 3c). On the contrary, in imbibed Cvi seeds, the polysomal peak was not detected (Fig. 3d). RNA extracted from individual fractionations was subjected to RNA-seq analysis (Table S1). The number of identified genes in Col samples ranged from 11,626 (Col-AR) to 12,087 (Col-FH), while in Cvi was slightly higher (13,419 in Cvi-FH, and 13,952 in Cvi-AR). Imbibed seeds monosomal fractions contained 19,674 (Col-IM) and 17,097 (Cvi-IM) genes (Fig. S2), respectively. PCA analysis showed the specific distribution of individual fractions (Fig. 4a). FH and AR samples clustered together, while Cvi and Col genotypes were clearly separated. Samples isolated from Col IM stages were distinct from FH and AR samples, indicating the germination switch. Cvi-IM samples were separated from the rest of the samples as well.

**Fig. 3:**
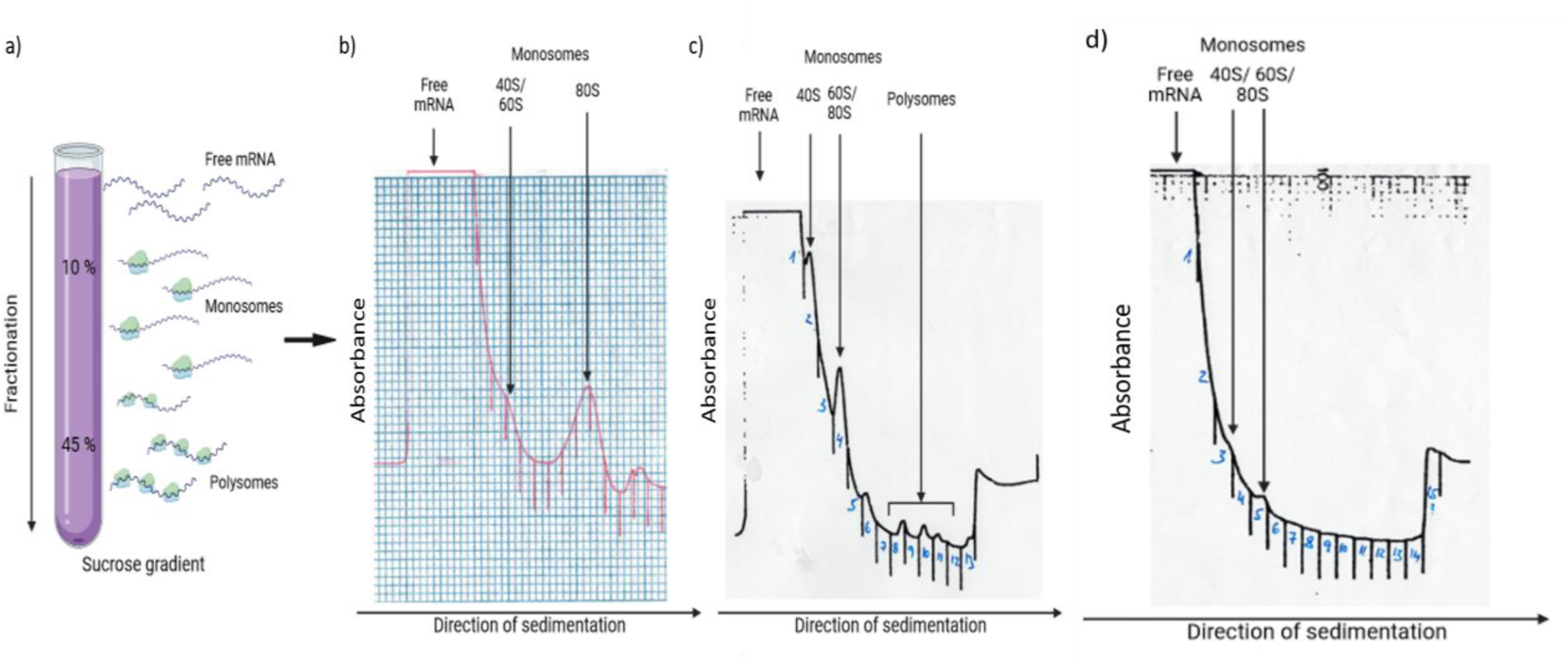
Sucrose gradient separation profiles. a) The scheme of mRNA and ribosomal distribution in a prepared sucrose gradient (10 a 45 % sucrose solutions) used for fractionation of total seed RNA/proteins. Polysome profiles of b) Col AR, c) Col IM, and d) Cvi IM samples. 40S/ 60S/ 80S-monosomal fractions. Marks visible on the absorbance charts represent border lines between collected fractions.

**Fig. 4:**
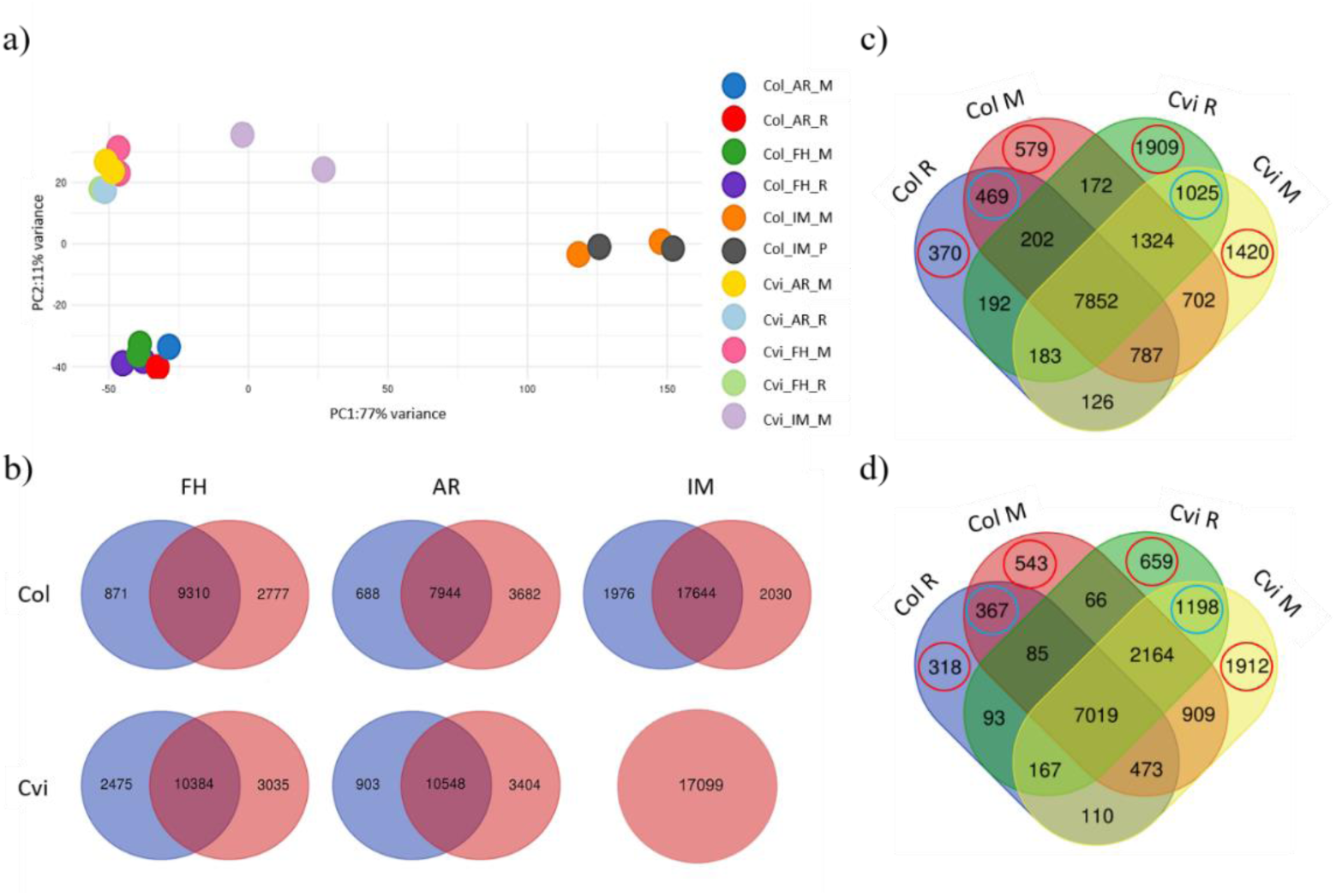
Comparison of polysome profiles between two different *Arabidopsis* genotypes. **a)** Principal component analysis (PCA) of RNA-seq samples isolated from *Arabidopsis* seeds of dormant (Cvi) and non-dormant (Col) genotypes in 3 different stages, FH-freshly harvested, AR-after-ripened, IM-48-h imbibition. **b)** Venn diagrams representing identified specific and shared genes from fractions isolated from Col and Cvi FH, AR and IM seeds. The upper line shows Col, and the bottom line shows Cvi fractions. In FH and AR seeds pre-monosomal (blue) and monosomal (red) fractions were compared. In IM seeds, monosomal (blue) and polysomal (red) fractions were analysed. **Comparison of transcripts from pre-monosomal and monosomal fractions of Col and Cvi genotypes** of certain developmental stages c) FH stage, d) AF stage. Red circles show genes specific for certain fractions, and blue circles genes shared between fractions of the specific genotype. R- premonosomal fraction, M- monosomal fraction.

First of all, we were interested in comparisons of respective fractions between two studied genotypes. In FH samples, there were 9,310 and 10,384 shared genes identified in Col and Cvi samples, respectively, while 2,777 and 3,035 were specific for respective fractions containing monosomes (Fig. 4b). Similarly, in AR samples, there were 7,944 and 10,548 shared and 3,682 and 3,404 monosome specific genes in Col and Cvi samples (Fig. 3). In pre-monosomal fraction (R) 688 to 903 genes were identified in Col and Cvi, respectively. Following RNA-seq analysis, we characterised specific and shared genes between different genotypes and stages (Fig. 4c,d). The highest number of identified genes was found in both fractions of Col IM seeds, composed of monosomes and polysomes. To investigate molecular functions, pathways and biological processes of identified genes, we performed enrichment analysis. Most genes were categorized as genes with Nucleic acid binding, Cation binding and Metal ion binding molecular function in case of both genotypes in all stages (Fig. S3a,b,c,d,e,f). In the IM stage of the Col and Cvi genotypes, GO term Transferase activity appeared (Fig. S3e,f). These results suggest a correlation between polysome-binding mRNA and transcription regulation. According to KEGG pathway enrichment analysis, most transcripts shared by fractions isolated from the particular genotypes in FH, AR, and IM stages referred to genes involved in Metabolic pathways, Biosynthesis of secondary metabolites, and Ribosome (Fig. S4a,b,c,d,e,f). Therefore, we can suggest that polysome-binding mRNAs contribute to metabolism and translation. Finally, GO term analysis revealed that fractions isolated from both genotypes in FH, AF, and IM stages shared between particular fractions genes conferring the following biological processes: Gene expression, Protein metabolic process, and Nucleobase-containing compound metabolic process (Fig. S5a,b,c,d,e,f). Altogether, polysome-binding mRNAs are probably involved in transcription and translation regulation, metabolism, and protein synthesis.

**Fig. 3:**
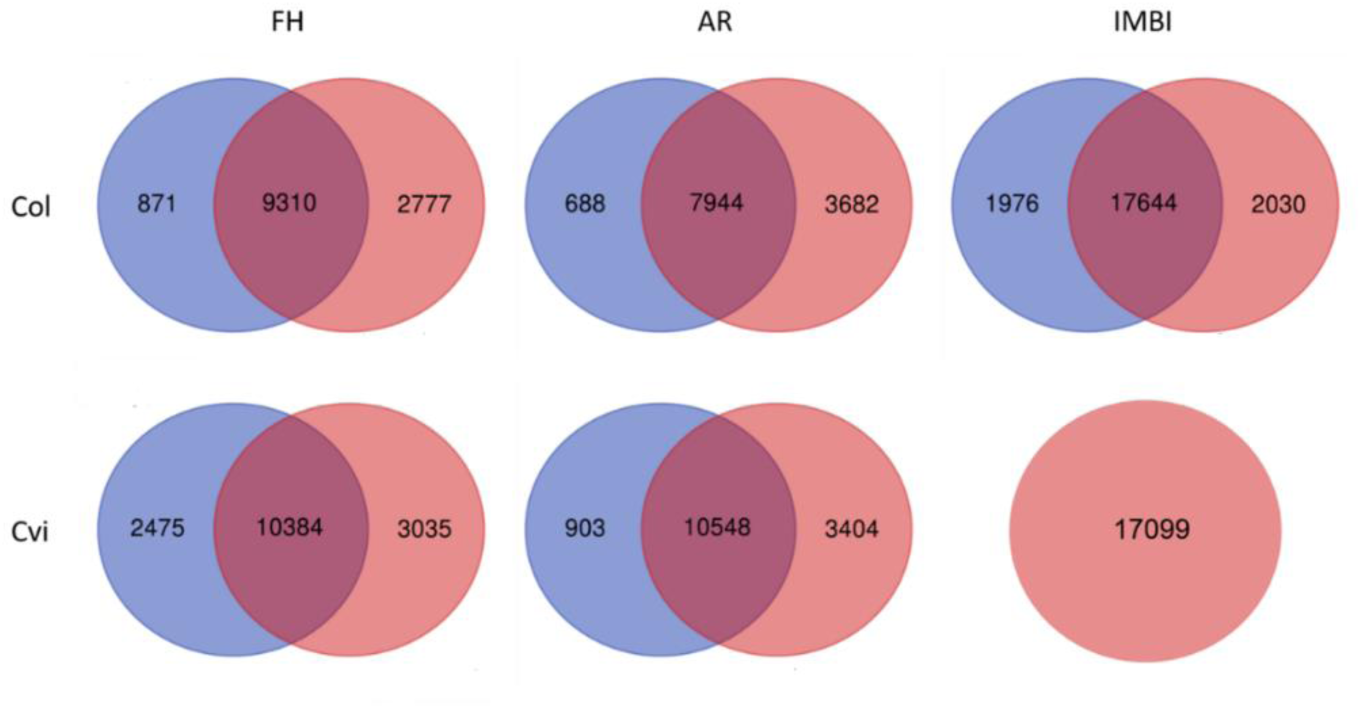
**Venn diagrams representing identified specific and shared genes** from fractions isolated from Col and Cvi FH, AR and IM seeds. The upper line shows Col, and the bottom line shows Cvi fractions. In FH and AR seeds pre-monosomal (blue) and monosomal (red) fractions were compared. In IM seeds, monosomal (blue) and polysomal (red) fractions were analysed.

To describe shared and unique genes for specific genotypes and stages, we compared pre-monosomal and monosomal fractions between Col and Cvi genotypes in FH (Fig. 4c) and AR stages (Fig. 4d). In FH seeds, a comparison of pre-monosomal and monosomal fractions of both genotypes showed that they shared 7852 genes (Fig. S6). Around 900 of them belong to the Metabolism pathway, 457 genes are involved in the Biosynthesis of secondary metabolites, 272 genes are categorized to the Ribosome pathway, 144 genes are part of the Carbon metabolism category, and 128 genes are categorized in Biosynthesis of the amino acid pathway. The molecular function of identified genes was generated as follows: 1572 genes-Nucleic acid binding, 1292 genes-Cation binding, 1283 genes-Metal ion binding, 892 genes-RNA binding, and 597 genes-mRNA binding. According to biological function, we identified groups of genes involved in the Protein metabolic process (1675 genes), the Gene expression (1667 genes), the Cellular protein metabolic process (1567 genes), the Response to abiotic stimulus (935 genes), and the Organonitrogen compound biosynthetic process (867 genes).

**Fig. 7:**
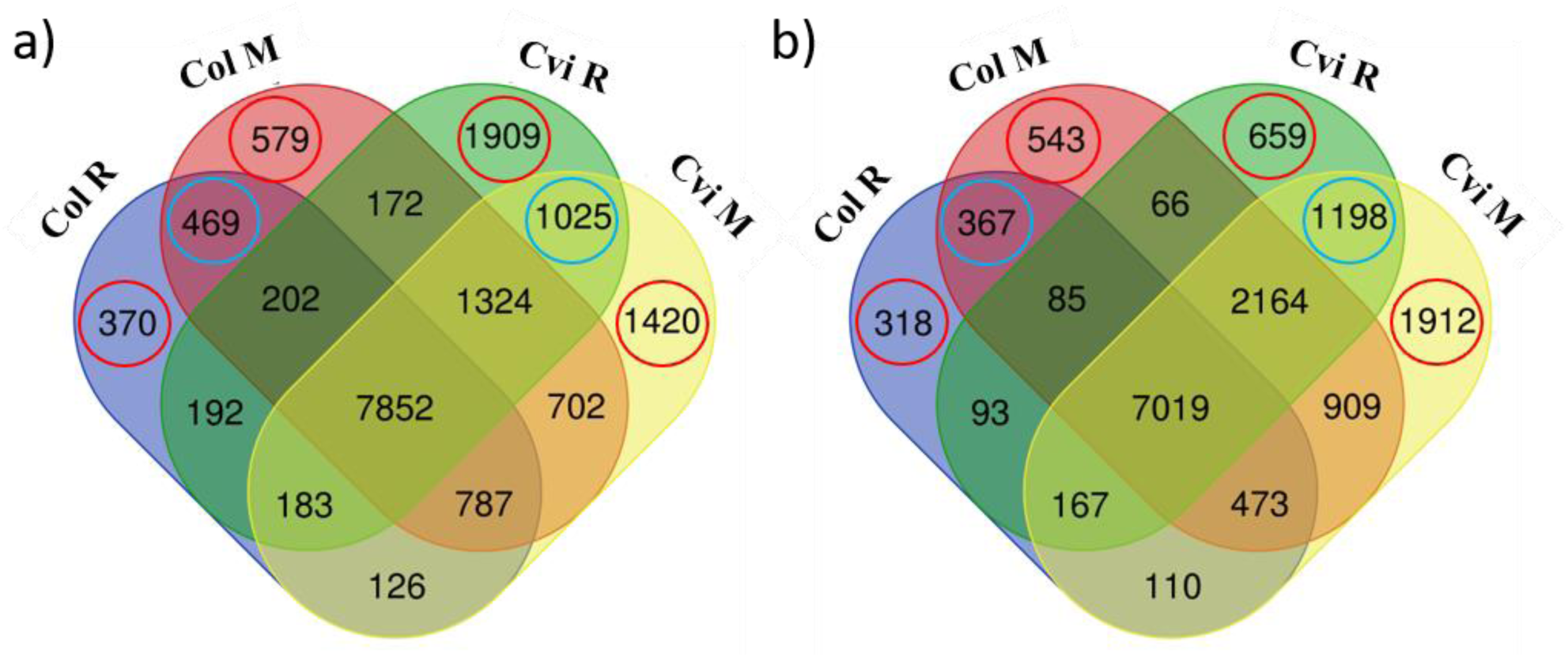
**Comparison of transcripts from pre-monosomal and monosomal fractions of Col and Cvi genotypes** of certain developmental stages. a) FH stage, b) AR stage. Red circles show genes specific for certain fractions, and blue circles genes shared between fractions of the specific genotype. Col R and Col M mean pre-monosomal and monosomal fractions, respectively, isolated from Col a) FH and b) AR seeds. Cvi R and Cvi M represent pre-monosomal and monosomal fractions, respectively, isolated from Cvi a) FH and b) AR seeds.

The 469 genes shared between pre-monosomal and monosomal fractions of Col FH seeds (Fig. S7). The genes found in the same fractions of Cvi seeds (1025 genes) were categorized as genes of Glycosylphosphatidylinositol (GPI)- anchor biosynthesis pathway (6 genes) (Fig. S8). The Nucleic acid binding (193 genes), the Transferase activity (186 genes), and the Cation binding (170) categories were identified in Molecular function category, with the highest number of genes included. GO terms in Biological process category showed the Nucleobase-containing compound metabolic process (216 genes), the Nucleic acid metabolic process (197 genes), and the Cellular protein metabolic process (189 genes) groups with the highest gene number. For the unique genes identified in the pre-monosomal fraction of FH Col seeds, no GO terms were generated because of inefficient enrichment. Unique genes in the monosomal fraction of the Col FH seeds were categorized into two GO terms within the Molecular function category: the RNA polymerase binding (5 genes) and the Starch binding (4 genes) (Fig. S9). The number of genes identified in the pre-monosomal fraction of the Cvi FHseeds was much higher than in the same fraction of Col (Fig. 4a, 1909 genes). For this group of genes, the Metabolic pathways (210 genes), the Carbon metabolism (39 genes), and the Spliceosome (24 genes) categories were generated according to the highest number of genes involved (Fig. S10). In the case of the Molecular function category, GO terms of the Nucleic acid binding (379 genes), the Cation binding (304 genes), and the Metal ion binding (301) showed a high abundance of identified genes. GO terms with the highest number of identified genes within the Biological process category were the Nucleobase-containing compound metabolic process (429 genes), the Gene expression (396 genes), and the Nucleic acid metabolism process (393 genes). KEGG pathway GO terms generated as unique for monosomal fractions of the Cvi FH seeds were the Homologous recombination (10 genes) and the Other glycan degradation (6 genes) groups (Fig. S11). Molecular function categories with the highest abundance of identified genes showed to be Small molecule binding (214 genes), Anion binding (209 genes), and Nucleotide-binding (207 genes). The highest number of genes from the fractions were categorized into the Nucleobase-containing compound metabolic process (279 genes), the Nucleic acid metabolism process (258 genes), and the Developmental process groups of Biological processes GO terms.

Afterwards, transcripts associated with individual fractions shared with both genotypes in the AR stage were investigated (Fig. 4d). The highest number of genes identified in KEGG groups of Metabolic pathways (766 genes), Biosynthesis of secondary metabolites (395 genes), and Ribosomes (272 genes) were shared with Col and Cvi genotypes in both types of fractions (7019 genes, Fig. 4d). In the case of the Molecular function category, GO terms of the Nucleic acid binding (1391 genes), the Cation binding (1120 genes), and the Metal ion binding (1113) showed a high abundance of identified genes. Go terms within the Biological process category showed Gene expression (1489 genes), Protein metabolic process (1457 genes), and Cellular protein metabolic process (1370 genes) groups with the highest gene number (Fig. S12).

The monosomal fractions of AR Col seeds contained genes with identified Molecular functions rRNA binding (11), Oxidoreductase activity acting on NAD(P)H quinone or similar compound as acceptor (6), NADH dehydrogenase (ubiquinone) activity (5) (Fig. S13). According to the Biological process, most genes were enriched in Gene expression (113), Macromolecule biosynthetic process (101), and Cellular amide metabolic process (38) categories. The pre-monosomal and monosomal fractions of Cvi AR seeds contained genes involved mostly in the mRNA surveillance pathway (14 genes) (Fig. S14). Moreover, the most frequent terms in the Molecular function category were Small molecule binding (171 genes), Nucleotide binding (161 genes), and Nucleoside phosphate binding (161 genes). The biological functions of genes within this genotype and stage were identified as Protein metabolic process (235 genes), Cellular protein metabolic process (218 genes), and Macromolecule modification (180 genes). For unique genes of pre-monosomal fractions of Cvi AR seeds, only biological functions were identified: Cellular protein modification process (94 genes), Protein modification process (94 genes), and Negative regulator of biological processes (50 genes) (Fig. S15). Monosomal fraction of Cvi AR seeds contained unique genes involved in Nucleic acid binding (352 genes), Transferase activity (350), and Cation binding (289 genes) molecular functions (Fig. S16). Moreover, the Nucleobase-containing compound metabolic process (391), Nucleic acid metabolic process (359), and Gene expression (327) are biological functions in which these genes were categorized mostly.

### RNA modifications change during the seed post-harvest ripening and germination

The RNA modifications were analyzed in dry and imbibed Col and Cvi seeds. The identity N-6-methyladenosine (m6A), N-1-methyladenosine (m1A), 5-methylcytidine (m5C) and 8-oxoguanosine (o8G) in RNA samples was unambiguously confirmed by UHPLC-MS analysis using chemical standards of selected modified nucleosides and their characteristic retention times and signals (m/z) are given in Table S2. The predominant modification was m1A (40x of m6A) both in Col and Cvi seeds. Higher content (up to 3x) of m5C was observed in Col, compared to Cvi seeds (Fig. 5). In addition to these modifications, 8-hydroxyguanosine was also detected, although at lower amounts in both seeds. Moreover, Col seeds have a larger (about 1.5-2x) amount of modified RNA, compared to Cvi (Fig. 5). There was a trend in the quantity of M6A RNA modification, with a clear peak at 3 months of post-harvest storage (Fig. 5) followed by a decline over 6 and 10 months of storage. A significant decrease was observed for other nucleosides that have been stored for more than 6 months apart from o8G amount in Col seeds, where storage longer than 6 months caused its increase. Changes in quantities of methyl and oxidized nucleoside derivatives confirm the assumption of gradual RNA degradation during seed ageing. When respective 3-month-old seeds were left to imbibe for 48h, this trend remained and the quantities of m6A, m1A and m5C dropped to half of the original amounts in both Col and Cvi seeds (Fig. 5).

**Fig. 5:**
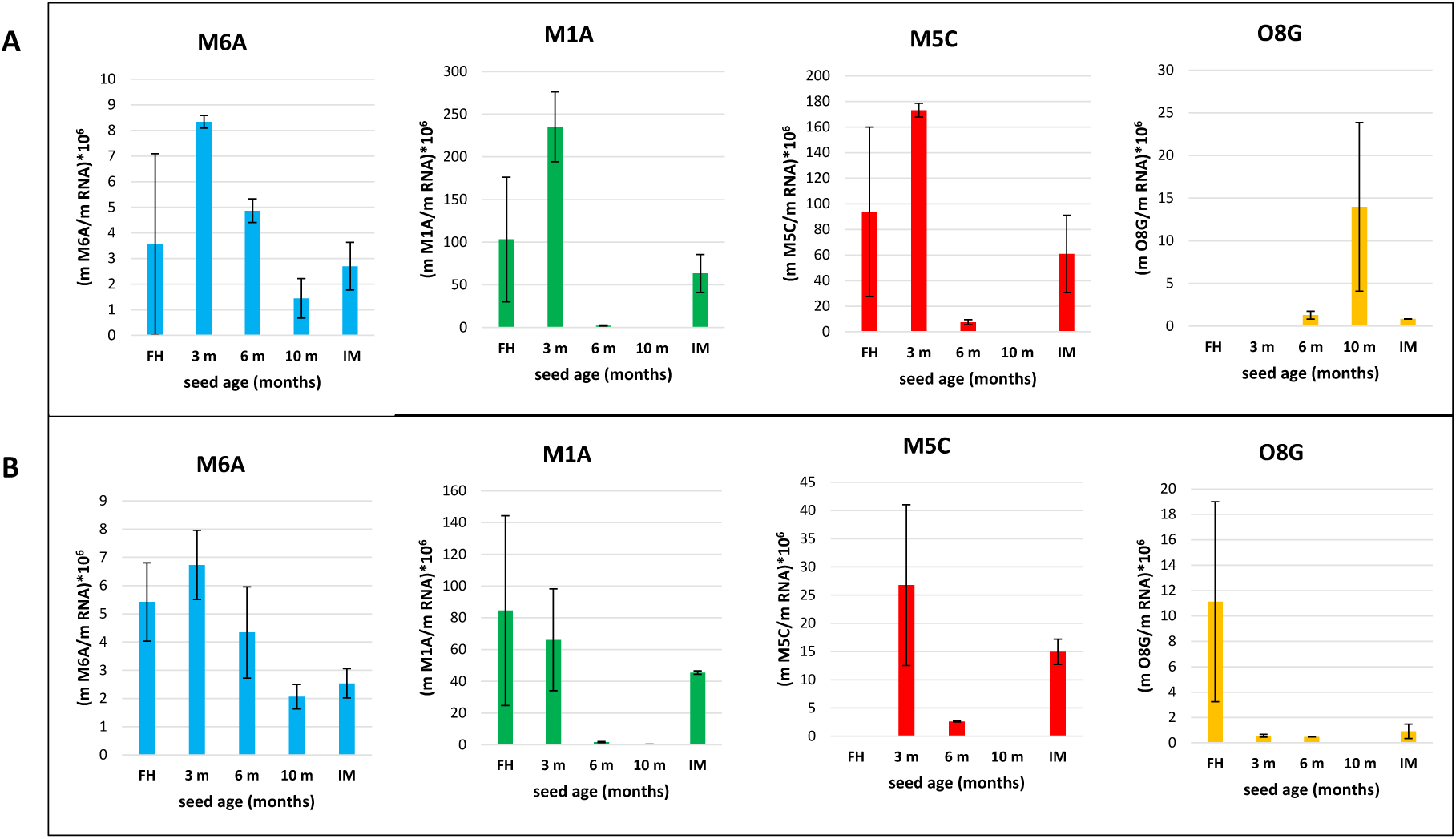
**Content of m6A, m1A, m5C and o8G RNA modifications in the A) Col and B) Cvi** after 3, 6 and 10-month ripening and 48-h imbibition of 3 months AR seeds. Expressed as a relation to total RNA.

Since the m6A is the most prevalent modification in eukaryotic mRNA affecting numerous aspects of mRNA metabolism (including translation), we used m6A sequencing to map this modification in dry (freshly harvested as well as after-ripened) and imbibed Col and Cvi seeds. We used an antibody-based approach, to enrich specifically for transcripts with m6A modifications. The obtained RNA fraction was then subjected to RNAseq analysis and subsequently bioinformatically processed. First of all, the m6A-modified genes were compared among the studied germination stages of each genotype separately (Fig. 6a). Afterwards, genes specific to the respective germination stage were compared between Col and Cvi seed fractions.

**Fig. 6:**
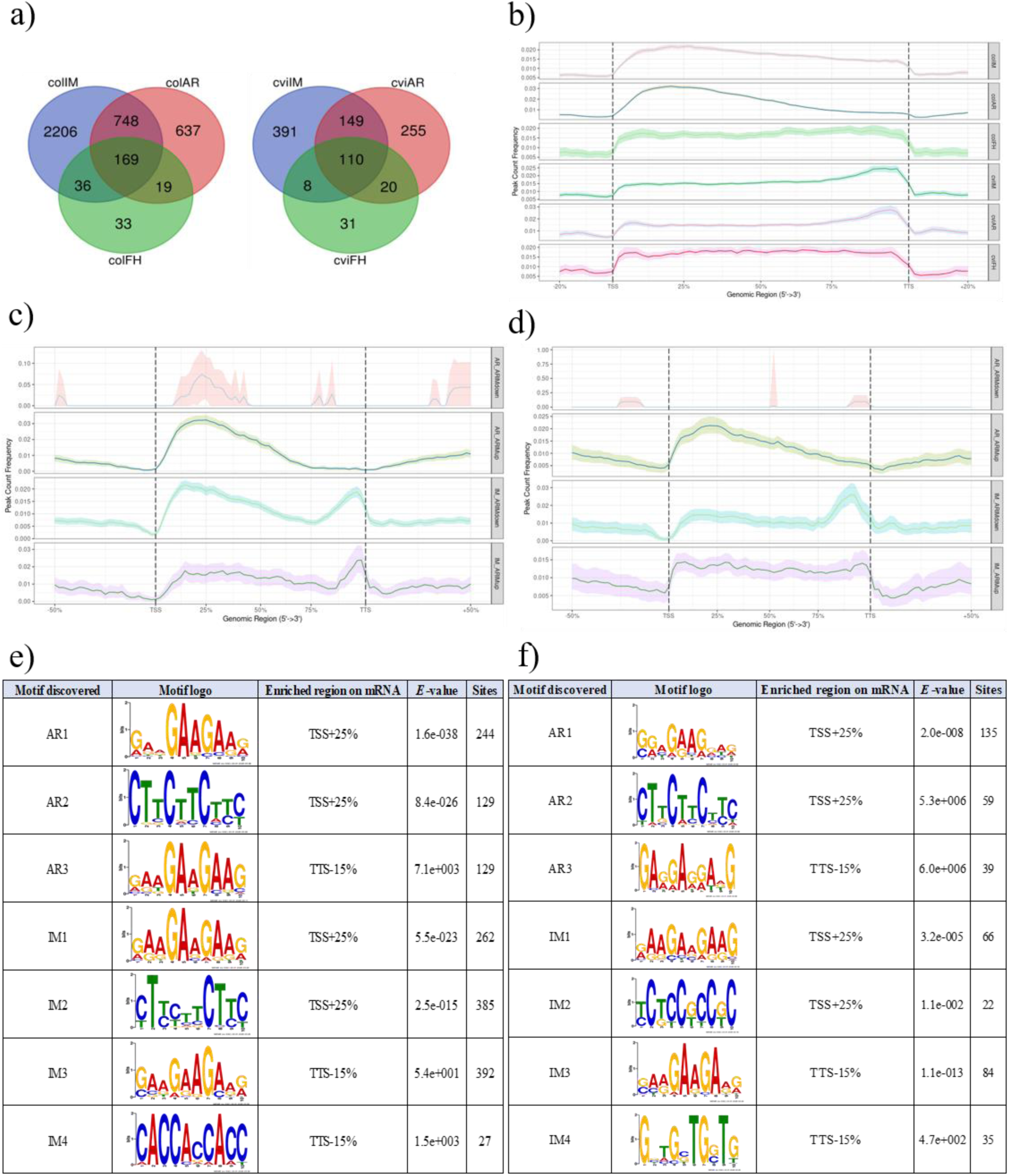
A characterization of the m6A methylome of Col and Cvi seeds. **a)-** The Venn diagrams illustrating shared and unique m6A-modified genes in dry (freshly harvested as well as after-ripened) and imbibed seeds of Col- (left) and Cvi (right). **b)-**The position of m6A modification in the sequences isolated from Col and Cvi seeds in freshly harvested, after ripening, and imbibed stages. **c), d)-** The position of m6A modification in the sequences of Col (left) and Cvi (right) m6A-modified genes. **e), f)-** Motifs identified in m6A-modified transcripts upregulated in Col (left) and Cvi (right) seeds at AR and IM stages. The enriched region was derived from the peak localization-TSS+25% - transcription start site + 25% of the sequence length forward, TTS-15% - transcription terminator site + 15% of the sequence length backward. Significant levels are indicated by the *E*-value. Motifs were searched in MEME-suite 5.5.7 software.

In Col samples, the most m6A-modified genes were detected in imbibed seeds, whereas the m6A-modified genes were the least abundant in dry freshly harvested seeds (FH). A similar trend was observed in Cvi seeds. Two genes were shared by Col and Cvi FH seeds (Fig. 1A). One of them belongs to the GRAS family transcription factor family protein. The second one is described as the transposable element gene. The genes unique to Col FH seeds can be characterized using the GO (Cellular component, Fig. 1B) terms Protein-containing complex and Cytosol, whereas those found in Cvi by GO terms (Fig. 1C) Autophagy, Fructose and mannose metabolism, Glycolysis/Gluconeogenesis, Catabolic process (Fig. 1D). The genes shared by Col and Cvi AR seeds (Fig. 2A) belong to GO term categories (Fig. 2B) Seed oilbody biogenesis and Galactolipid metabolism.

The genes specific to Col AR seeds were enriched in Go terms (Fig. 2C) Metabolic pathway, Biosynthesis of secondary metabolites and Response to various stimuli such as stress, chemicals, organics etc. (Fig. 2D). On the other hand, genes found only in Cvi AR seeds were characterized by GO terms (KEEG, Fig. 2E) Spliceosome, various Catabolic processes and terms associated with reproductive development (Fig. 2F). The genes shared (Fig. 3A) by Col and Cvi imbibed (IM) seeds belong to GO term categories (Fig. 3B) Ribosome, Spliceosome and Gene expression and Protein metabolic process (Fig. 3C). The genes specific to Col imbibed seeds were enriched in GO terms (Fig. 3D) Metabolic pathway, Biosynthesis of secondary metabolites and Cellular component organization and biogenesis and Localization (Fig. 3E). Imbibed Cvi seeds were characterized using the GO terms (Fig. 3F): Ribosome, Carbon metabolism, Spliceosome and Protein metabolic process, Gene expression (Fig. 3G)

### The position of m6A RNA modification varies between Col and Cvi genotypes

We localized the position of m6A modification in m6A-modified genes and found that there is a significant difference between the Col and Cvi genotypes (Fig. 6b). Since the genotypes differing in the level of their seed dormancy were used for analysis, we focused on differences between transcriptionally-inactive, after-ripened (AR), and transcriptionally-active, imbibed (IM), seeds. While AR and IM Col samples showed the highest m6A peak near the start codon, the respective stages of Cvi contained transcripts with m6A modification near the stop codon in the coding sequence (Fig. 6b). To describe a connection between a m6A-peak position and gene expression, we analysed m6A-modified up- and downregulated genes between AR and IM seeds of each genotype (Fig. 6c,d). In AR and IM Col samples, the prevalence of m6A peaks was measured around the transcription start site (TSS) in 25 % of the length of the sequence from the TSS point (TSS+25) (Fig. 6c). On the contrary, AR and IM Cvi samples had the highest number of peaks at the end of the coding sequence (CDS), around 15 % of the length prior to the transcription terminal site (TTS-15). Interestingly, m6A peaks in both genotypes, isolated and upregulated at the AR stage, were detected in TSS+25 only (Fig. 6c,d). The motif analysis of detected peaks at the TSS+25 site within transcripts upregulated in the AR Col seeds (446) indicated that GAAGAAGAAG and CTTCTTCTTC motifs were presented in 244 and 129 generated sequences, respectively (Fig. 6e). At TTS-15, the GAAGAAGAAG and TCTTCTTC motifs were identified in 129 and 40 sequences, respectively. The involvement of these genes in biological processes according to GO terms (Fig. S20-22) was investigated. Genes with these specific motifs at the TSS+25 site were suggested to be involved in Response to stress, Cellular nitrogen compound biosynthetic process, and Response to chemical stimulus. Moreover, they belong to the following molecular function categories: Metal ion binding, Cation binding, and Hydrolase activity. Interestingly, genes with GAAGAAGAAG motif at the TTS-15 site were suggested to be involved in the Seed dormancy, Dormancy, and the Autophagy biological processes (Fig. S22). From the molecular function point of view, the categories were the same as in the first case. On the other hand, m6A peaks found and upregulated in the IM Col seeds were modified in two sites within CDS (Fig. 6c). In both sites, TSS+25 and TTS-15, the motif GAAGAAGAAG was identified (Fig. 6e). From the identified 902 transcripts (genes), 262 contained this specific motif in the TSS+25 site and 394 in the TTS-15 site. The second motif identified in 385 sequences of the TSS+25 site was the CTCCTTCTTC. Additionally, another motif, CACCACCACC, was found in 27 sequences at the TTS-15. GO terms for genes containing GAAGAAGAAG motif at both analyzed sites were very similar (Fig. S23,S24). The most enriched biological functions for these genes were the Protein metabolic process, the Cellular protein metabolic process, and the Gene expression. Similarly, in the case of the molecular functions, the Nucleic acid binding and RNA binding functions were the prevalent categories. Moreover, the highest number of genes from both groups were suggested to be involved in the Ribosome pathway. Sequences containing the CTCCTTCTTC motif belongs into Metabolic process and Biosynthesis of secondary metabolites categories of the KEGG pathway (Fig. S25). Small molecules binding and Nucleotide binding molecular functions were enriched in this group of genes. Finally, these genes were supposed to be involved in Protein metabolic process. Genes with the CACCACCACC motif at the TTS-15 site were shown to be involved in the Cell wall organization process, The external encapsulating structure organization, and the Cell wall organization or biogenesis processes (Fig. S26). Furthermore, the GO terms the Structural molecule activity, The structural constituent of the cell wall, and the Demethylase activities were generated for this particular group of genes. Finally, these genes were supposed to be involved in Oxidative phosphorylation and the Endocytosis KEGG pathways.

In Cvi seeds, transcripts with m6A modification upregulated at the AR stage contained m6A peak at the TSS+25 site (Fig. 6d). Furthermore, the analysis of motifs of respective genes (135 sequences) indicated that GGAGAAGGAG and CTTCTTCTTC motifs were presented in 135 and 59 generated sequences, respectively (Fig. 6f). At the TTS-15 site, the GAGGAGGAAG motif was identified in 39 sequences. Genes with motifs at the TSS+25 site were categorized into Oxidoreductase activity and Ubiquitin-like protein transferase activity groups (Fig. S27, 28). Moreover, they belong to the following biological processes: Response to stress, Response to abiotic stimulus, and Catabolic process. Genes with the motif identified at the TTS-15 site referred to biological processes as Response to abiotic stimulus and Proteolysis (Fig. S29). The m6A-modified transcripts upregulated in the IM Cvi seeds were modified mainly at the TTS-15, and with the lower frequency also at the TSS+25 site (Fig. 6f). Again, in both sites, TSS+25 and TTS-15, the motif GAAGAAGAAG was found. From 171 transcripts, 66 contained this specific motif in the TSS+25 site and 86 in the TTS-15 site. The second motif identified in 22 sequences of the TSS+25 site was the TCTCCGCCGC. The second motif, GGTGCTGGTG, was also found in 35 sequences at the TTS-15. These motifs were identified in genes GO categorized as gene expression, translation and ribosome pathways (Fig. S30-33). IM Col seeds had a higher number of transcripts with m6A modification at the beginning of CDS, upregulated in the IM stage (Fig. 6c), compared with those of IM Cvi seeds (Fig. 6d). This suggests a positive regulation of transcription by m6A modification at this particular position.

### Proteome composition of mono- and polysomal fractions show differences in translation machinery between Col and Cvi

To investigate the protein composition among different seed stages and fractions, the proteins from monosome and polysome fractions were extracted and analysed by LC-MS/MS. The protein group intensities ratios were used to identify the differences between mono- and polysomal fractions. Altogether, 14488 proteins were identified. In Col, monosomes contained 5051, 5303, and 5329 proteins in freshly harvested, after-ripened, and imbibed seeds, respectively. For Cvi, monosomes had 4437 proteins in freshly harvested seeds, 5,398 in after-ripened seeds, and 4,251 in imbibed seeds. The comparison of respective monosomal and polysomal fractions between Col and Cvi genotypes at FH and AR, FH and IM, AR and IM stages, revealed similarities in the case of major translation-related proteins, but also some qualitative and quantitative differences (Fig. 7). The differences were particularly in the case of monosomes and polysomes at FH and AR stages, where in Col there were 1139 and 2052 shared between monosomes and polysomes, while 3827 and only 67 were in Cvi samples. Comparisons of freshly harvested and AR with imbibed showed a higher number of shared in the case of Col (1169 and 3088 compared to 78 and 55 in Cvi) which corresponds to seed germination and higher translational activity.

**Fig. 7:**
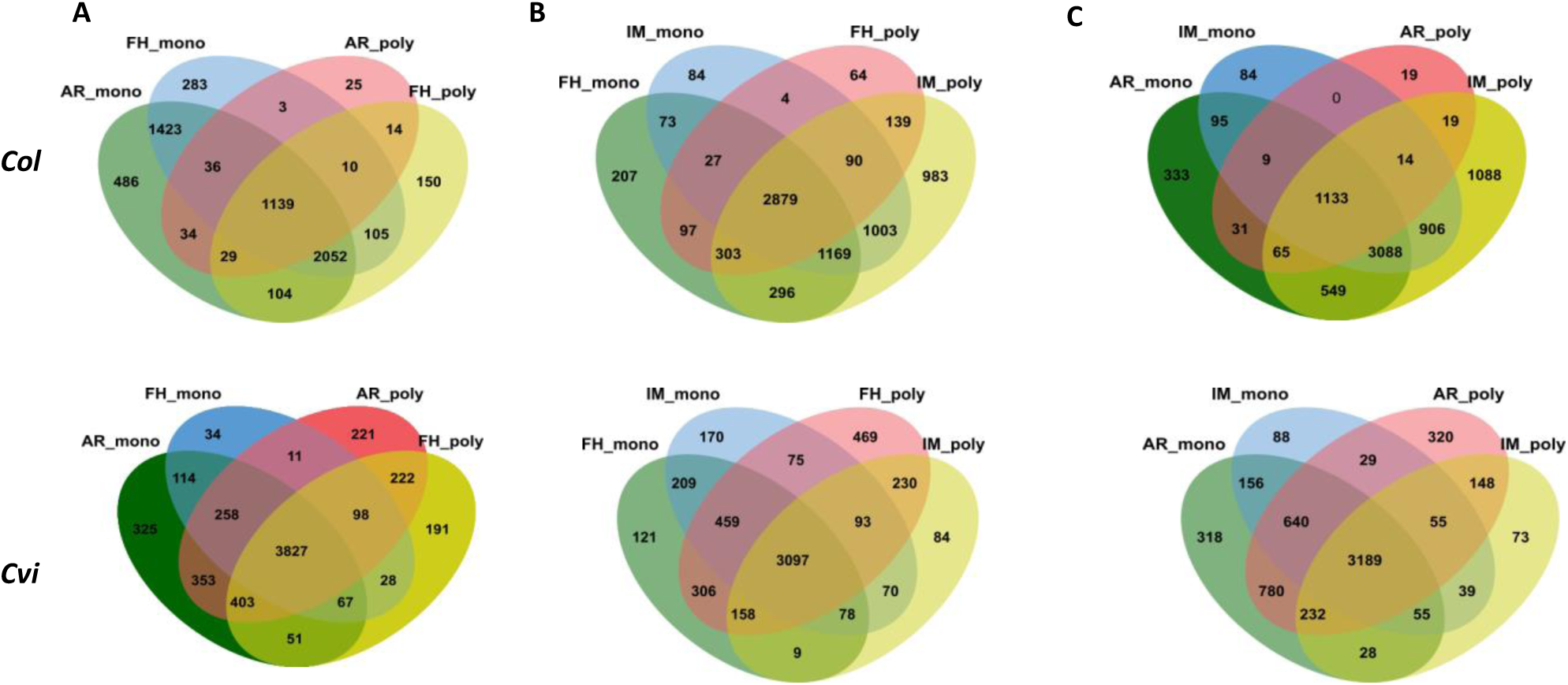
**The Venn diagram of proteins comparison between monosome and polysome fractions** in Columbia (Col) Cape Verde Islands (Cvi) seeds at FH, AR and IMB stages.

In the case of RNA-binding proteins (Zhang *et al*. 2023) there was a comparable number of proteins at FH and AR stages, while there was a difference at IM stage (1877 in Col and 1544 proteins in Cvi). There were Col-specific proteins in 40S, procession bodies and PABP/ALBA/ECT families (Fig 8,9). In Col monosomes enrichment analysis showed GO terms such as organic acid binding, carboxylic acid binding, and oxidoreductase activity for FH and AR seeds, while proteins of IM seeds were characterized by RNA helicase activity and ATP-dependent activity acting on RNA. The main KEGG pathways across all stages included aminoacyl-tRNA biosynthesis, carbon fixation, and the TCA cycle. Polysomes of Col contained 3603 proteins in FH, 1290 in AR, and 6862 in IM seeds. These were enriched in GO terms such as ATP-dependent protein folding chaperone and oxidoreductase activity across stages, with RNA helicase activity being prominent in imbibed seeds. KEGG pathways consistently included carbon fixation, glyoxylate and dicarboxylate metabolism, and the TCA cycle.

**Fig. 8:**
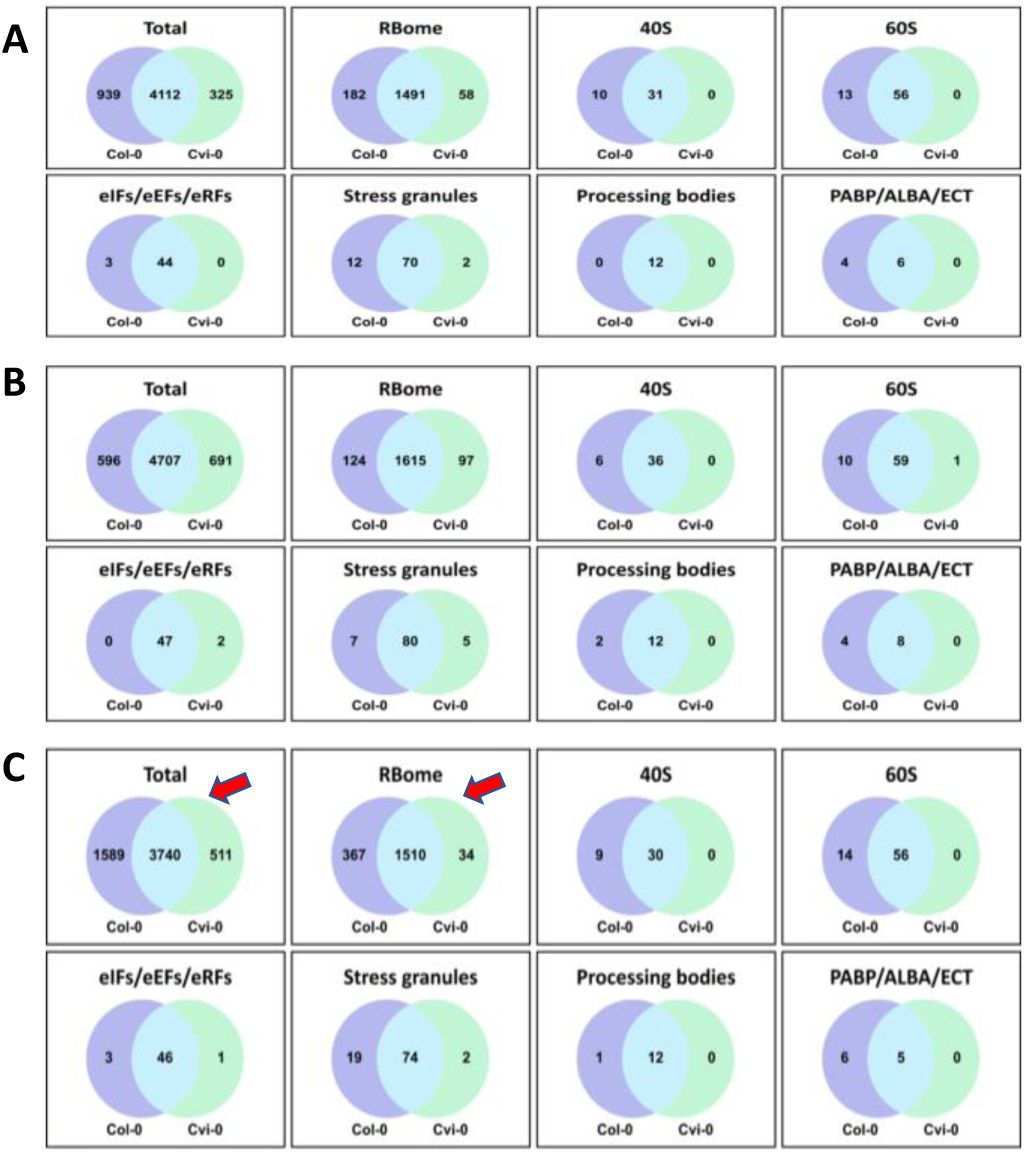
Comparative analysis of monosome fractions comparing Columbia (Col) and Cape Verde Islands (Cvi) at FH (A), AR (B) and IM (C) stages. Red arrows point to the significant differences between Col and Cvi.

**Fig. 9:**
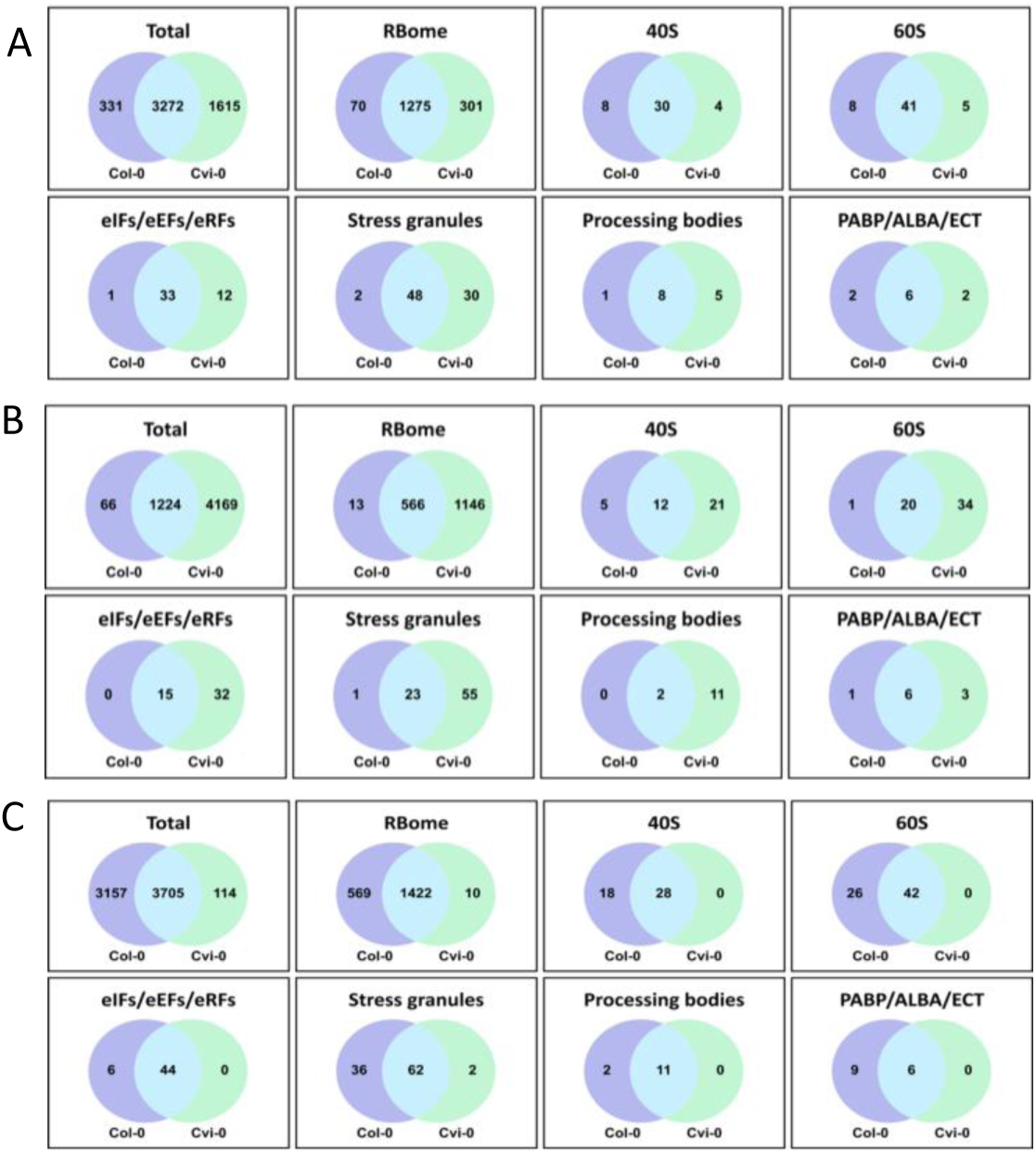
Comparative analysis of polysomal fractions comparing Columbia (Col) and Cape Verde Islands (Cvi) at FH (A), AR (B) and IM (C) stages. Protein groups were classified into total proteins, RNA-binding, 40S and 60S ribosomal subunits, eIFs/eEFs/eRFs, stress granules, processing bodies, and PABP/ALBA/ECT proteins.

For Cvi monosomes GO terms included ATP-dependent protein folding chaperone, NAD binding, and oxidoreductase activity. KEGG categories such as carbon fixation, aminoacyl-tRNA biosynthesis, and glyoxylate and dicarboxylate metabolism were dominant across all stages. Polysomes of Cvi contained 4887 proteins in FH, 5393 in AR, and 3819 in IM seeds, with GO enrichment for ATP-dependent protein folding chaperone, NAD binding, and oxidoreductase activity across all stages. KEGG categories included carbon fixation, glyoxylate and dicarboxylate metabolism, the TCA cycle, and the pentose phosphate pathway. Our data revealed differences in 40S and 60S ribosomal proteins between monosomes and polysomes, with additional detection of several 30S and 50S ribosomal proteins. RNA-binding proteins (RBPs) regulators of mRNA metabolism, including stability and translational activation of long-lived mRNAs. Notably, the dataset included RBP families such as ALBA, PUM, eIF, RAMP4, RRM/RBD/RNP, RGG, and glycine-rich RNA-binding proteins, with PUM and RRM/RBD/RNP proteins being the most abundant. There were similarities in GO molecular function assignments between respective Col and Cvi fractions, but also differences in the number of RNA binding proteins at AR and IM stages (Table 1). In the case of monomes of Col at IM stage there were 1436 proteins with Go term organic cyclic compound binding (GO:0097159), absent in Cvi. In addition, Col polysomes contained substantially higher (1344) RNA binding proteins, compared to only 386 in Cvi fraction (Table 1) likely related to differences in germination. This analysis demonstrated that both fractions in FH and AR seeds exhibit strong ribosomal and RNA-binding functions, with additional folding and stress-related activities in the monosome fraction and energy-transducing processes in the polysome fraction. The respective monosome fractions showed additional functional diversity, encompassing RNA binding, ion binding, and redox-related activities. In case of imbibed seeds (IM) the results demonstrate the functional focus on ribosomal, RNA-binding, protein-folding activities, and metabolic and stress-related regulatory processes. These functions reflect the translational activation and regulatory adjustments occurring in the imbibed stage.

**Table 1:**
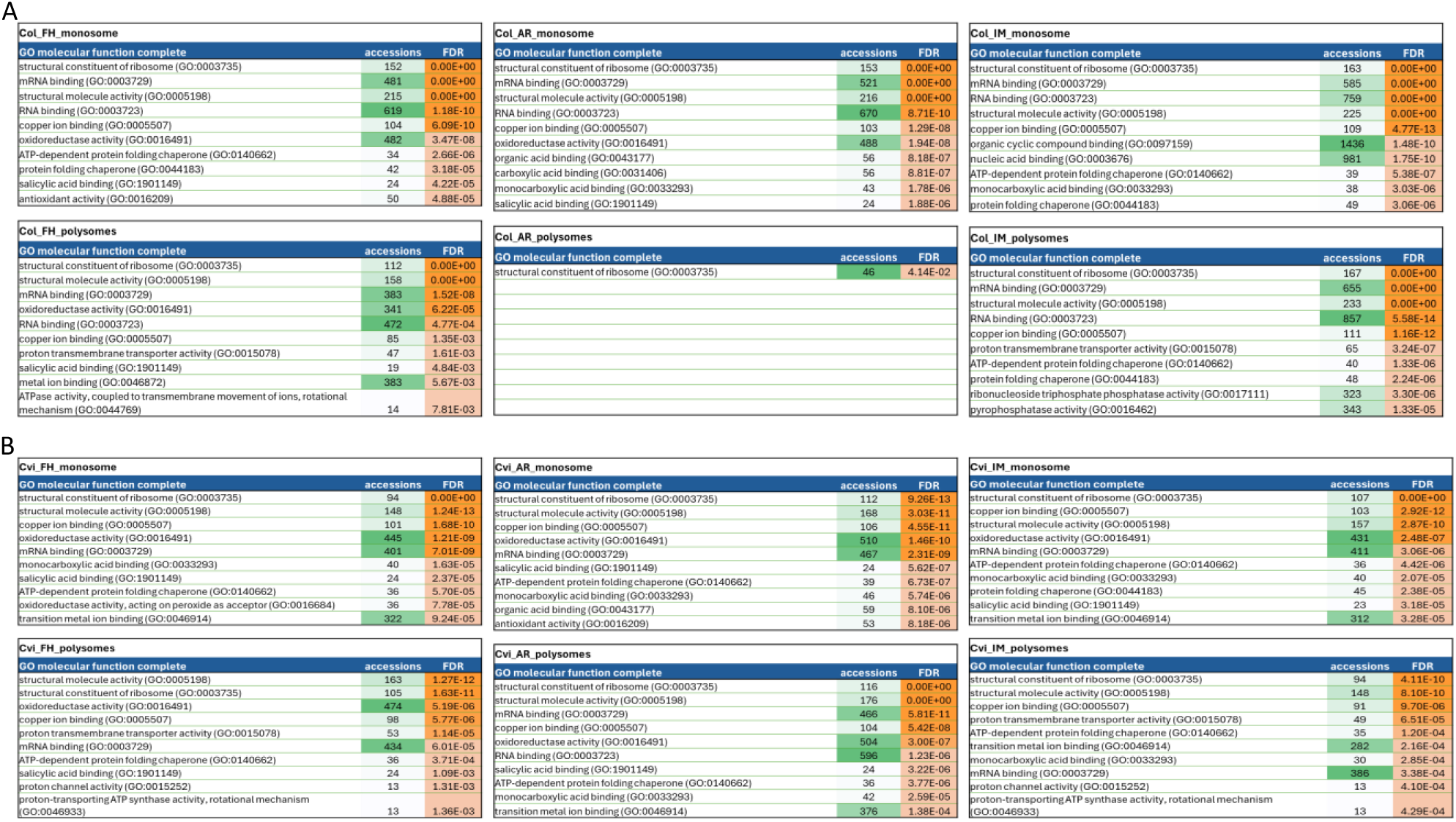
List of GO molecular functions of proteins isolated from monosomal and polysomal fractions of Col (A) and Cvi (B) at FH, AR and IM stages.

In the Total protein group, 4707 proteins were shared between Col and Cvi, with 596 unique to Col and 691 were specific to Cvi. Within the RBome (based on the dataset of Zhang *et al*. (2023), 1615 proteins are shared, while 124 are unique to Col and 97 were specific to Cvi. The 40S ribosomal subunit contained 36 shared proteins, with 6 unique to Col and none specific to Cvi. For the 60S ribosomal subunit, 59 proteins were shared, with 10 unique to Col and 1 specific to Cvi (Fig. 8,9).

The analysis of Col and Cvi seed fractions at AR stage showed differences in protein distributions across monosome and polysome fractions in various functional groups. In the Total protein group, 4065 or 4841 proteins were shared between monosome and polysome fractions in Col or Cvi, respectively, with 1238 or 557, and 52 or 552 proteins uniquely associated with monosomes and polysomes, respectively (Fig. 8,9). Within the RBome, 1168 proteins were shared, while 571 are specific to monosomes and 8 are specific to polysomes in Col, while in Cvi 1629 proteins were shared and 83 were unique to each fraction. Analysis of ribosomal subunits indicates that for the 40S ribosomal subunit, 29 or 33 proteins were shared, with 13 or 7 unique to monosomes and 4 or 1 to polysomes in Col or Cvi, respectively. Similarly, for the 60S ribosomal subunit, 48 or 53 proteins were shared, with none uniquely found in polysomes in Col, while 7 unique to monosome and 1 specific to polysome in Cvi. In the group of eIFs/eEFs/eRFs, 32 proteins are shared between fractions, with no unique proteins in either fraction. For Stress Granules, 63 proteins are shared, with 24 unique to monosomes and none specific to polysomes. The Processing Bodies contain 12 shared proteins, with 0 and 2 unique proteins in monosome and polysome fractions, respectively. Finally, in the PABP/ALBA/ECT category, 11 proteins are shared between fractions, with none uniquely present in either monosomes or polysomes.

In case of imbibed seeds, the polysome fraction (Col-IM), several proteins associated with active translation and enzymatic processes were upregulated, including small ribosomal subunit proteins such as RPS14C, RPS3AA, and RPS27D, metabolic enzymes such as asparagine synthetase 2 (ASN2), indole glucosinolate O-methyltransferase 4 (IGMT4), and NADPH-cytochrome P450 reductase 1 (ATR1) and proteins involved in cell division and regulation, such as cell division-related protein-like (At5g60110). In the monosome fraction (Col-IM), a wide range of ribosomal proteins were upregulated, including large subunit proteins such as RPL13D, RPL22B, and RPL36C, and small subunit proteins such as RPS10A and RPS11A. Translation initiation factors, such as EIF2B2, EIF4B1, and EIF6-2, were also prominent, along with regulatory proteins such as the mRNA-decapping enzyme subunit 2 (DCP2) and polyadenylate-binding protein RBP45C. Additional proteins like thioredoxin H5 (TRX5) and V-type proton ATPase subunit G1 (VHA G1) were present, reflecting broader regulatory and stress-related functions. Altogether, this analysis highlights distinct functional roles of polysome and monosome fractions in Col IM, showcasing proteins associated with active translation in polysomes and structural or preparatory roles in monosomes during seed imbibition. To identify the differences between mono- and polysomal fractions, the protein intensities ratios were used. These ratios were calculated, using medians of the given sample types. As a result of this analysis, 430 and 589 proteins were identified to be significantly up-regulated in polysomes as compared to monosomes in Col and Cvi, respectively. There were 28 DNA-binding, transcription factors, while in case of down-regulated 127 were of ribosomal proteins as expected. Among upregulated proteins in polysome fraction of both Col-FH and Col-AR were, glycolate oxidase 2 (GLO2), a protein involved in oxidative metabolism; eukaryotic translation initiation factor 3 subunit E (TIF3E1), a critical component of translation initiation; along with metabolic enzymes like asparagine synthetase 2 (ASN2) and glycolate oxidase 2 (GLO2). In the monosome fraction of Col-AR, numerous ribosomal proteins were upregulated, reflecting their presence in translationally inactive complexes. These include small ribosomal subunit proteins such as RACK1A (RACK1z) and RPS12C, large ribosomal subunit proteins like RPL13B, RPL8A, and RPL30W. Additional upregulated proteins include translation-related factors such as the 60S ribosomal export protein NMD3 and translationally controlled tumor protein 1 (TCTP1).

The monosome fraction of dry seeds contained unique ribosomal proteins including eL36x (RPL36C) or eL18y (RPL18B) from the large subunit, along with small subunit proteins uS9y (RPS16B) or eS8y (RPS15AC). Regulatory proteins such as ECT3 (YTH DOMAIN-CONTAINING FAMILY PROTEIN 3) and ECT4 were uniquely present, indicating their role in m6A-mediated RNA regulation in seeds contributing to maintaining mRNA stability and controlling translation. This corresponds well with differences in m6A RNA modifications. In dry seed polysomes, unique proteins formed a large group of all studied groups of proteins, including RPs, ALBA family proteins or eIF3 complex subunits. In imbibed seed monosomes, unique proteins included ribosomal proteins uS7y (RPS5A) and eS28x (RPS28C). Translation-related factors like IF3-2 (TRANSLATION INITIATION FACTOR IF3-2, CHLOROPLASTIC) and RNA-binding proteins such as ECT2 (YTH DOMAIN-CONTAINING FAMILY PROTEIN ECT2) were also identified (Fig. 10). RNA-binding proteins in polysomes from imbibed seeds included ECT2 and ECT6 (YTH DOMAIN-CONTAINING FAMILY PROTEINS), alongside translational machinery components such as EIF4E1 (EUKARYOTIC TRANSLATION INITIATION FACTOR 4E-1) and mitochondrial IF3A-1 (TRANSLATION INITIATION FACTOR 3 SUBUNIT A). Ribosomal proteins unique to seeds included eL18y (RPL18B), eS19y (RPS19C), and uL22y (RPL22Y).

**Fig. 10:**
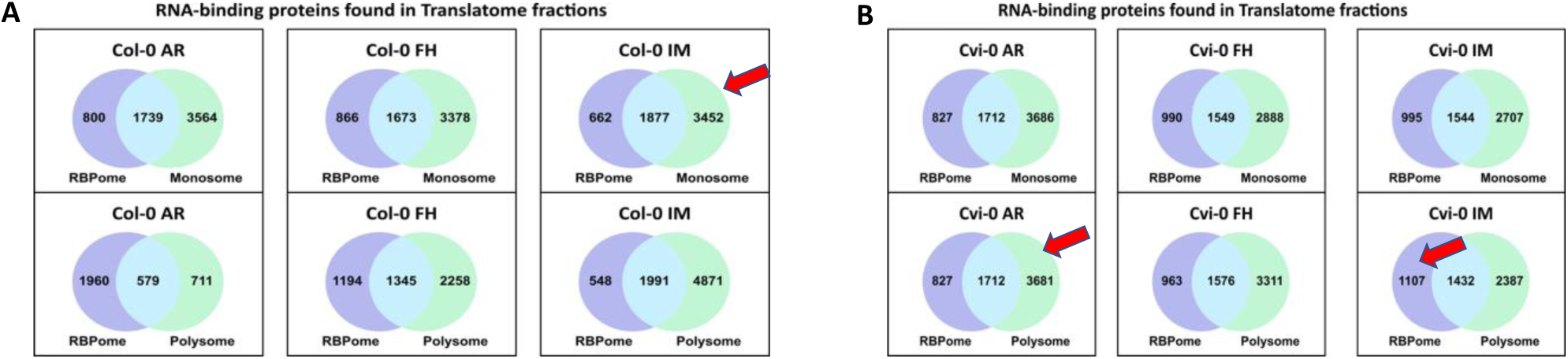
**Venn diagrams of RNA-binding proteins identified in the seed RBPome** (based on the dataset of Zhang *et al*. (2023) and monosome and polysome fractions at AR, FH and IM stages of Col and Cvi seeds. Red arrows point to the significant differences between Col and Cvi.

## Discussion

Seed dormancy is a critical trait shaped by both natural selection and agricultural practices. In agriculture, a delicate balance is required to prevent premature germination while avoiding excessive dormancy, as both extremes negatively impact crop yield and seed utilization (Smýkal *et al*. 2018). Germination is pivotal for seedling establishment and subsequent plant generations. This process relies on the longevity of seed-stored mRNAs, which are translated during germination. The separation of transcription and translation during seed maturation and germination makes seeds (and pollen) unique systems to study developmentally regulated translational switches (Sanjeev *et al*. 2019). In dormant stages, such as seeds and pollen, gene expression is predominantly regulated post-transcriptionally. Processes like mRNA sequestration into ribonuclear complexes and translational regulation play key roles (Hafidh and Honys, 2021). The activation of stored mRNA translation during germination begins with the coupling of the 40S ribosomal subunit to the 60S subunit to form an active 80S ribosome, initiating the elongation phase (Bailey-Serres *et al*. 2009). Translational control has been extensively studied in pollen tube growth (Hafidh *et al*. 2018; Sze *et al*. 2024) and seed germination (Basbouss-Serhal *et al*. 2015; Layat *et al*. 2014). These studies revealed significant differences in the translatomes of dormant and non-dormant seeds, driven by selective recruitment of mRNAs to polysomes. This suggests that long-lived mRNAs synthesized during seed development are stored and translated only upon imbibition (Kimura and Nambara, 2010).

Our study compared two *Arabidopsis* accessions with contrasting dormancy phenotypes: non-dormant Col and highly dormant Cvi. We analyzed polysomal and monosomal mRNA loading at three key stages: mature dry seeds (FH), after-ripened (AR) seeds, and seeds 48 hours post-imbibition (IM). This work provides a comprehensive RNA-seq-based analysis, extending beyond previous microarray-based studies (Basbouss-Serhal *et al*. 2015; Layat *et al*. 2014; Bai *et al*. 2017, 2020). Unlike earlier studies that focused on a single genotype, our comparison of contrasting dormancy phenotypes highlights the functional differences in translational regulation. In agreement with previous findings (Cadman *et al*. 2006; Finch-Savage *et al*. 2007), Col seeds exhibited a germination rate of ∼30 % immediately after shedding, while Cvi seeds required an after-ripening period of ∼100 days to achieve 85 % germination. We selected a 48-h imbibition period for analysis, a stage when Col seeds had fully germinated, with root emergence and cotyledon exposure, marking a developmental transition analogous to pollen tube growth (Hafidh *et al*. 2024). This stage aligns with the germination translational shift reported by Bai *et al*. (2017), coinciding with the full activation of the seedling developmental program. We expected that the different germination rates of Col and Cvi seeds could be explained by differential contents of ABA and GAs in the seed. Surprisingly, there were no substantial difference between Col and Cvi seeds in the levels of ABA and its metabolites (Fig. 1) indicating that the low Cvi germination rate could not be explained by higher ABA concentrations in its seeds. Carrera *et al*. (2008) proposed that ABA does not affect after-ripening, which showed also our data. On contrary, among m6A-modified genes specific for AR Col seeds were found the DOG1-like 4 (AT4G18650) and RDO5 (AT4G11040) genes, both involved in the regulation of level of seed dormancy. These genes were not found among m6A-modified genes of Cvi seeds. Whether the m6A-modification have inhibitory or promoting effect on gene expression need further research. Nevertheless, it was proposed that seed dormancy of imbibed seeds is mainly governed at the transcriptional level (Bai *et al*. 2018). Previously, GA 3-oxidases, the enzyme producing the bioactive GAs, were shown to play a role in seed germination in Cvi seeds (Cadman *et al*. 2006, Yazdanpanah *et al*. 2017). Thus, we analysed GA levels in Col and Cvi seeds as well. Our data indicated that GAs of the 13-hydroxylated pathway were dominant in germinating Col and Cvi seeds. Beside GA_1_, GA_4_, the bioactive GA of the 13-non-hydroxylated pathway, was one of the most abundant bioactive GAs in both genotypes, although the GAs of these pathway were not detected in our samples (except GA_34_) (Fig. 2). Similarly, GA_4_ was the prevalent bioactive GA also in germinating Landsberg erecta (Ler) seeds. However, the 13-non-hydroxylated GAs were dominant in Ler germinating seeds (Ogawa *et al*. 2003). Non-dormant Col and dormant Cvi accessions were compared at the mature dry seed stage using total RNA, comparing the effect of cycloheximide preventing translation (Kimura and Nambara 2010). This study found no large differences in transcriptomes of these two accessions, concluding that stored mRNAs do not reflect the degree of dormancy but a developmental stage. However, the drawback of this study is based on total RNA used as a proxy for being used for translation during germination process.

We have identified that over 10,000 genes are expressed and present as RNA in the mature seeds. This is a high proportion of the total (38,000 genes with 27,500 protein-encoding genes, TAIR10.1 genome assembly) and matches earlier studies (Basbouss-Serhal *et al*. 2015; Bai *et al*. 2017, 2020). On the other hand, there were around 5,000 proteins detected in respective monosomal and polysomal fractions. This included a large proportion of ribosomal proteins and also RNA-binding proteins. The comparison of mRNA level, translational activity, and protein abundance emphasized that selective mRNA translation is a major regulatory mechanism of seed germination (Gallard *et al*. 2014). Transcriptomic studies have documented differential accumulation of stored mRNAs during after-ripening (Basbouss-Serhal *et al*. 2015; Bai *et al*. 2017, 2020). These changes may result from transcriptional activity, mRNA turnover, or differential loading onto polysomes. In our analysis, ribosomes in dry seeds were predominantly in the monosome form, with polysomes absent. Upon 48-hour imbibition, polysome peaks emerged in non-dormant Col but remained undetectable in dormant Cvi seeds, consistent with Bai *et al*. (2017). This underscores the translational activation associated with germination in non-dormant seeds. Our focus on translatome dynamics complements earlier transcriptome studies (Buijs *et al*. 2020; Dekkers *et al*. 2020), which highlighted distinct gene expression profiles between seed compartments (testa, endosperm, and embryo). These compartments contribute differentially to germination, as shown by Dekkers *et al*. (2016), who analyzed dormant and after-ripened Cvi seeds at four time points and across seed compartments. Their work revealed early transcriptional responses in the endosperm, particularly stress-related gene categories, suggesting its protective role in dormant seeds within the soil seed bank.

We also explored mRNA sequence features influencing mono/polysome distribution. U-rich motifs, particularly in the 5′ UTR, were enriched in transcripts associated with polysomes. These motifs, consistent with previous studies (Basbouss-Serhal *et al*. 2015; Bai *et al*. 2017, 2020), may facilitate recruitment of specific RNA-binding proteins (Bai *et al*. 2020, 2021). Structural features such as decreased secondary structure at start and stop codons, known to enhance ribosome accessibility (Kozak, 2005; Kertesz *et al*. 2010; Li *et al*. 2012), were also observed. A methylation of N6-adenosine is one of the most prevalent covalently bound modifications of RNA (Shi *et al*. 2017). It is a dynamic and reversible feature possessing a wide range of regulatory functions (Meyer and Jaffrey, 2014). It was proposed that m6A modification could be involved in the regulation of seed dormancy during after-ripening (Hu, Xu and Kang 2024). In this study, we isolated mRNA from dry and imbibed *Arabidopsis* seeds, and using the m6A antibody, we obtained RNA fragments containing m6A modification. In our IM Col seeds, transcripts with peaks near the stop codon and start codon were found. On the other hand, IM Cvi seeds had the highest m6A peak around the stop codon and decreased peaks at other sites in comparison with IM Col seeds (Fig. 6c,d). Wang et al. (2022) presented an *Arabidopsis* mutant in m6A RNA demethylase (AtALKBH10B) leading to the increase of m6A modifications around the stop codon which negatively regulated gene expression. Moreover, there was a decrease of m6A around the start codon and the rest of CDS. This finding suggested that gene expression was suppressed with a prevalence of m6A modifications around the stop codon. On the contrary, m6A modifications at a position around the start codon were identified in maize and had a positive impact on translation (Luo et al., 2020). Therefore, in analyzed Col and Cvi seeds, there is probably connection between the position of m6A modification around stop and start codons with positive and negative translation regulation, respectively. This is supported with the fact, that Col seeds were germinating after 48h of imbibition, while Cvi seeds were imbibed at the most, suggesting that Col seeds were in a more transcriptional-active stage than Cvi seeds. We annotated transcripts with m6A modification according to the Araport 10 database and analyzed DNA motifs within coding sequences of particular genes (Fig. 6c,d) GAA and CTT tandem repetitions occurred mostly around the start and stop codons. Generally, the CDS contains three-fold nucleotides tandem repeats to avoid a frame-shift mutation (Metzgar *et al*. 2002; Legendre *et al*. 2007). Zhao *et al*. (2014) revealed that in dicots, the most frequent tandem repeats within the CDS are mononucleotides A/T, dinucleotides AT, and trinucleotides AAG/CTT. There is evidence of GAA repetitive sequences found in exons in different species, such as moss (Wu *et al*. 2014), humans and other vertebrates (Tacke and Manley, 1995), and also plants (Thomas *et al*. 2012). All of these were connected to splicing regulation. Proteomic studies of dormant and geminated seeds were conducted by classic 2D gel analysis, resulting in the detection of the most abundant proteins including LEA, seed storage, heat shock and proteins involved in energetic and protein metabolisms (Gallardo *et al*. 2001, Chibani *et al*. 2006) in line with the view of seed stores reserve mobilisation during the germination. Our MS-based study allowed a comprehensive analysis of the entire proteome associated with monosomes or polysomes. In this study we focused on RNA-binding proteins (RBPs) playing a pivotal role in the translational control of gene expression, acting as regulators of mRNA stability, processing, and translation in plants (Hentze *et al*. 2018; Cho *et al*. 2019; Lou *et al*. 2020; Sajeev *et al*. 2022). Among these, Pumilio (PUF) proteins are particularly significant, given their RNA-binding Pumilio-homology domain (PUM-HD), which mediates diverse post-transcriptional processes, including ribosomal RNA (rRNA) processing, mRNA stability, and translation. Our analysis detected several PUMILIO proteins in polysomal fractions, indicating their involvement in active translation during seed germination. The *Arabidopsis* PUMILIO (APUM) protein family consists of 26 members with roles in seed development and stress responses (Francischini and Quaggio, 2009; Tam *et al*. 2010). APUM9 and APUM11 were proposed to function in the regulation of translation of abundant stored mRNAs in imbibing seeds (Xiang *et al*. 2014). APUM24 has been previously implicated in seed maturation and rRNA processing (Huang *et al*. 2021). Its essential role in early embryogenesis and ribosome biogenesis underscores its significance in regulating translation. Mutants of APUM23 and APUM24 demonstrate altered rRNA processing and pleiotropic effects in the form of developmental phenotypes that are similar to those of other genes involved in ribosome biogenesis. Deficiencies in APUM24 expression result in abnormal seed maturation, reduced seed oil content, and embryonic lethality (Shanmugam *et al*. 2017). These findings support our observation of PUM proteins in polysomal fractions, suggesting a role in translation during seed germination.

In conclusion, our findings provide new insights into the translational dynamics underlying seed dormancy and germination. By elucidating the interplay between mRNA storage, post-transcriptional regulation, and polysome recruitment, this study advances our understanding of the molecular mechanisms driving seedling establishment and offers potential targets for improving seed performance in agricultural contexts.

## Acknowledgements

The authors are grateful to Renata Plotzová and Marie Vitásková for their excellent technical assistance.

## DATA AVAILABILITY

Sequencing data have been deposited on the NCBI Gene Expression Omnibus repository (GEO, http://ncbi.nlm.nih.gov/geo), the mass spectrometry proteomics data have been deposited to the ProteomeXchange Consortium via the PRIDE partner repository and will be accessible upon publication.

