## Supplemental tables and figures for "*Arabidopsis* seed stored mRNAs and translation regulation during post-harvest ripening and imbibition"

#### Table S1. List of samples subjected to RNAseq and m6A RIPseq analysis

| sample | genotype | stage | fraction | sequencing | Fraction | Raw reads | Filtered | Mapped | Genes TPM>1 |
|---|---|---|---|---|---|---|---|---|---|
| **R1** | col | AR | whole_seed | RNAseq | WS | 27516440 | 23228603 | 2437794 | 10329 |
| **R3** | cvi | AR | whole_seed | RNAseq | WS | 21131419 | 19083903 | 17237536 | 11538 |
| **R4** | cvi | AR | whole_seed | RNAseq | WS | 21996626 | 19761798 | 17345248 | 10941 |
| **R5** | col | IM | whole_seed | RNAseq | WS | 25564993 | 24072363 | 23216267 | 16832 |
| **R6** | col | IM | whole_seed | RNAseq | WS | 22176947 | 21366999 | 20558813 | 17259 |
| **R7** | cvi | IM | whole_seed | RNAseq | WS | 23742131 | 22311170 | 20359891 | 15519 |
| **R8** | cvi | IM | whole_seed | RNAseq | WS | 27266534 | 25578191 | 22857571 | 14489 |
| **R9** | col | FH | ribo | RNAseq | R | 24346453 | 22698770 | 21921704 | 8611 |
| **R10** | col | FH | ribo | RNAseq | R | 27701705 | 24717268 | 22886752 | 8008 |
| **R11** | col | FH | mono | RNAseq | M | 23116780 | 21625483 | 20901840 | 10164 |
| **R12** | col | FH | mono | RNAseq | M | 22788846 | 21269021 | 20607286 | 10867 |
| **R13** | cvi | FH | ribo | RNAseq | R | 23921587 | 22179353 | 19160856 | 10541 |
| **R14** | cvi | FH | ribo | RNAseq | R | 39100170 | 36476281 | 27488082 | 7270 |
| **R15** | cvi | FH | mono | RNAseq | M | 24040850 | 22340861 | 20397647 | 11246 |
| **R16** | cvi | FH | mono | RNAseq | M | 21765172 | 20011506 | 18594917 | 11812 |
| **R17** | col | FH | extract | RNAseq | EX | 21959030 | 20445035 | 19499717 | 9753 |
| **R18** | cvi | FH | extract | RNAseq | EX | 23756369 | 22171433 | 20518924 | 12608 |
| **R19** | cvi | FH | extract | RNAseq | EX | 29788997 | 27719958 | 25753344 | 12801 |
| **R20** | col | AR | ribo | RNAseq | R | 21264264 | 19885920 | 19074473 | 7995 |
| **R21** | col | AR | mono | RNAseq | M | 21329448 | 19914807 | 19209519 | 11217 |
| **R22** | cvi | AR | ribo | RNAseq | R | 21679806 | 20173988 | 18523874 | 10825 |
| **R23** | cvi | AR | mono | RNAseq | M | 32599961 | 30741052 | 28578761 | 12444 |
| **R24** | cvi | AR | mono | RNAseq | M | 24252233 | 22691600 | 21337927 | 12209 |
| **R25** | col | AR | extract | RNAseq | EX | 22833241 | 21302782 | 20655721 | 13583 |
| **R26** | col | AR | extract | RNAseq | EX | 26170351 | 24483909 | 23736898 | 13666 |
| **R27** | cvi | AR | extract | RNAseq | EX | 19921637 | 18461434 | 17349481 | 12858 |
| **R28** | cvi | AR | extract | RNAseq | EX | 27493857 | 25691013 | 23805447 | 11306 |
| **R29** | col | IM | mono | RNAseq | M | 20881505 | 19954837 | 18719112 | 16431 |
| **R30** | col | IM | mono | RNAseq | M | 25314933 | 24062876 | 22185139 | 17589 |
| **R31** | col | IM | poly | RNAseq | P | 23856336 | 22693687 | 21778872 | 16486 |
| **R32** | col | IM | poly | RNAseq | P | 24411779 | 23301905 | 21942633 | 15584 |
| **R33** | cvi | IM | mono | RNAseq | M | 26243503 | 24905464 | 21950283 | 14821 |
| **R34** | cvi | IM | mono | RNAseq | M | 24697646 | 23246741 | 20584964 | 14985 |
| **R35** | col | IM | extract | RNAseq | EX | 25982247 | 24692181 | 23570374 | 17726 |
| **R36** | col | IM | extract | RNAseq | EX | 25563868 | 24499395 | 23477097 | 16904 |
| **R37** | cvi | IM | extract | RNAseq | EX | 23684989 | 22588546 | 20642592 | 15200 |
| **R38** | cvi | IM | extract | RNAseq | EX | 20965542 | 19463086 | 17867873 | 15363 |
| **A12** | col | IM | whole_seed | RNAseq | WS | 21173443 | 19982666 | 19039064 | 14016 |
| **A13** | col | IM | whole_seed | RNAseq | WS | 20425521 | 19389406 | 18448818 | 15294 |
| **A14** | col | IM | whole_seed | RNAseq | WS | 23197159 | 21925243 | 20852534 | 14182 |
| **A15** | col | AR | whole_seed | RNAseq | WS | 26060471 | 24215078 | 22757361 | 10614 |
| **A16** | col | AR | whole_seed | RNAseq | WS | 25017806 | 23394356 | 22163619 | 10589 |
| **A17** | col | AR | whole_seed | RNAseq | WS | 25853386 | 24208833 | 22919145 | 10872 |
| **A18** | col | IM | m6A | RIP | m6A | 32774012 | 24159691 | 18917327 | 2504 |
| **A19** | col | IM | m6A | RIP | m6A | 29983285 | 23761558 | 18672872 | 5644 |
| **A20** | col | IM | m6A | RIP | m6A | 22421459 | 18947614 | 15009942 | 7792 |
| **A21** | col | IM | C- | RIP | C- | 42442260 | 17956994 | 11275610 | 2766 |
| **A22** | col | AR | m6A | RIP | m6A | 21740303 | 17854606 | 14433890 | 3459 |
| **A23** | col | AR | m6A | RIP | m6A | 21619739 | 19342729 | 17487608 | 3597 |
| **A24** | col | AR | m6A | RIP | m6A | 28718944 | 21126522 | 16389732 | 2875 |
| **A1_23** | col | FH | m6A | RIP | m6A | 20792807 | 19837253 | 17585078 | 1137 |
| **A2_23** | col | FH | m6A | RIP | m6A | 22003684 | 21065479 | 18623838 | 1342 |
| **A3_23** | col | FH | m6A | RIP | m6A | 23535077 | 22327756 | 19293311 | 1359 |
| **A4_23** | cvi | FH | m6A | RIP | m6A | 14301365 | 13629681 | 11542145 | 1533 |
| **A5_23** | cvi | FH | m6A | RIP | m6A | 24615612 | 23359238 | 20342668 | 1022 |
| **A6_23** | cvi | FH | m6A | RIP | m6A | 11936936 | 11375042 | 9839222 | 1238 |
| **A7_23** | cvi | AR | m6A | RIP | m6A | 23386046 | 22458022 | 19832180 | 2281 |
| **A8_23** | cvi | AR | m6A | RIP | m6A | 29585334 | 28407962 | 25186471 | 2179 |
| **A9_23** | cvi | AR | m6A | RIP | m6A | 25407149 | 24247649 | 21490226 | 3995 |
| **A10_23** | cvi | IM | m6A | RIP | m6A | 19508878 | 18514150 | 15145131 | 1745 |
| **A11_23** | cvi | IM | m6A | RIP | m6A | 19635062 | 18711256 | 15734353 | 3212 |
| **A12_23** | cvi | IM | m6A | RIP | m6A | 24698644 | 23633317 | 19745190 | 5221 |
| **A13_23** | col | FH | whole_seed | RNAseq | WS | 22377050 | 21041554 | 19742042 | 10146 |
| **A14_23** | col | FH | whole_seed | RNAseq | WS | 25408900 | 23567992 | 22451371 | 11172 |
| **A15_23** | col | FH | whole_seed | RNAseq | WS | 26885418 | 25022313 | 23855407 | 11428 |
| **A16_23** | cvi | FH | whole_seed | RNAseq | WS | 22909124 | 21443956 | 19748780 | 11811 |
| **A17_23** | cvi | FH | whole_seed | RNAseq | WS | 9686534 | 8823771 | 7787175 | 10214 |
| **A18_23** | cvi | FH | whole_seed | RNAseq | WS | 19893347 | 18960571 | 17071335 | 11857 |
| **A19_23** | cvi | AR | whole_seed | RNAseq | WS | 21324406 | 20354870 | 18531606 | 12271 |
| **A20_23** | cvi | AR | whole_seed | RNAseq | WS | 30212710 | 27984844 | 25307877 | 12209 |
| **A21_23** | cvi | AR | whole_seed | RNAseq | WS | 22961219 | 21911598 | 20111610 | 11903 |
| **A22_23** | cvi | IM | whole_seed | RNAseq | WS | 3140744 | 2925404 | 2630591 | 8593 |
| **A23_23** | cvi | IM | whole_seed | RNAseq | WS | 19978663 | 19075899 | 17244413 | 14822 |
| **A24_23** | cvi | IM | whole_seed | RNAseq | WS | 24207265 | 23145094 | 20734215 | 14937 |

**Table S2.** Characterization of calibration curves of studied modified nucleosides.

| Nucleoside | Regression function | Reliability value (R2) |
|---|---|---|
| N-6-methyladenosine | y = 915443x + 2543.4 | 0.9997 |
| N-1-methyladenosine | y = 117821x – 24.691 | 0.9997 |
| 5-methylcytidine | y = 8128.4x + 70.061 | 0.9945 |
| 8-oxoguanosine | y = 41578x – 11.478 | 0.9997 |

**Table S3.** Selected standards of modified nucleosides and their characteristic retention times and m/z signals (in positive ionization mode).

| Nucleoside | Retention time (min) | Signal (m/z) |
|---|---|---|
| N-6-methyladenosine | 12.74 | 282.1202 |
| N-1-methyladenosine | 7.59 | 282.1202 |
| 5-methylcytidine | 7.69 | 258.1096 |
| 8-oxoguanosine | 10.89 | 300.0948 |

**Table S4:** The comparison of **monosomal fraction** among freshly-harvested, after-ripened and imbibed seeds of **Col-0** and results of the enrichment analysis. FH: freshly-harvested seeds, AR: after-ripened seeds; IM: imbibed seeds, 1:present, 0:absent

| FH | AR | IM | number of proteins | enriched KEEG pathway |
|---|---|---|---|---|
| **1** | **0** | | 130 | no matching pathway |
| **1** | **0** | **0** | 64 | Steroid biosynthesis |
| **1** | | **0** | 455 | Ether lipid metabolism; Cutin suberin and wax biosynthesis; Glycerophospholipid metabolism; Phenylpropanoid biosynthesis; Protein processing in endoplasmic ruticulum; Biosynthesis of secondary metabolites |
| **0** | **1** | | 254 | Nucleotide excision repair |
| **0** | **1** | **0** | 85 | no matching pathway |
| | **1** | **0** | 494 | Glycosphingolipid biosynthesis; Phenylpropanoid biosynthesis; Biosynthesis of secondary metabolites |
| | **0** | **1** | 634 | Mismatch repair; ABC transporters; Ubiquinone and other terpenoid-quinone biosynthesis; Porphyrin metabolism; Phenylpropanoid biosynthesis; Ascorbate and aldarate metabolism |
| **0** | | **1** | 780 | DNA replication; Mismatch repair; Ascorbate and aldarate metabolism; ABC transporters; phenylpropanoid biosynthesis |
| **0** | **0** | **1** | 517 | Mismatch repair; Glucosinolate biosynthesis; Porphyrin metabolism; DNA replication; ABC transporters; Phenylpropanoid biosynthesis |
| **1** | **1** | **0** | 308 | Cutin suberine and wax biosynthesis; Phynylpropanoid biosynthesis; Biosynthesis of secondary metabolites |
| **1** | **0** | **1** | 41 | no matching pathway |
| **0** | **1** | **1** | 118 | Nucleotide excision repair |

**Table S5:** The comparison of **monosomal fraction** among freshly-harvested, after-ripened and imbibed seeds of **Cvi-0** and results of the enrichment analysis. Abbreviations: FH: freshly-harvested seeds, AR: after-ripened seeds; IM: imbibed seeds, 1:present, 0:absent

| **FH** | **AR** | **IM** | **number of proteins** | **enriched KEEG pathway** |
|---|---|---|---|---|
| **1** | **0** | | 56 | Porphyrin metabolism; Spliceosome |
| **1** | **0** | **0** | 13 | Zeatin biosynthesis; Fatty acid degradation; Porphyrin metabolism |
| **1** | | **0** | 291 | Non-homologous end-joining; Zeatin biosynthesis; Ubiquinone and other terpenoid-quinone biosynthesis; Arginine and proline metabolism; Fatty acid biosynthesis |
| **0** | **1** | | 492 | Circadian rhythm-plant; Nitrogen metabolism; Phenylpropanoid biosynthesis; Biosynthesis of secondary metabolites |
| **0** | **1** | **0** | 322 | Biosynthesis of secondary metabolites |
| | **1** | **0** | 837 | Arginine and proline metabolism; Ubiquinone and other terpenoid-quinone biosynthesis; Biosynthesis of nucleotide sugars; Amino szgar and nucleotide sugar metabolism; Fatty acid metabolism; Biosynthesisi of cofactor |
| | **0** | **1** | 95 | no matching pathway |
| **0** | | **1** | 145 | no matching pathway |
| **0** | **0** | **1** | 43 | no matching pathway |
| **1** | **1** | **0** | 236 | Non-homologous end-joining; Ubiquinone and other terpenoid-quinone biosynthesis; Arginine and proline metabolism; Biosynthesis of nucleotide sugars; Amino sugar and nucleotide sugar metabolism |
| **1** | **0** | **1** | 27 | Spliceome |
| **0** | **1** | **1** | 76 | no matching pathway |

**Table S6:** The comparison of **polysomal fraction** among freshly-harvested, after-ripened and imbibed seeds of **Col-0** and results of the enrichment analysis. FH: freshly-harvested seeds, AR: after-ripened seeds; IM: imbibed seeds, 1:present, 0:absent

| **FH** | **AR** | **IM** | **number of proteins** | **enriched KEEG pathway** |
|---|---|---|---|---|
| **1** | **0** | | 201 | Ribosome |
| **1** | **0** | **0** | 37 | Monoterpenoid biosynthesis; Brassinosteroid biosynthesis; Fatty acid biosynthesis; Ribosome |
| **1** | | **0** | 100 | Fatty acid biosynthesis; Phenylpropanoid biosynthesis; Biosynthesis of secondary metabolites |
| **0** | **1** | | 22 | no matching pathway |
| **0** | **1** | **0** | 2 | no matching pathway |
| | **1** | **0** | 28 | Phenylalanine metabolism; Cutin suberine and wax biosynthesis; Phenylpropanoid biosynthesis |
| | **0** | **1** | 1636 | Porphyrin metabolism; Sulfur metabolism; ABC transporters; Phenylpropanoid biosynthesis; Glycerophospholipid metabolism; Oxidative phosphorylation |
| **0** | | **1** | 1652 | DNA replication; Porphyrin metabolism; Folate biosynthesis; Sulfur metabolism; Nucleotide Excision repair; Phenylpropanoid biosynthesis |
| **0** | **0** | **1** | 1167 | Porphyrin metabolism; Sulfur metabolism; DNA replication; Phenylpropanoid biosynthesis; Ascorbate and aldarate metabolism; Biosynthesis of secondary metabolites |
| **1** | **1** | **0** | 18 | Phenylalanine metabolism; Phenylpropanoid biosynthesis |
| **1** | **0** | **1** | 143 | Oxidative phosphorylation; Endocytosis; Ribosome |
| **0** | **1** | **1** | 13 | no matching pathway |

**Table S7:** The comparison of **polysomal fraction** among freshly-harvested, after-ripened and imbibed seeds of **Cvi-0** and results of the enrichment analysis. FH: freshly-harvested seeds, AR: after-ripened seeds; IM: imbibed seeds, 1:present, 0:absent

| **FH** | **AR** | **IM** | **number of proteins** | **enriched KEEG pathway** |
|---|---|---|---|---|
| **1** | **0** | | 96 | no matching pathway |
| **1** | **0** | **0** | 52 | no matching pathway |
| **1** | | **0** | 602 | Thiamine metabolism; Biotin metabolism; Panthothenate and CoA biosynthesis; Arginine and proline metabolism; Photosynthesis; Inositol phosphate metabolism; Fructose and mannose metabolism |
| **0** | **1** | | 255 | Biosynthesis od secondary metabolites |
| **0** | **1** | **0** | 194 | Nitrogen metabolism; Biosynthesis of amino acids; Biosynthesis of secondary metabolites |
| | **1** | **0** | 984 | Non-homologous end-joining; Steroid biosynthesis; Thiamine metabolism; Biotine metabolism; Folate biosynthesis; Photosynthesis; Porphyrin metabolism; Arginine and proline metabolism |
| | **0** | **1** | 73 | Protein processing in endoplasmic reticulum |
| **0** | | **1** | 85 | Protein processing in endoplasmic reticulum |
| **0** | **0** | **1** | 36 | Protein processing in endoplasmic reticulum |
| **1** | **1** | **0** | 448 | Thiamine metabolism; Biotin metabolism; Arginine and proline metabolism; Various types of N-glycan biosynthesis; Pantothenate and CoA biosynthesis; Photosynthesis |
| **1** | **0** | **1** | 22 | no matching pathway |
| **0** | **1** | **1** | 28 | no matching pathway |

### Supplemental Figures

a)

| enzyme | gene | Col-0 FH | Col-0 AR | Col-0 IM | Cvi-1 FH | Cvi-1 AR | Cvi-1 IM |
|---|---|---|---|---|---|---|---|
| *ent-kaurene oxidase* | *AT5G45340* | 0,04 | 0,04 | 0,46 | 0,04 | 0,08 | 0,47 |
| *GA 20-ox* | *AT4G25420* | 0,00 | 0,16 | 0,55 | 0,03 | 0,10 | 0,07 |
| | *AT5G07200* | 0,13 | 0,04 | 4,74 | 0,41 | 0,47 | 3,08 |
| | *AT5G51810* | 0,03 | 0,00 | 0,39 | 0,00 | 0,01 | 0,06 |
| *GA 3-ox* | *AT1G15550* | 0,00 | 0,00 | 0,11 | 0,00 | 0,00 | 0,07 |
| | *AT1G80330* | 0,00 | 0,00 | 0,00 | 0,02 | 0,04 | 0,01 |
| | *AT1G80340* | 0,00 | 0,02 | 2,49 | 0,00 | 0,00 | 1,13 |
| *GA 2-ox* | *AT1G02400* | 1,21 | 0,94 | 0,06 | 0,25 | 0,10 | 0,00 |
| | *AT1G30040* | 0,47 | 0,50 | 0,20 | 0,41 | 0,57 | 0,54 |

b)

| enzyme | gene | Col-0 FH | Col-0 AR | Col-0 IM | Cvi-1 FH | Cvi-1 AR | Cvi-1 IM |
|---|---|---|---|---|---|---|---|
| *AAO3* | *AT2G27150* | 0,06 | 0,13 | 0,00 | 0,22 | 0,16 | 0,04 |
| *xanthoxin dehydrogenase* | *AT1G52340* | 1,00 | 0,35 | 1,30 | 1,01 | 0,63 | 0,77 |
| *NCED* | *AT1G30100* | 0,18 | 0,16 | 0,00 | 0,17 | 0,09 | 0,27 |
| | *AT1G78390* | 0,40 | 0,30 | 0,12 | 0,64 | 0,29 | 0,27 |
| | *AT3G14440* | 0,28 | 0,25 | 0,04 | 0,42 | 0,33 | 0,16 |
| | *AT4G18350* | 0,38 | 0,36 | 0,02 | 0,72 | 0,29 | 0,18 |
| | *AT4G19170* | 32,16 | 22,13 | 0,39 | 19,73 | 10,78 | 0,29 |
| *ZEP* | *AT5G67030* | 8,93 | 6,54 | 0,30 | 10,82 | 6,03 | 1,41 |
| *beta-ring hydroxylase* | *AT1G31800* | 0,00 | 0,00 | 0,45 | 0,06 | 0,02 | 0,22 |
| *beta-carotene 3-hydroxylase* | *AT4G25700* | 8,03 | 8,09 | 27,48 | 10,06 | 14,07 | 10,05 |
| | *AT5G52570* | 28,10 | 29,20 | 0,42 | 43,13 | 44,49 | 1,29 |
| *ABA 8'-hydroxylase* | *AT5G45340* | 0,04 | 0,04 | 0,46 | 0,04 | 0,08 | 0,47 |
| | *AT2G29090* | 2,21 | 1,10 | 0,27 | 1,90 | 0,74 | 1,82 |
| | *AT4G19230* | 0,15 | 0,17 | 0,89 | 0,20 | 0,15 | 0,57 |

**Fig. S1: The heatmap of genes encoding enzymes involved in biosynthesis and catabolism of ABA (a) and GAs (b)** in the dry freshly harvested (FH), dry after-ripened (AR) and imbibed (IM) Col-0 and Cvi-1 seeds. The heatmap is based on average TMM (Trimmed Mean of M values) values from RNA sequencing. The red colours are used to represent larger values, while the green colours represent smaller values of gene expression for each group of genes encoding the same enzyme. Legend: ZEP: zeaxanthin epoxidase; NCED: 9-*cis*-epoxycarotenoid dioxygenase; SDR1: short-chain dehydrogenase reductase; AAO3: abscisic aldehyde oxidase.

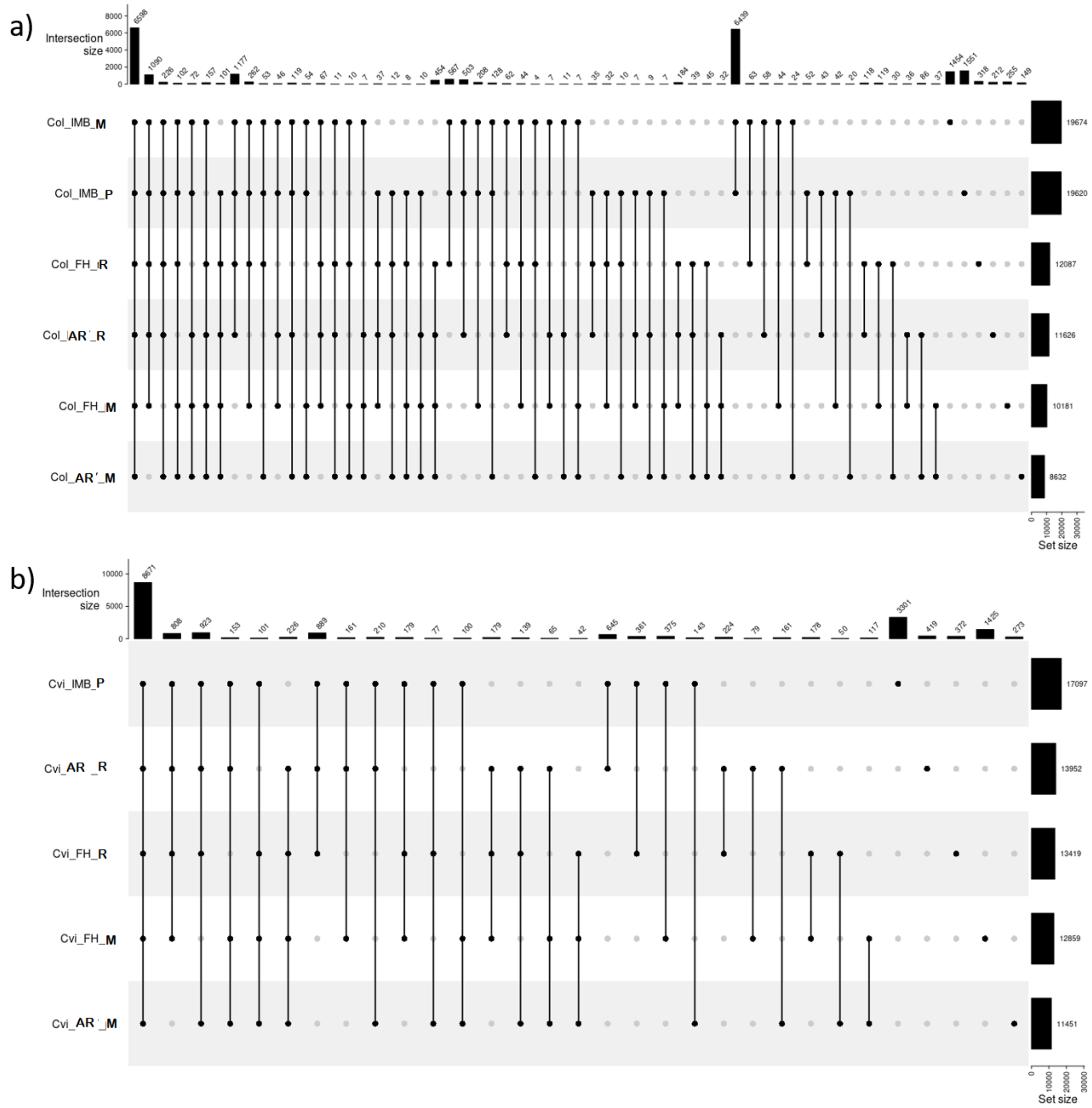

**Fig. S2: Upset plot representing the relationship of pre-monosomal, monosomal, and polysomal genes from freshly harvested (FH), after-ripened (AR), and imbibed (IM) Col and Cvi seeds.** R- pre-monosomal fraction, M- monosomal fraction, P- polysomal fraction.

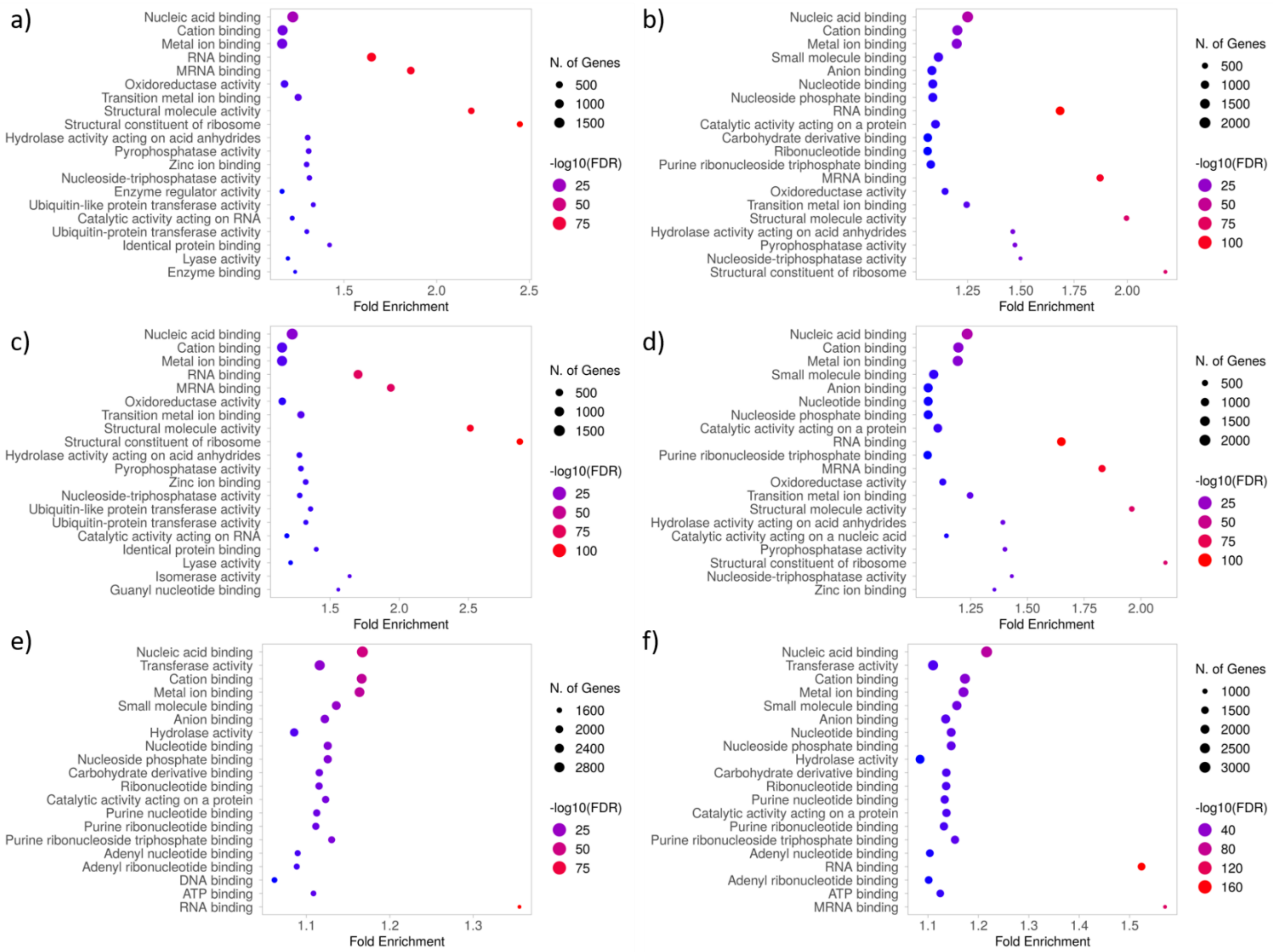

**Fig. S3: Molecular function** enrichment GO terms analysis of genes shared by fractions isolated from particular genotypes within the specific stages. a) Col FH, b) Cvi FH, c) Col AR, d) Cvi AR, e) Col IMBI, f) Cvi IMBI – only one fraction.

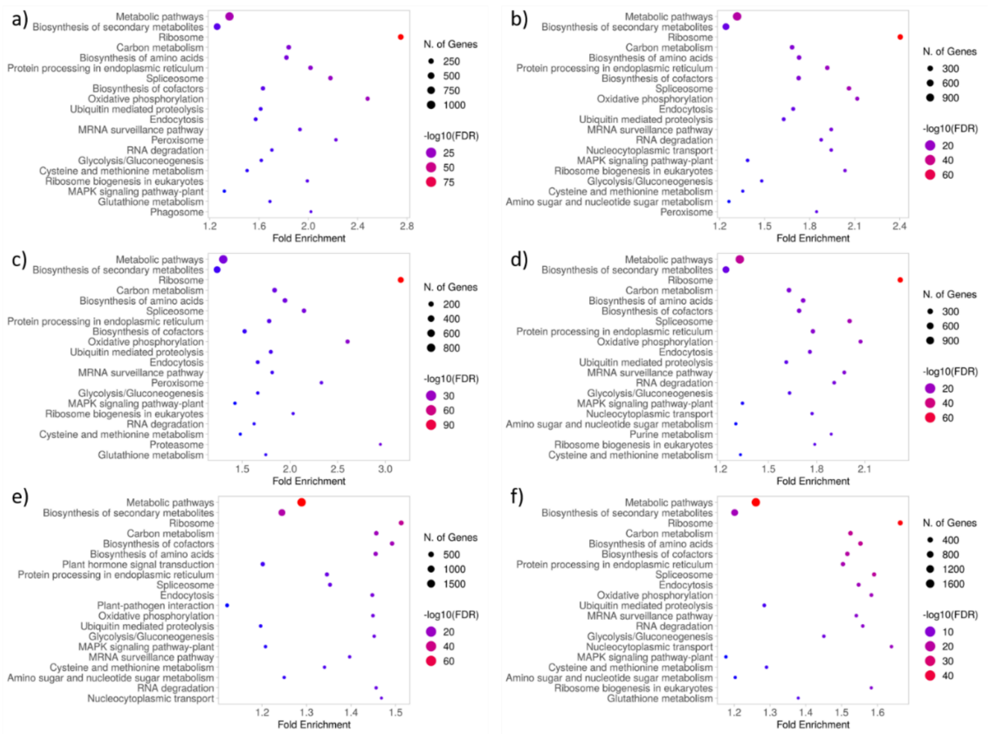

**Fig. S4: KEGG pathway** enrichment for shared genes between fractions isolated from particular genotypes within the specific stages. a) Col FH, b) Cvi FH, c) Col AR, d) Cvi AR, e) Col IMBI, f) Cvi IMBI – only one fraction.

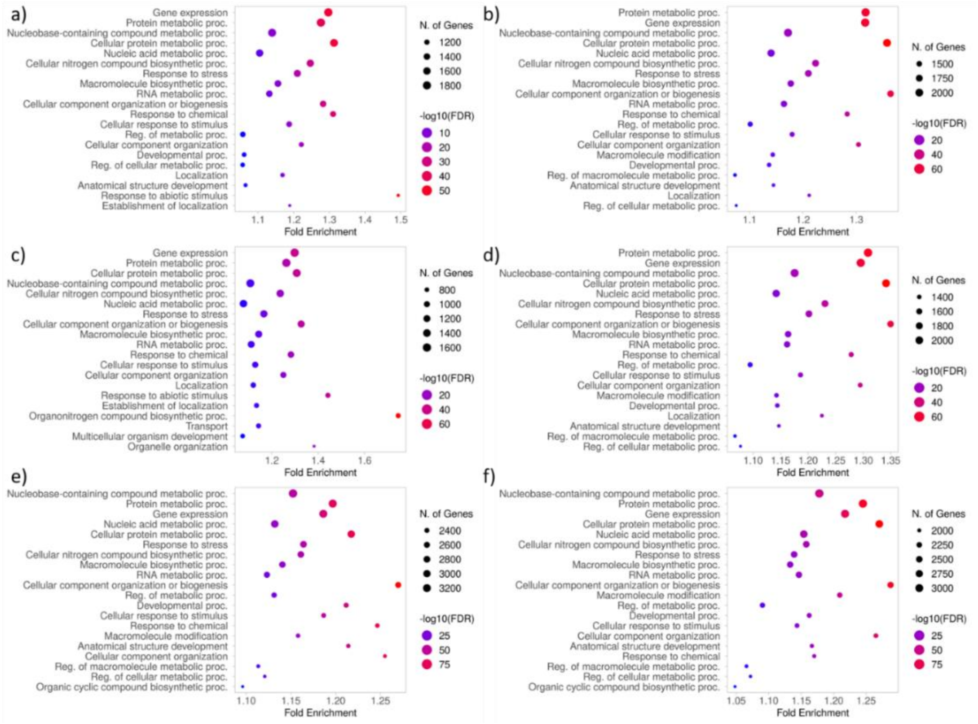

**Fig. S5: Biological process** enrichment GO terms analysis of genes shared by fractions isolated from particular genotypes within the specific stages. a) Col FH, b) Cvi FH, c) Col AR, d) Cvi AR, e) Col IMBI, f) Cvi IMBI – only one fraction.

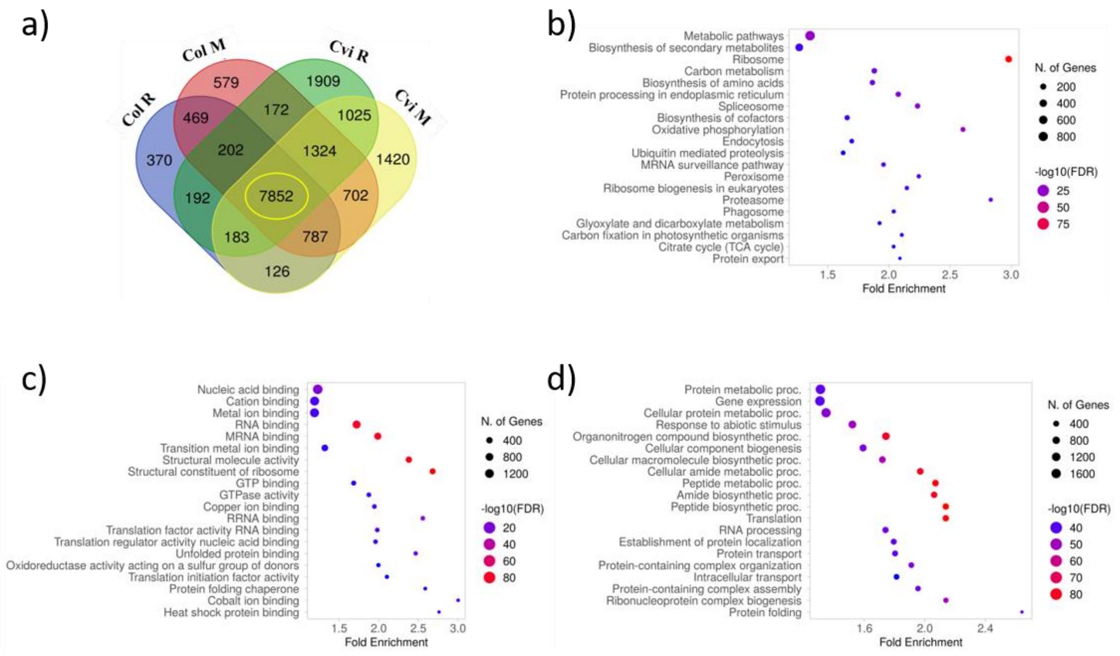

**Fig. S6:** GO term analysis of genes isolated from pre-mono- and monosomal fractions, shared by Col and Cvi genotypes at the freshly harvested stage. a) The Venn diagram represents numbers of genes shared or unique for particular genotypes and fractions. The yellow circle shows a group of analyzed genes. b) The KEGG pathway enrichment. c) The molecular functions enrichment. d) The biological process enrichment.

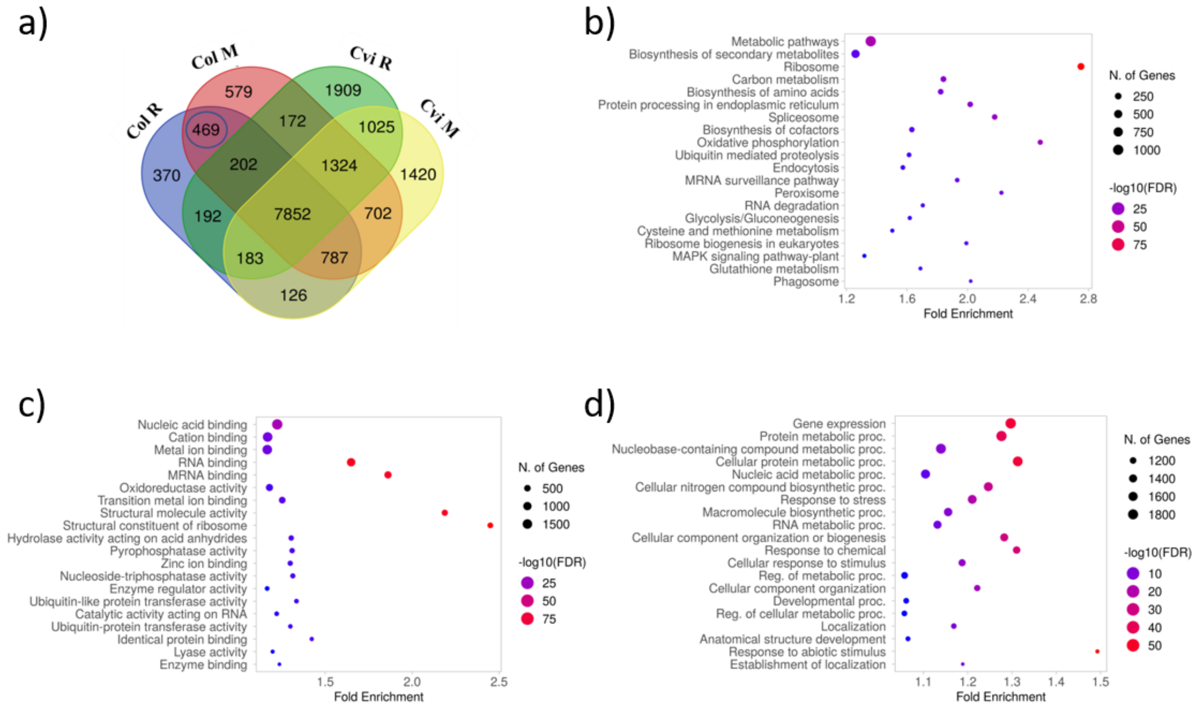

**Fig. S7:** GO term analysis of common genes isolated from pre-mono- and monosomal fractions of Col genotype at the freshly harvested stage. a) The Venn diagram represents numbers of genes shared or unique for particular genotypes and fractions. The blue circle shows a group of analyzed genes. b) The KEGG pathway enrichment. c) The molecular functions enrichment. d) The biological process enrichment.

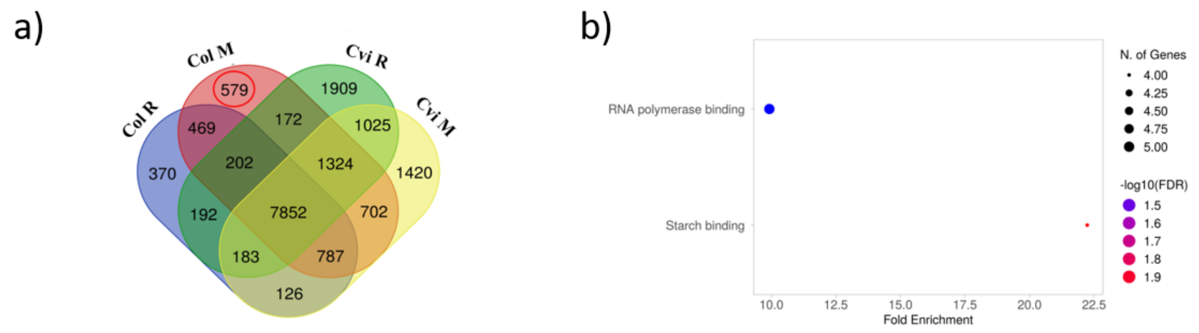

**Fig. S8:** GO term analysis of unique genes isolated from monosomal fraction of Col genotype at the freshly harvested stage. a) The Venn diagram represents numbers of genes shared or unique for particular genotypes and fractions. The red circle shows a group of analyzed genes. b) The molecular functions enrichment.

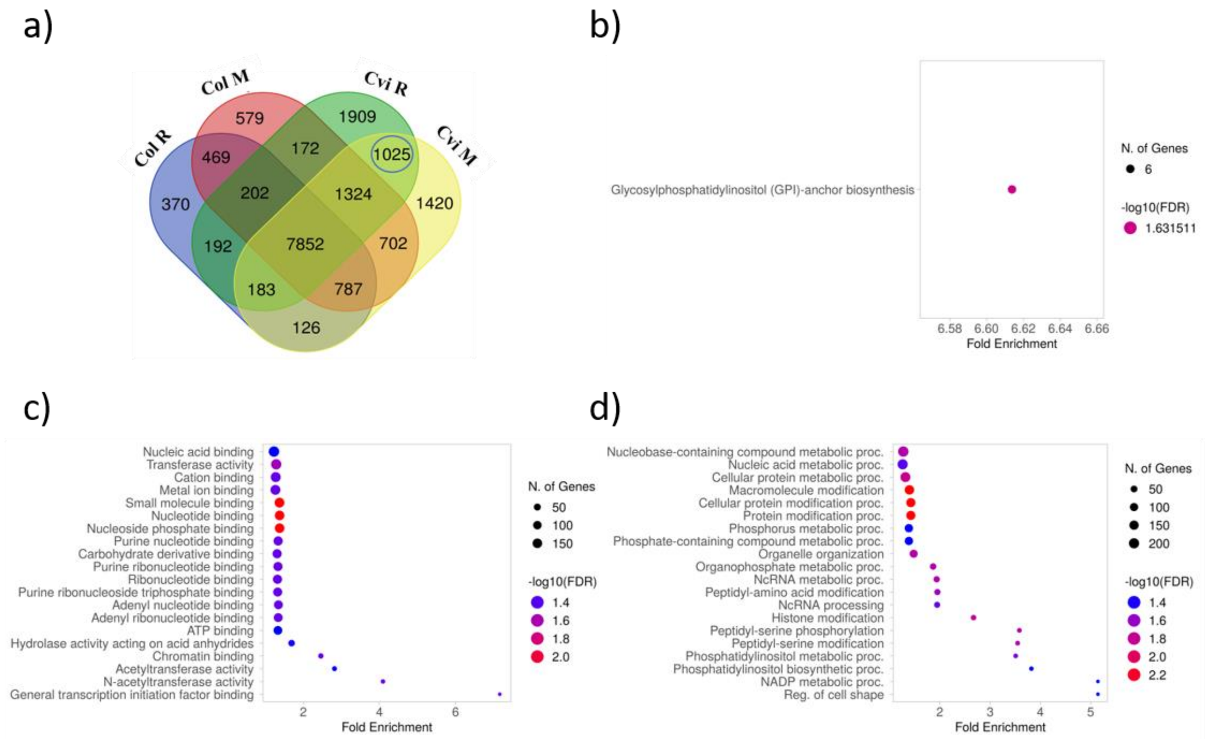

**Fig. S9:** GO term analysis of common genes isolated from pre-mono- and monosomal fractions of Cvi genotype at the freshly harvested stage. a) The Venn diagram represents numbers of genes shared or unique for particular genotypes and fractions. The blue circle shows a group of analyzed genes. b) The KEGG pathway enrichment. c) The molecular functions enrichment. d) The biological process enrichment.

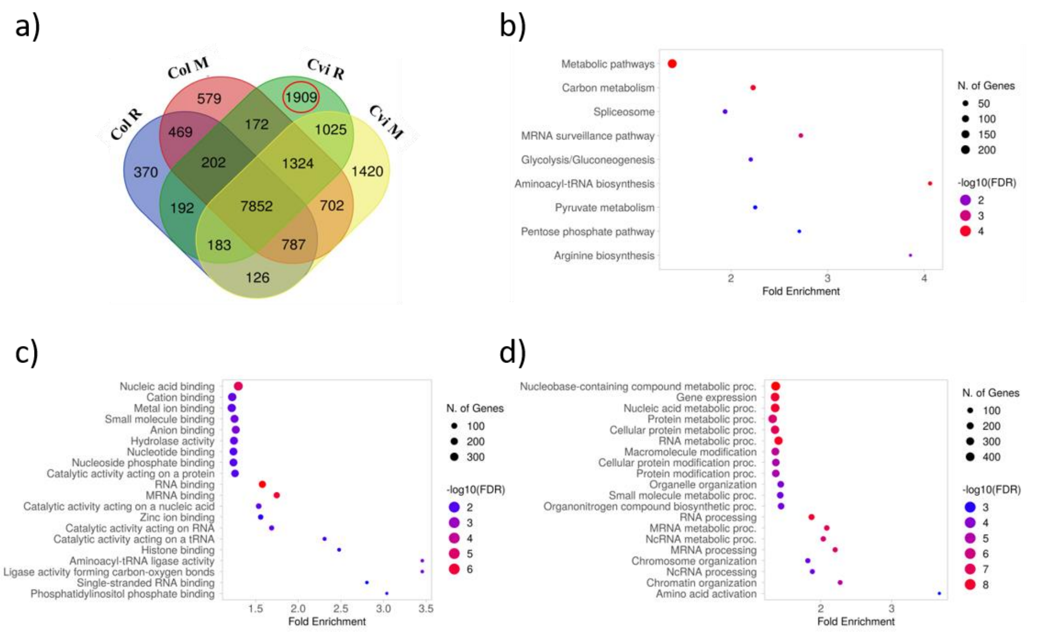

**Fig. S10:** GO term analysis of unique genes isolated from pre-monosomal fraction of Cvi genotype at the freshly harvested stage. a) The Venn diagram represents numbers of genes shared or unique for particular genotypes and fractions. The red circle shows a group of analyzed genes. b) The KEGG pathway enrichment. c) The molecular functions enrichment. d) The biological process enrichment.

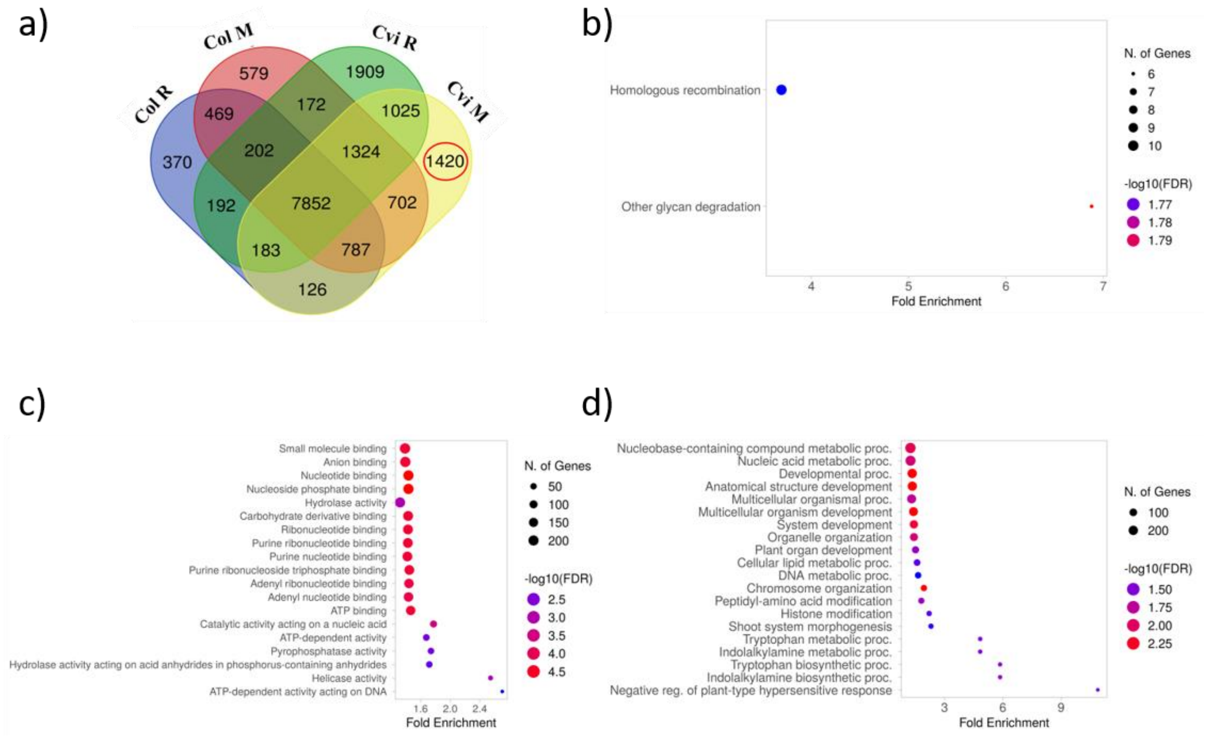

**Fig. S11:** GO term analysis of unique genes isolated from monosomal fraction of Cvi genotype at the freshly harvested stage. a) The Venn diagram represents numbers of genes shared or unique for particular genotypes and fractions. The red circle shows a group of analyzed genes. b) The KEGG pathway enrichment. c) The molecular functions enrichment. d) The biological process enrichment.

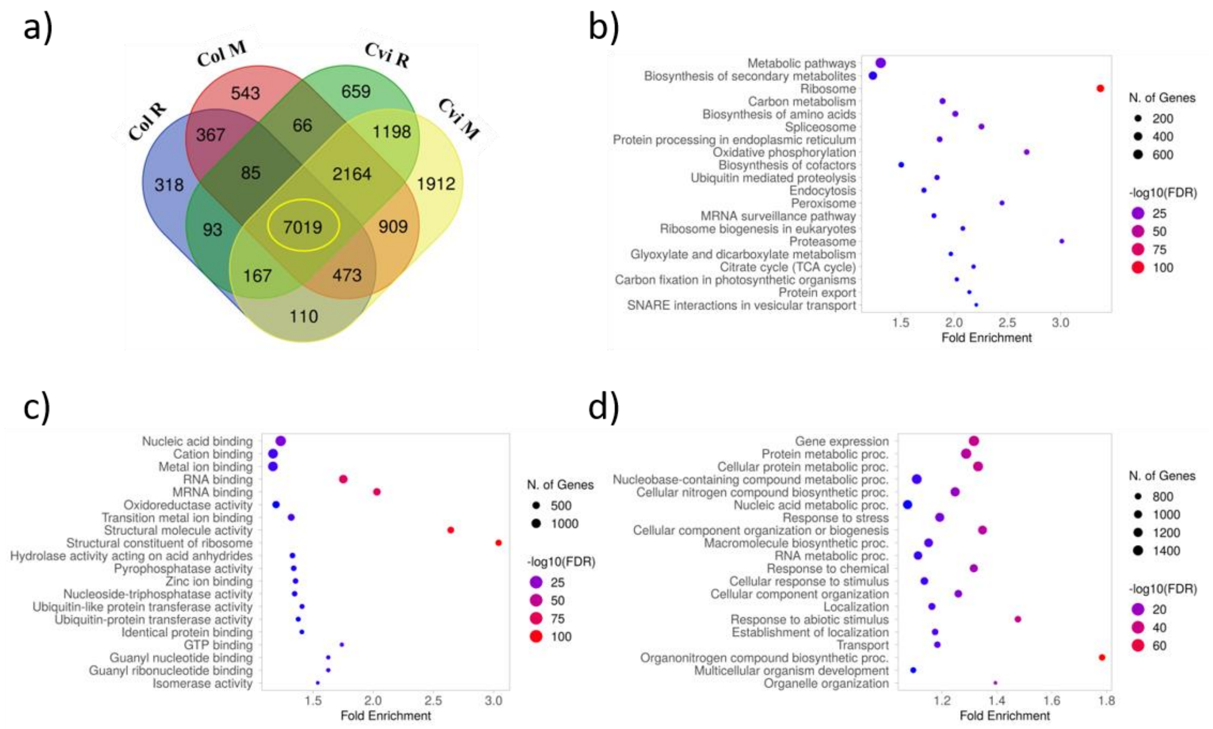

**Fig. S12:** GO term analysis of genes isolated from pre-mono- and monosomal fractions, shared by Col and Cvi genotypes at the after ripen stage. a) The Venn diagram represents numbers of genes shared or unique for particular genotypes and fractions. The yellow circle shows a group of analyzed genes. b) The KEGG pathway enrichment. c) The molecular functions enrichment. d) The biological process enrichment.

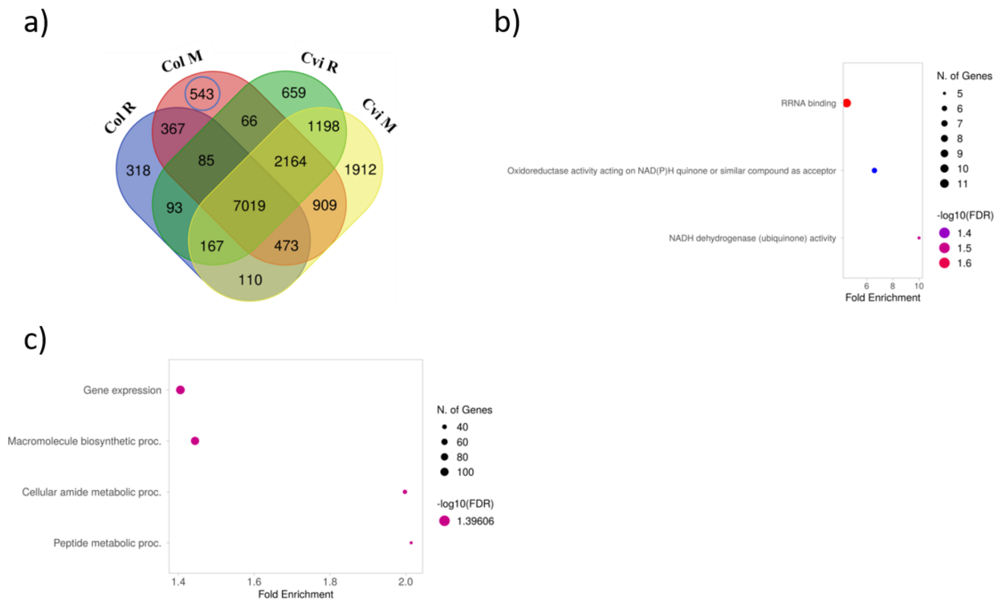

**Fig. S13:** GO term analysis of unique genes isolated from monosomal fractions of Col genotype at the after ripen stage. a) The Venn diagram represents numbers of genes shared or unique for particular genotypes and fractions. The blue circle shows a group of analyzed genes. b) The molecular functions enrichment. c) The biological process enrichment.

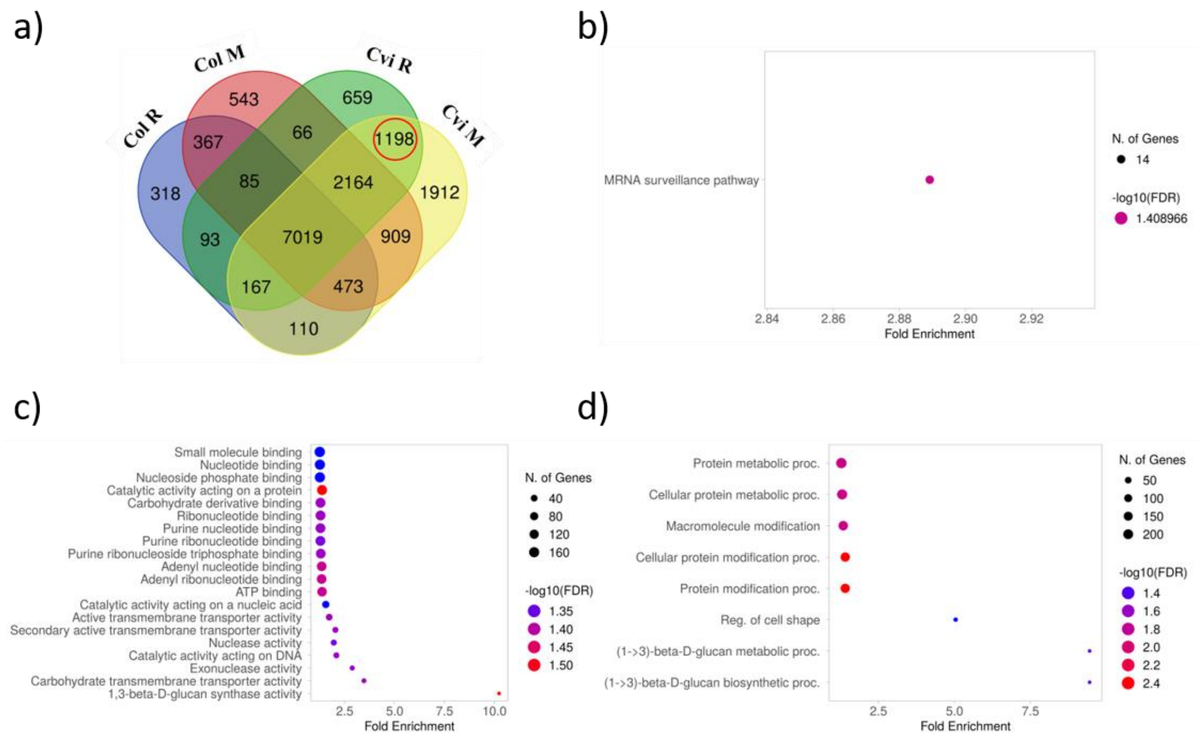

**Fig. S14:** GO term analysis of common genes isolated from pre-mono- and monosomal fractions of Cvi genotype at AR stage. a) The Venn diagram represents numbers of genes shared or unique for particular genotypes and fractions. The red circle shows a group of analyzed genes. b) The KEGG pathway enrichment. c) The molecular functions enrichment. d) The biological process enrichment.

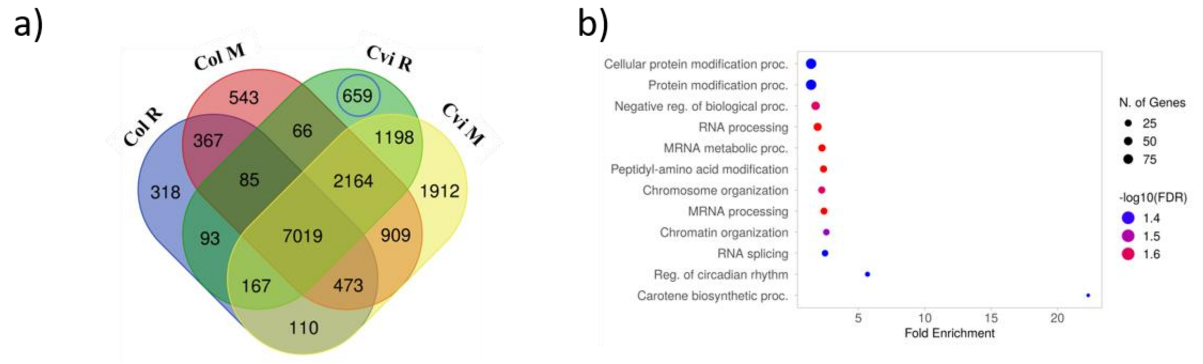

**Fig. S15:** GO term analysis of unique genes isolated from pre-monosomal fractions of Cvi genotype at the after ripen stage. a) The Venn diagram represents numbers of genes shared or unique for particular genotypes and fractions. The blue circle shows a group of analyzed genes. b) The molecular functions enrichment. c) The biological process enrichment.

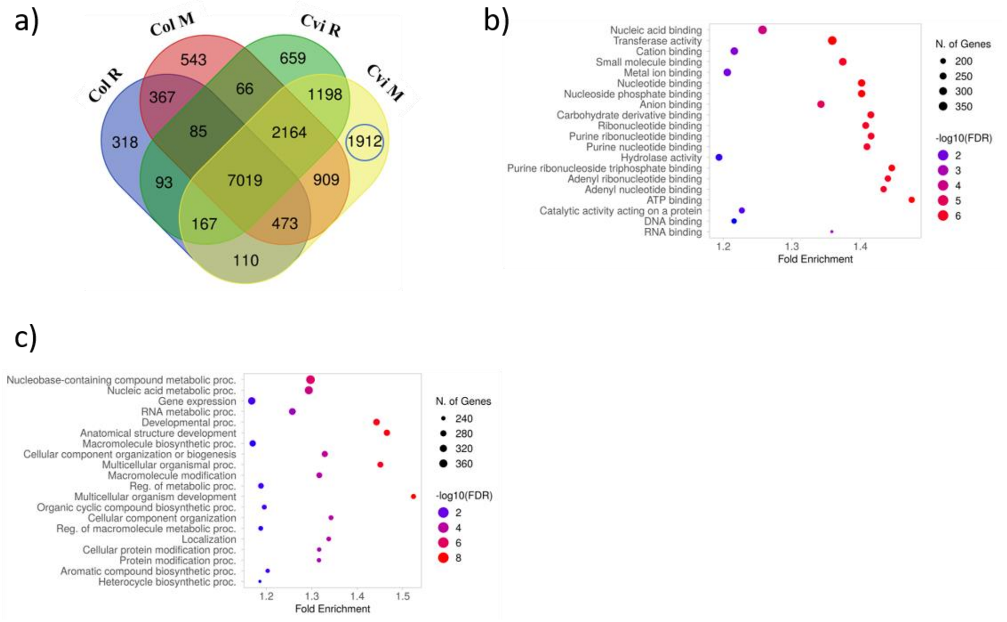

**Fig. S16:** GO term analysis of unique genes isolated from monosomal fractions of Cvi genotype at the after ripen stage. a) The Venn diagram represents numbers of genes shared or unique for particular genotypes and fractions. The blue circle shows a group of analyzed genes. b) The molecular functions enrichment. c) The biological process enrichment.

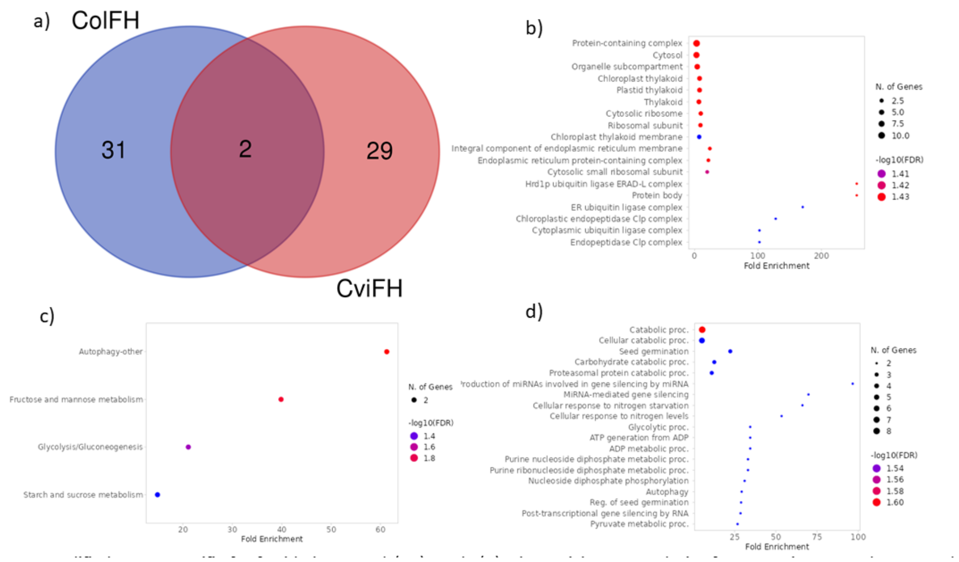

**Fig. S17:** M6A-modified genes specific for freshly harvested (FH) seeds (a). The enrichment analysis of genes unique to Col FH seeds showed as GO Cellular component (a). The genes unique to Cvi FH seeds described using GO terms KEEG (c) and Biological Process (d).

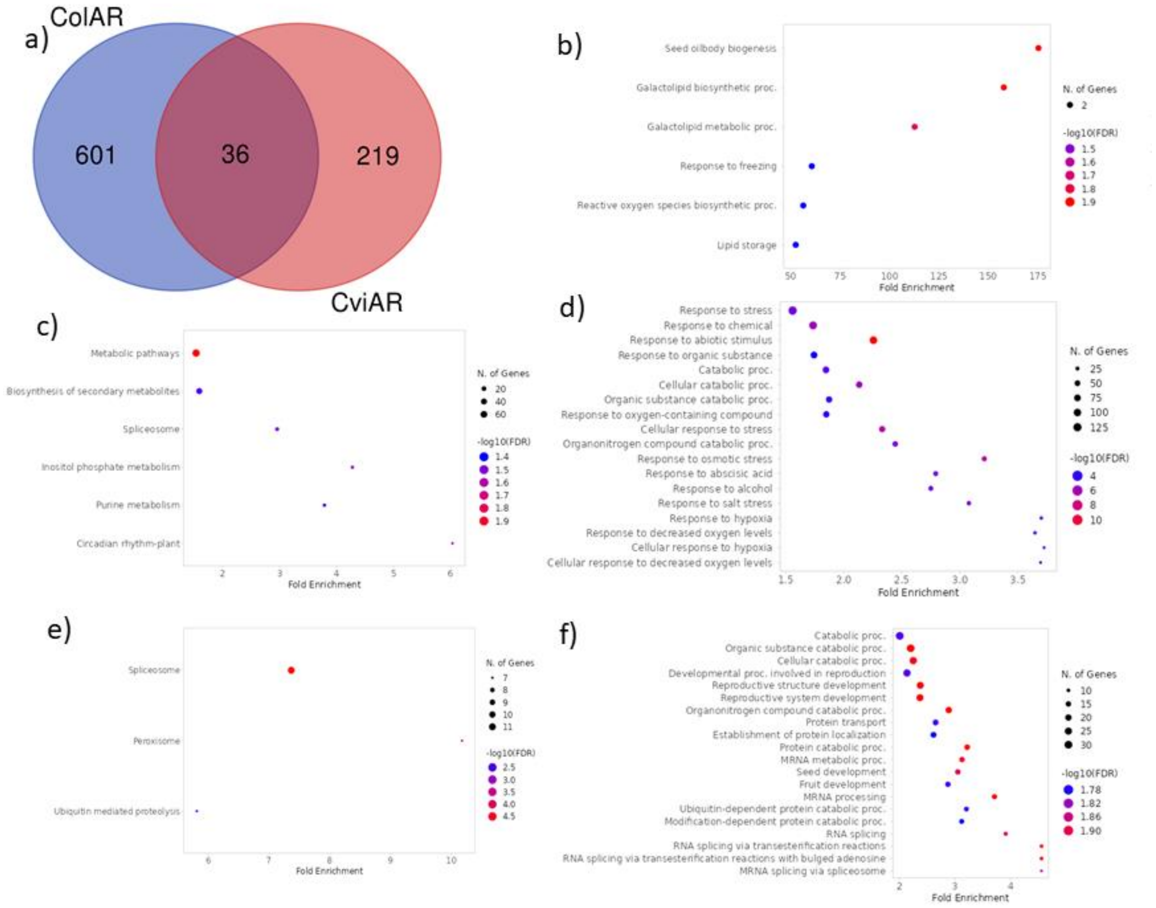

**Fig. S18:** M6A-modified genes specific for after-ripened (AR) seeds (a). The enrichment analysis of genes shared by Col and Cvi AR seeds (Biological Process, b), unique to Col-0 seeds (KEEG:c, Biological Function:d) and unique to Cvi-1 seeds (KEEG: e, Biological process:f).

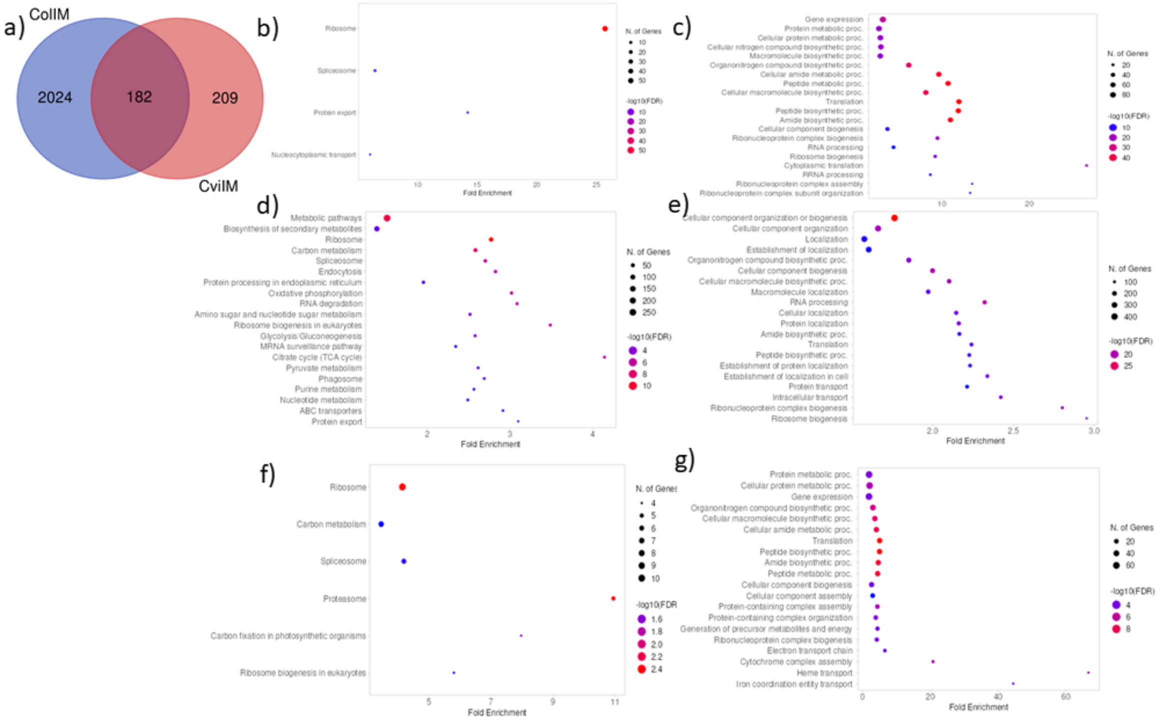

**Fig. S19:** M6A-modified genes specific for imbibed (IM) seeds (a). The enrichment analysis of genes shared by Col-0 and Cvi AR seeds (KEEG:b, Biological Process: c), unique to Col-0 seeds (KEEG:d, Biological Process:e) and unique to Cvi-1 seeds (KEEG: f, Molecular function:g).

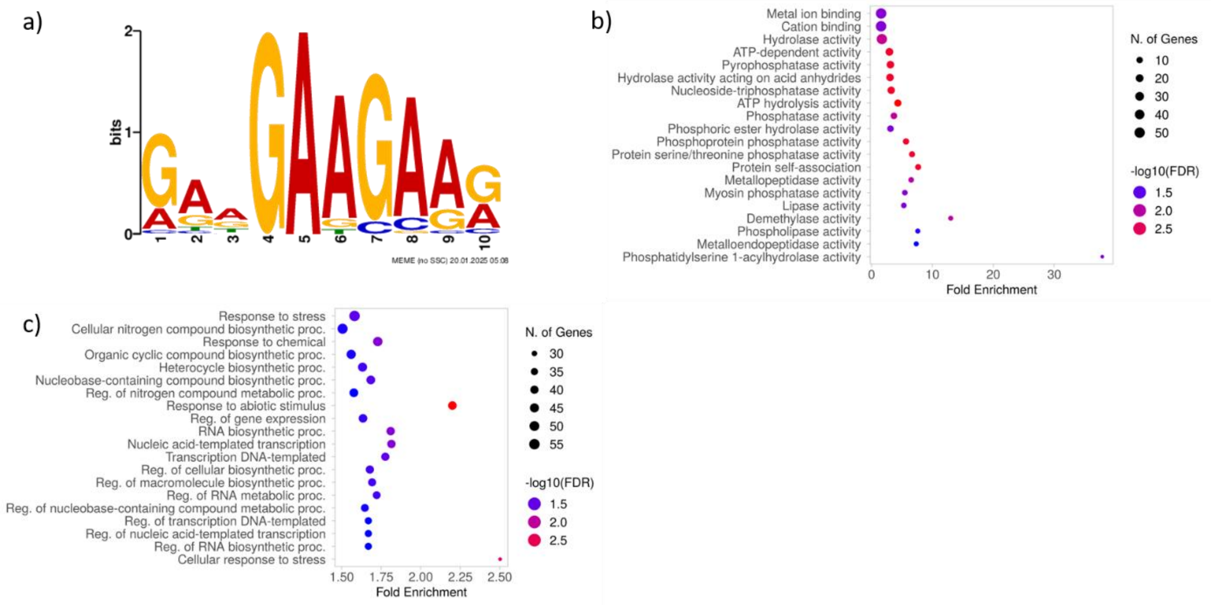

**Fig. S20:** Characterisation of genes with specific motifs, upregulated in AR Col. a) The GAAGAAGAAG motif identified at TSS+25 site. b) The Molecular function enrichment. c) The Biological process enrichment.

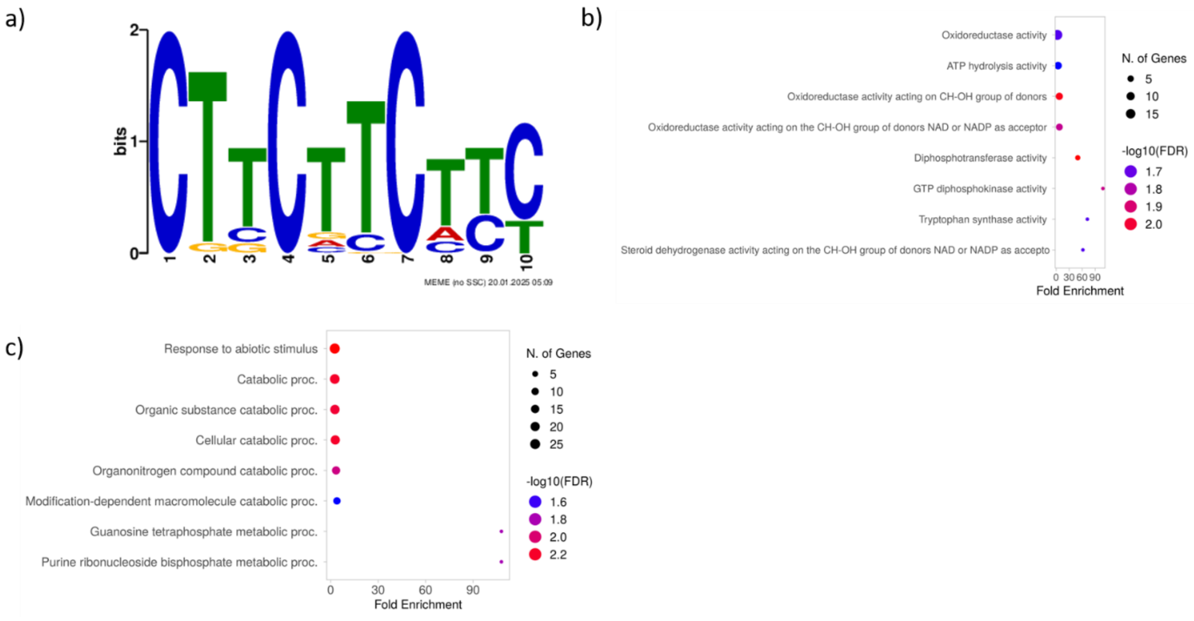

**Fig. S21**: Characterisation of genes with specific motifs, upregulated in AR Col. a) The CTTCTTCTTC motif identified at TSS+25 site. b) The Molecular function enrichment. c) The Biological process enrichment.

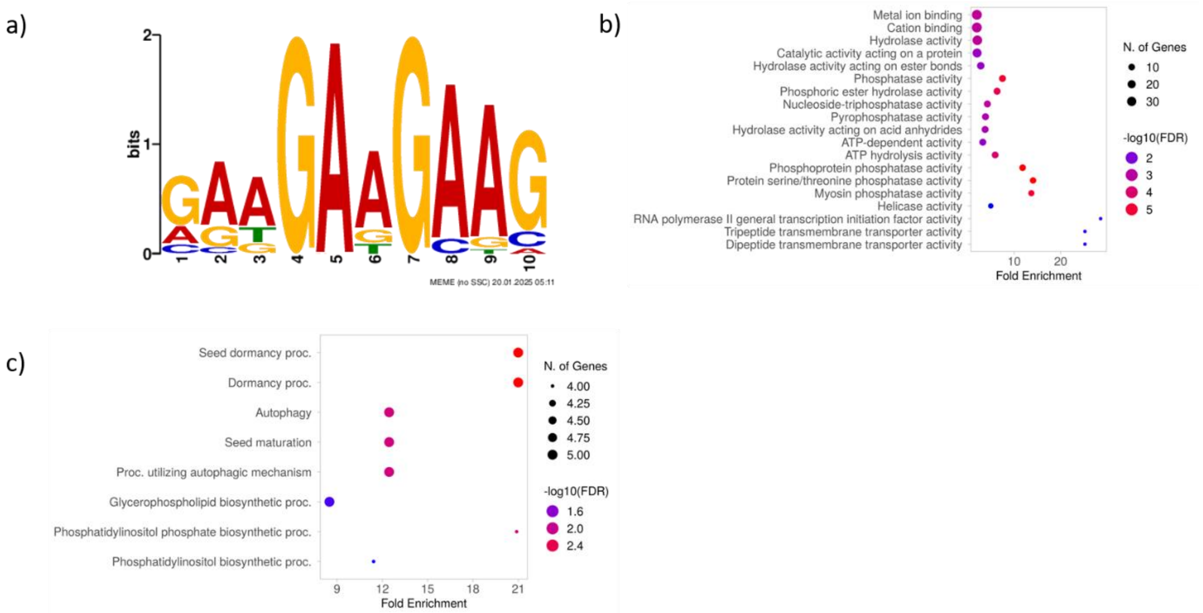

**Fig. S22**: Characterisation of genes with specific motifs, upregulated in AR Col. a) The GAAGAAGAAG motif identified at TSS-15 site. b) The Molecular function enrichment. c) The Biological process enrichment.

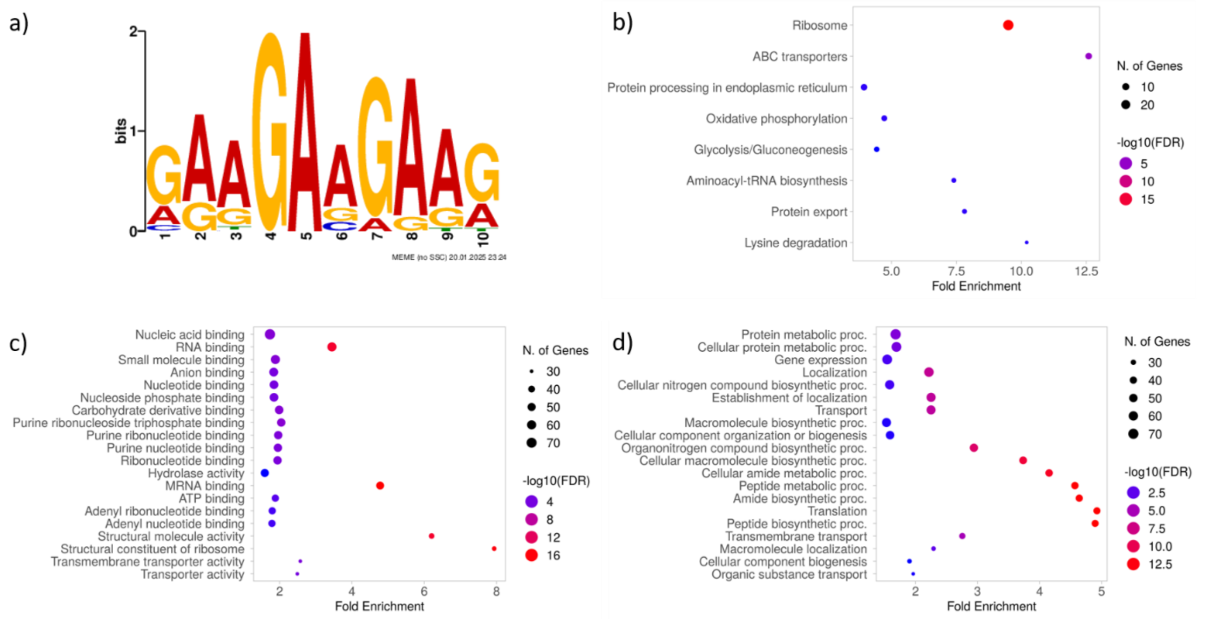

**Fig. S23**: Characterisation of genes with specific motifs, upregulated in IM Col. a) The GAAGAAGAAG motif identified at TSS+25 site. b) The KEGG pathway enrichment. c) The Molecular function enrichment. d) The Biological process enrichment.

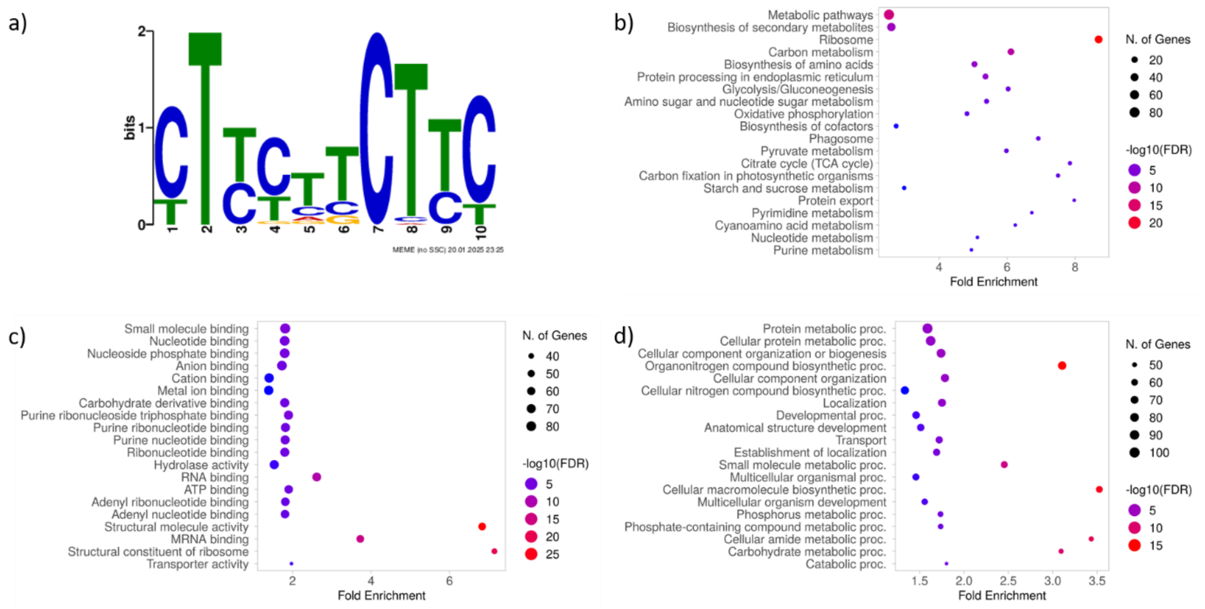

**Fig. S24**: Characterisation of genes with specific motifs, upregulated in IM Col. a) The CTTCTTCTTC motif identified at TSS+25 site. b) The KEGG pathway enrichment. c) The Molecular function enrichment. d) The Biological process enrichment.

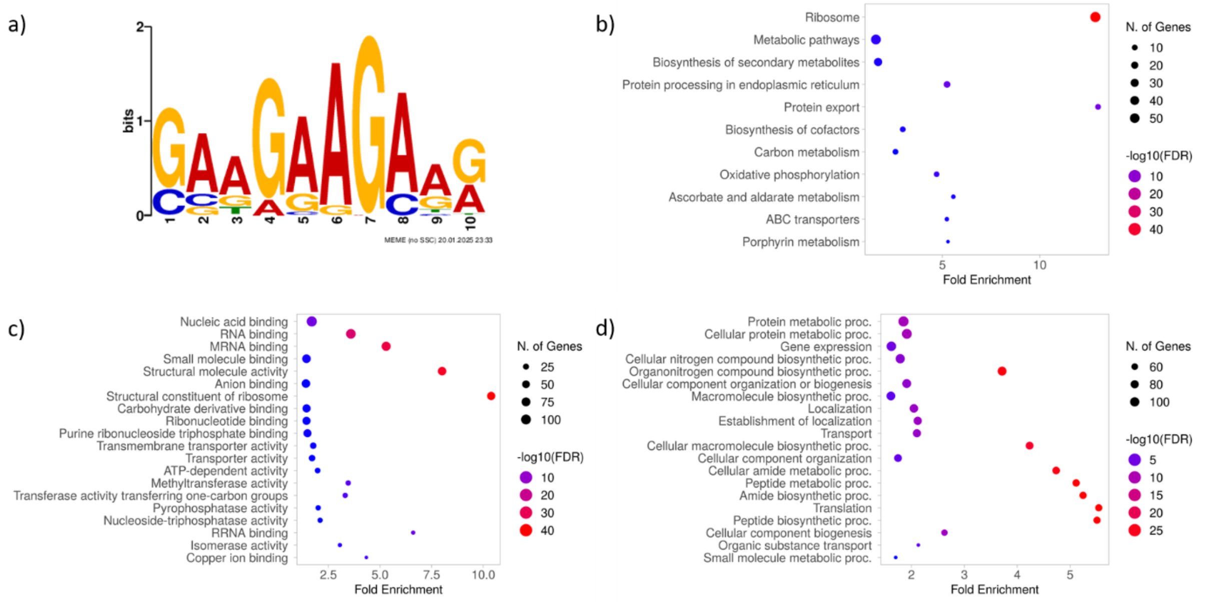

**Fig. S25**: Characterisation of genes with specific motifs, upregulated in IM Col. a) The GAAGAAGAAG motif identified at TSS-15 site. b) The KEGG pathway enrichment. c) The Molecular function enrichment. d) The Biological process enrichment.

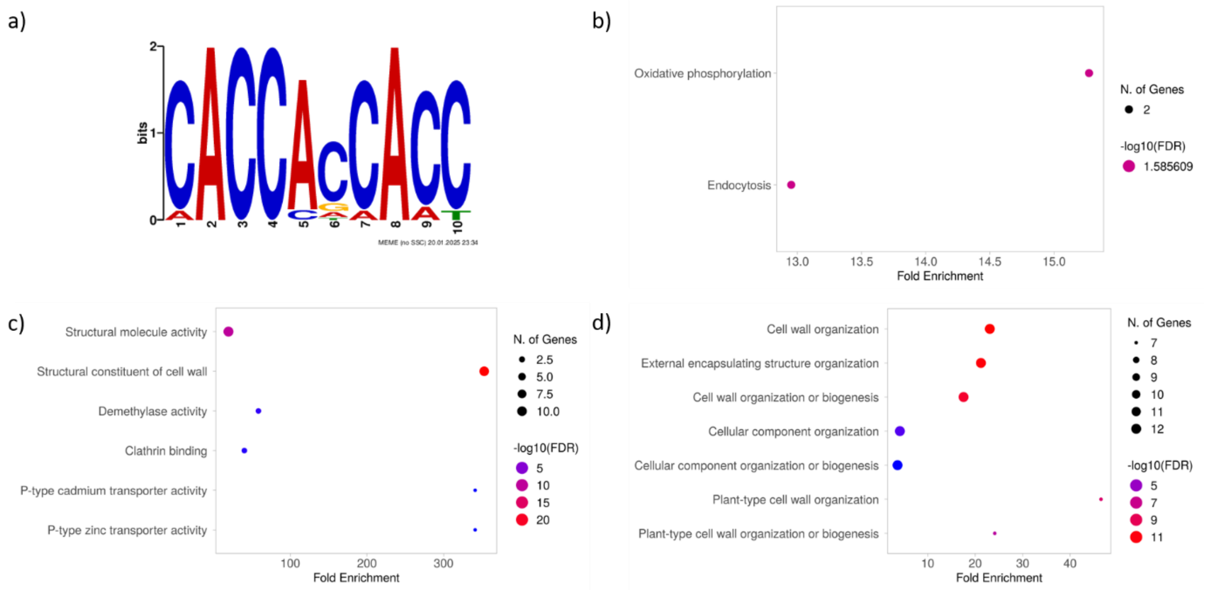

**Fig. S26**: Characterisation of genes with specific motifs, upregulated in IM Col. a) The CACCACCACC motif identified at TSS-15 site. b) The KEGG pathway enrichment. c) The Molecular function enrichment. d) The Biological process enrichment.

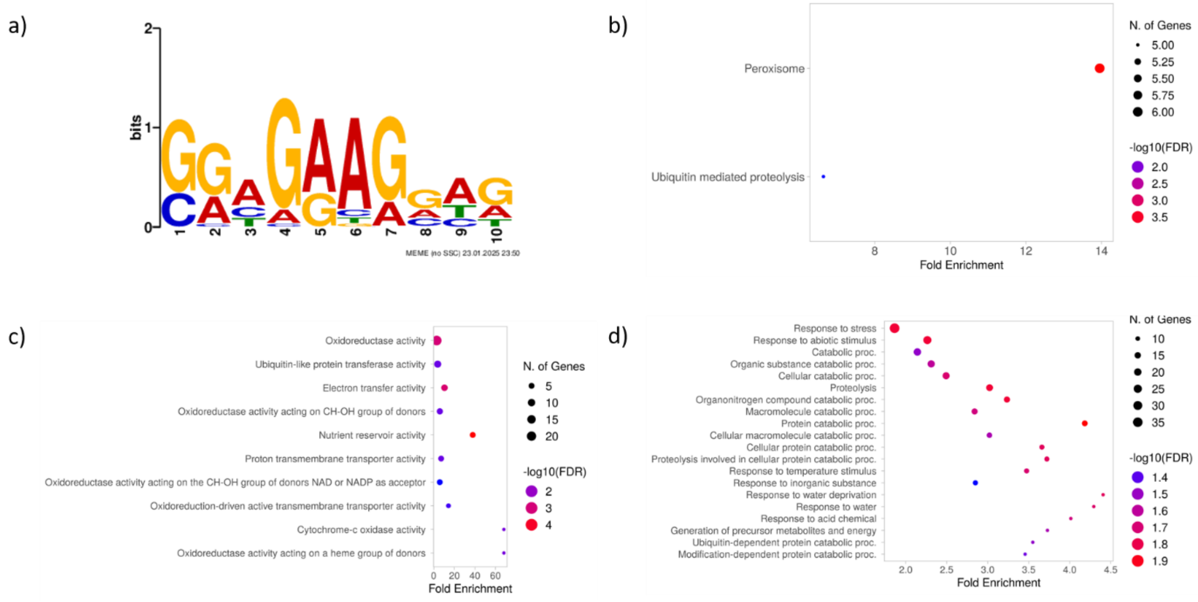

**Fig. S27**: Characterisation of genes with specific motifs, upregulated in AR Cvi. a) The GGAGAAGGAG motif identified at TSS+25 site. b) The KEGG pathway enrichment. c) The Molecular function enrichment. d) The Biological process enrichment.

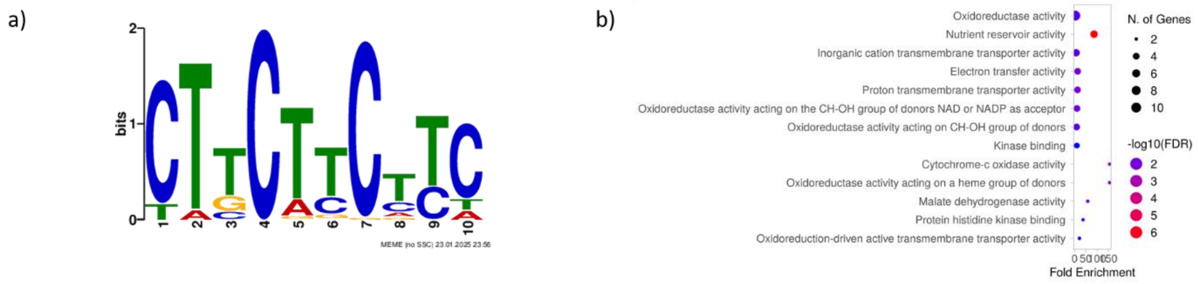

**Fig. S28:** Characterisation of genes with specific motifs, upregulated in AR Cvi. a) The CTTCTTCTTC motif identified at TSS+25 site. b) The KEGG pathway enrichment. c) The Molecular function enrichment. d) The Biological process enrichment.

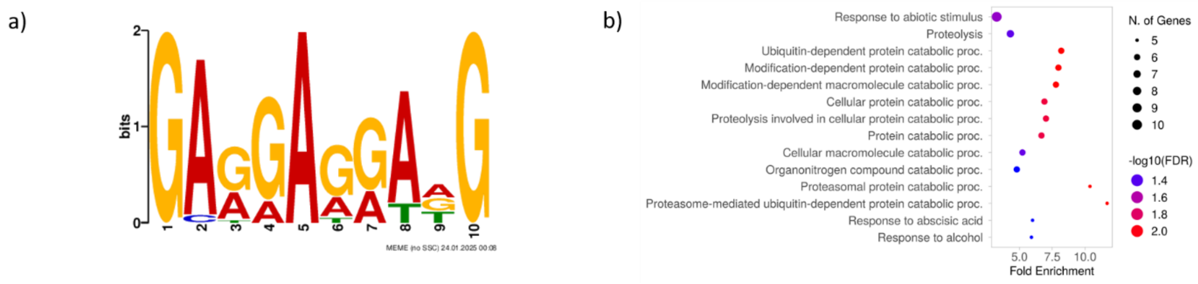

**Fig. S29:** Characterisation of genes with specific motifs, upregulated in AR Cvi. a) The GAGGAGGAAG motif identified at TSS-15 site. b) The KEGG pathway enrichment. c) The Molecular function enrichment. d) The Biological process enrichment.

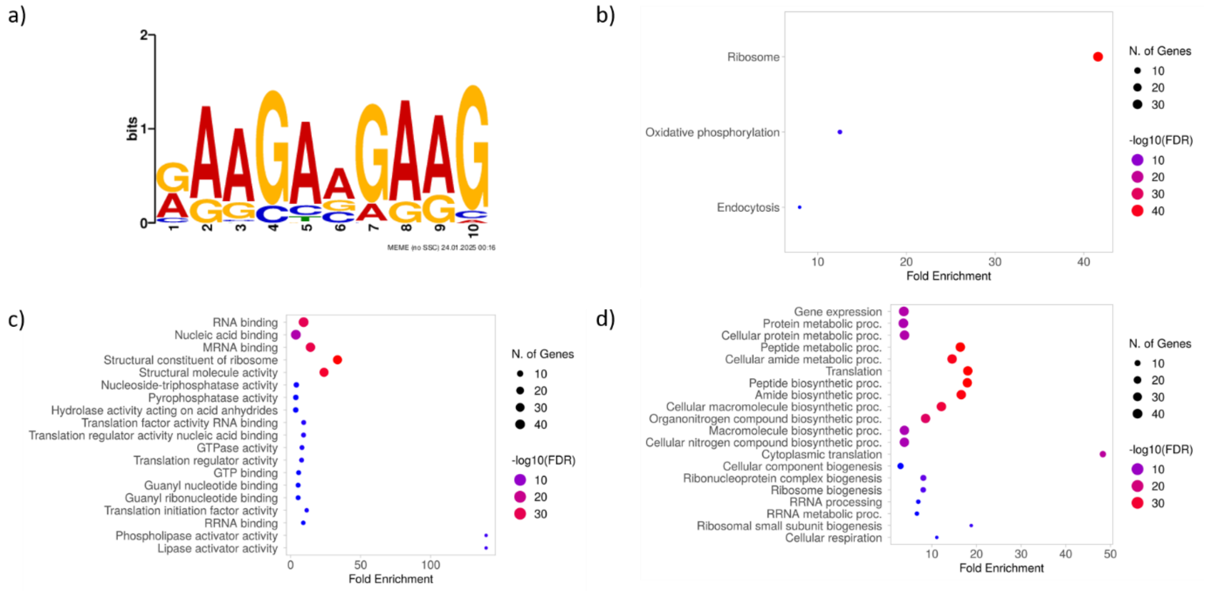

**Fig. S30:** Characterisation of genes with specific motifs, upregulated in IM Cvi. a) The GAAGAAGAAG motif identified at TSS+25 site. b) The KEGG pathway enrichment. c) The Molecular function enrichment. d) The Biological process enrichment.

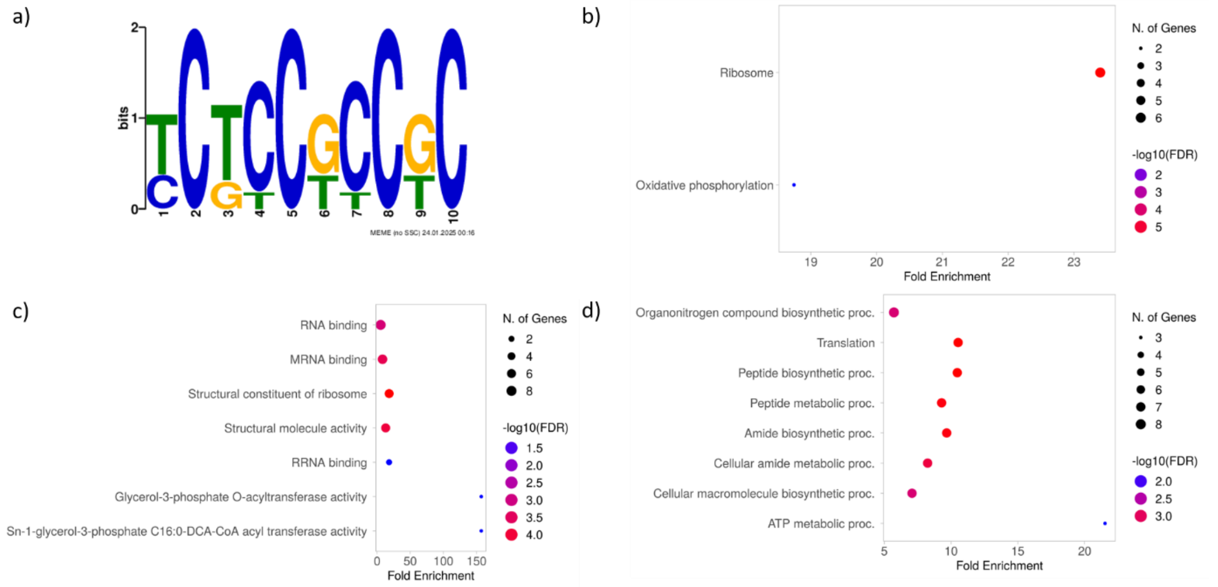

**Fig. S31:** Characterisation of genes with specific motifs, upregulated in IM Cvi. a) The TCTCCGCCGC motif identified at TSS+25 site. b) The KEGG pathway enrichment. c) The Molecular function enrichment. d) The Biological process enrichment.

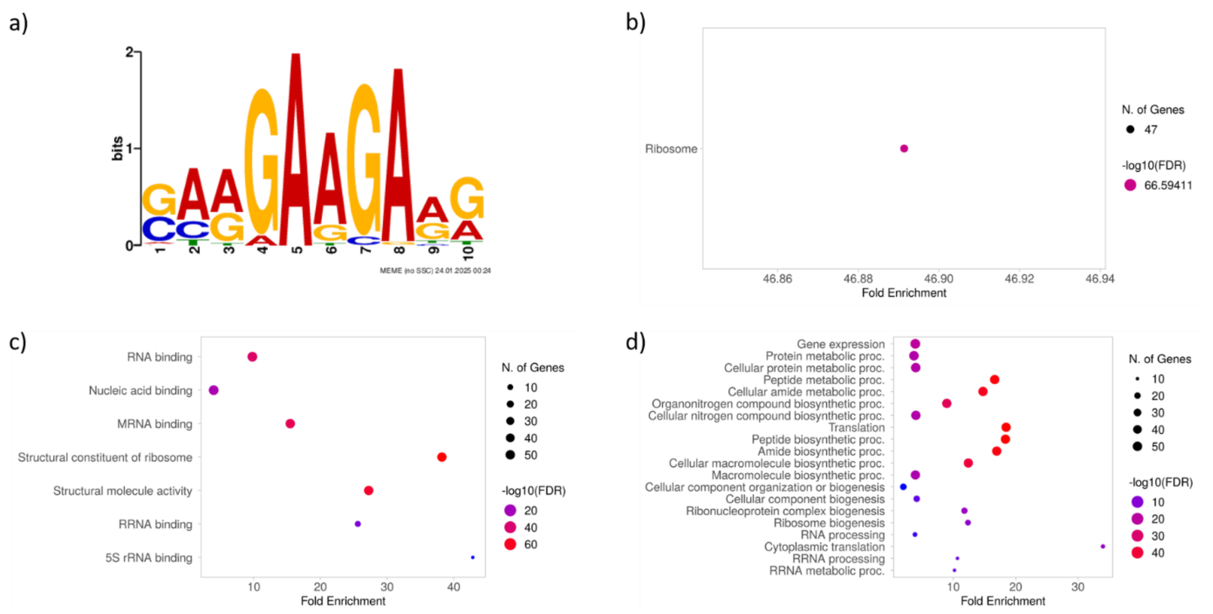

**Fig. S32:** Characterisation of genes with specific motifs, upregulated in IM Cvi. a) The GAAGAAGAAG motif identified at TSS-15 site. b) The KEGG pathway enrichment. c) The Molecular function enrichment. d) The Biological process enrichment.

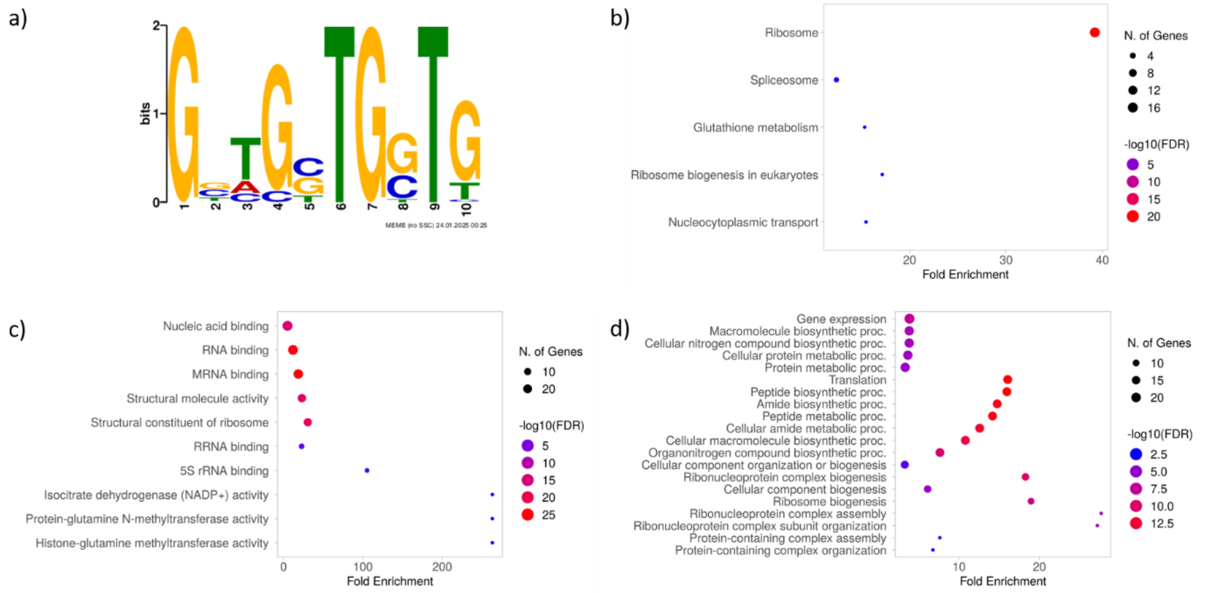

**Fig. S33:** Characterisation of genes with specific motifs, upregulated in IM Cvi. a) The GGTGCTGGTG motif identified at TSS-15 site. b) The KEGG pathway enrichment. c) The Molecular function enrichment. d) The Biological process enrichment.

#### Proteomic analysis

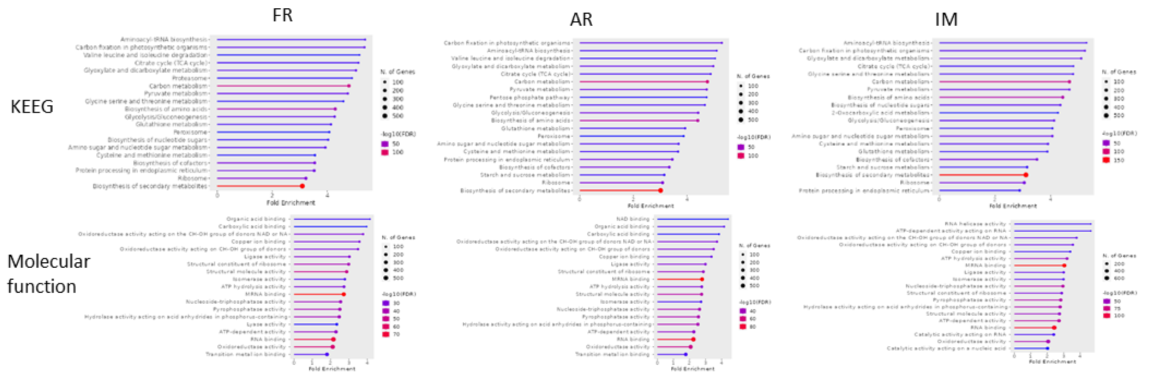

**Fig. S34:** KEEG and Molecular function GO term analysis of proteins contained in a monosomal fraction of freshly harvested (FH), after ripened (AR) and **imbibed seeds (IM) of Col-0.**

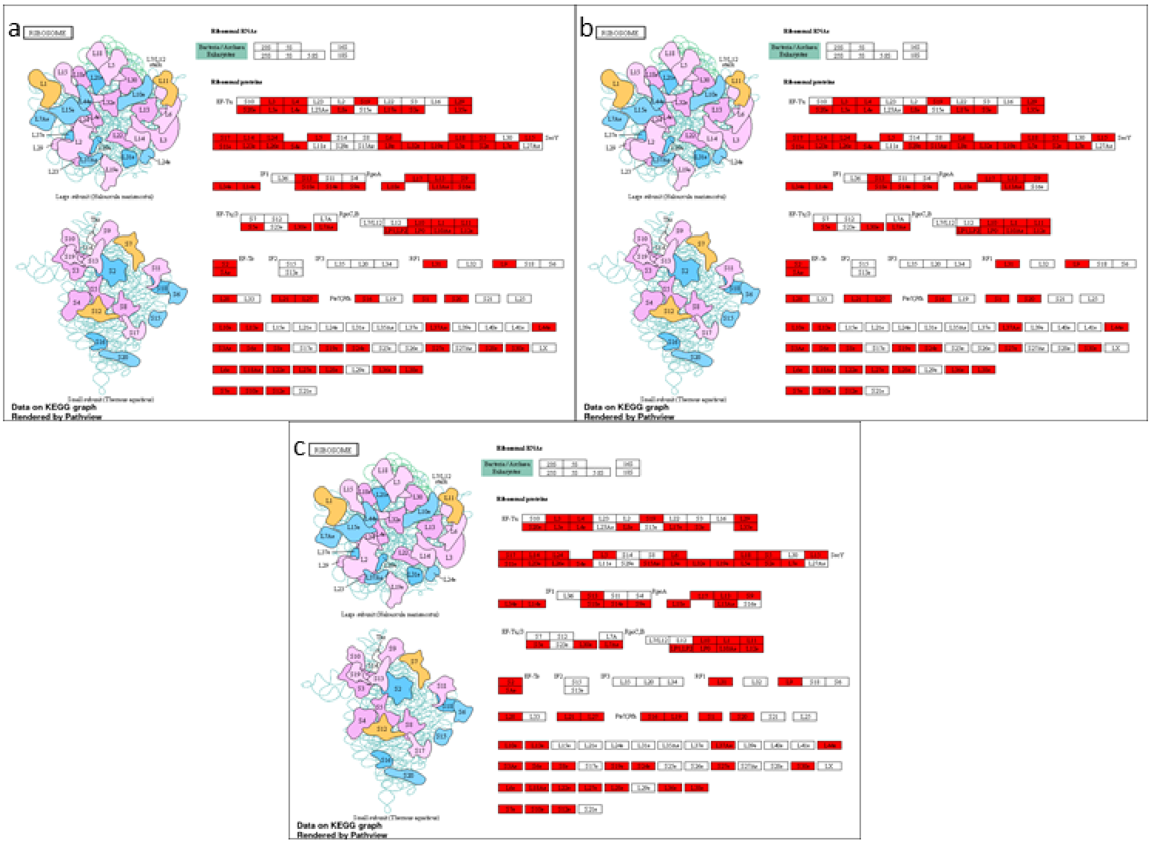

**Fig. S35:** The diagram of the enriched pathway Ribosome with the proteins found in the monosomal fraction of Col-0 freshly harvested (a), after ripened (b) and imbibed (c) seed highlighted in red.

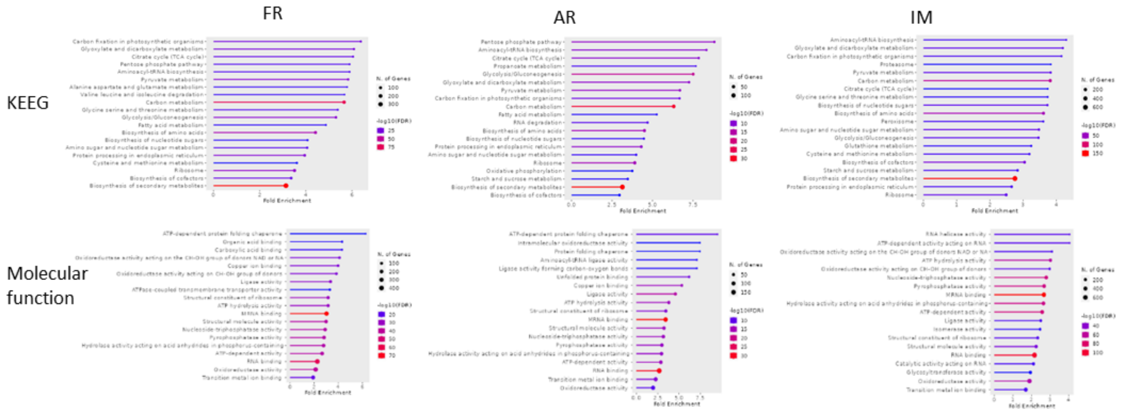

**Fig. S36:** KEEG and Molecular function GO term analysis of proteins contained in a polysomal fraction of freshly harvested (FH), after ripened (AR) and imbibed seeds (IM) of Col-0.

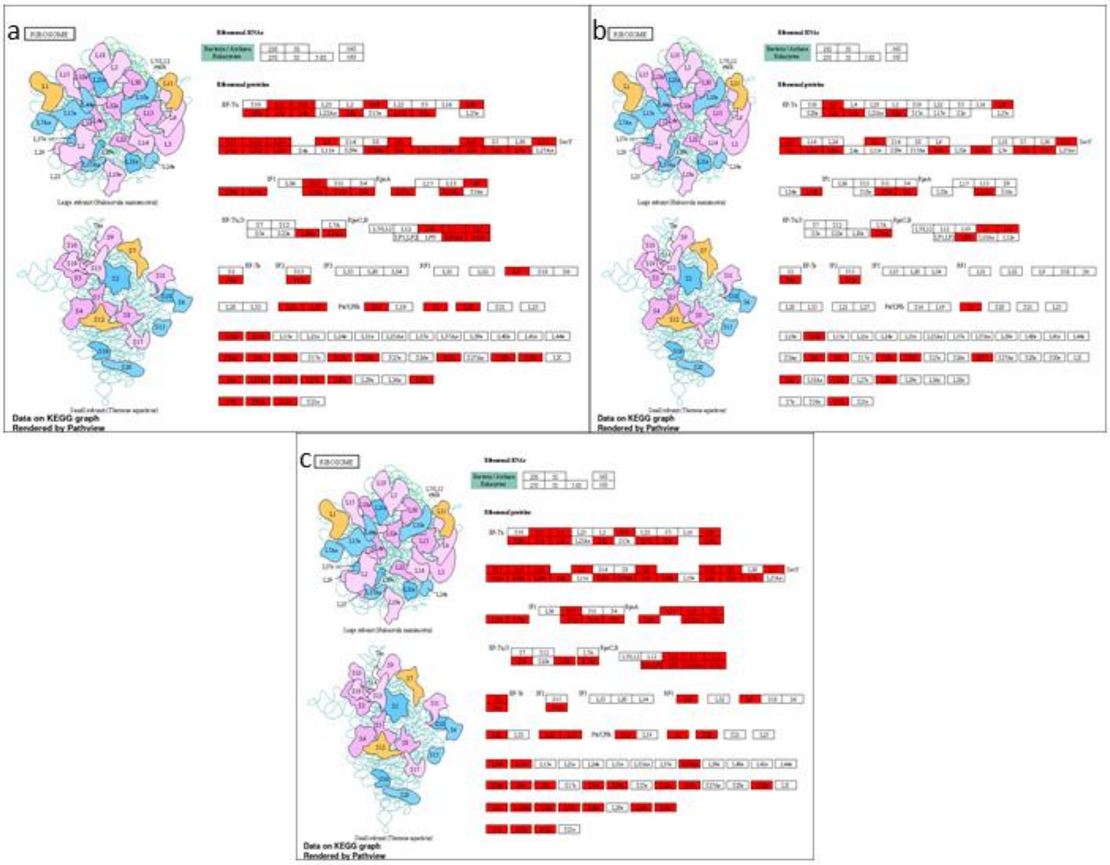

**Fig. S37:** The diagram of the enriched pathway Ribosome with the proteins found in the polysomal fraction of Col-0 freshly harvested (a), after ripened (b) and imbibed (c) seed highlighted in red.

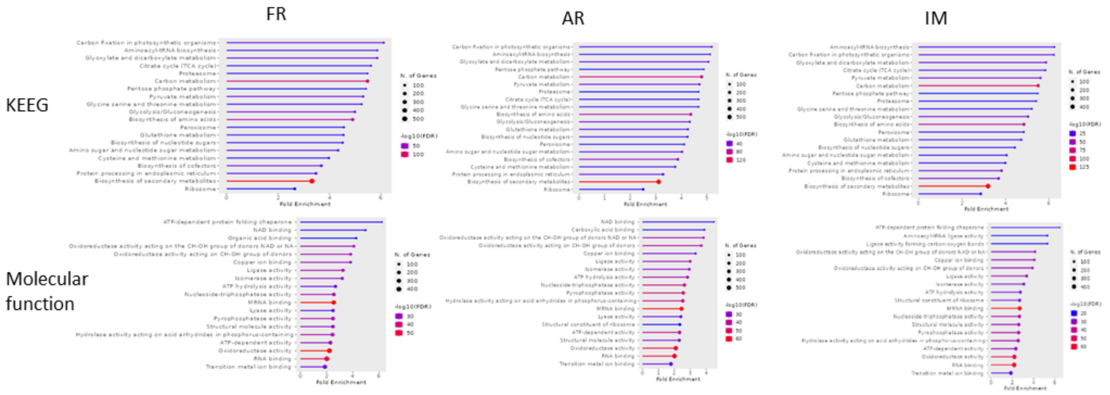

.

**Fig. S38:** KEEG and Molecular function GO term analysis of proteins contained in a monosomal fraction of freshly harvested (FH), after ripened (AR) and imbibed seeds (IM) of Cvi-0**.**

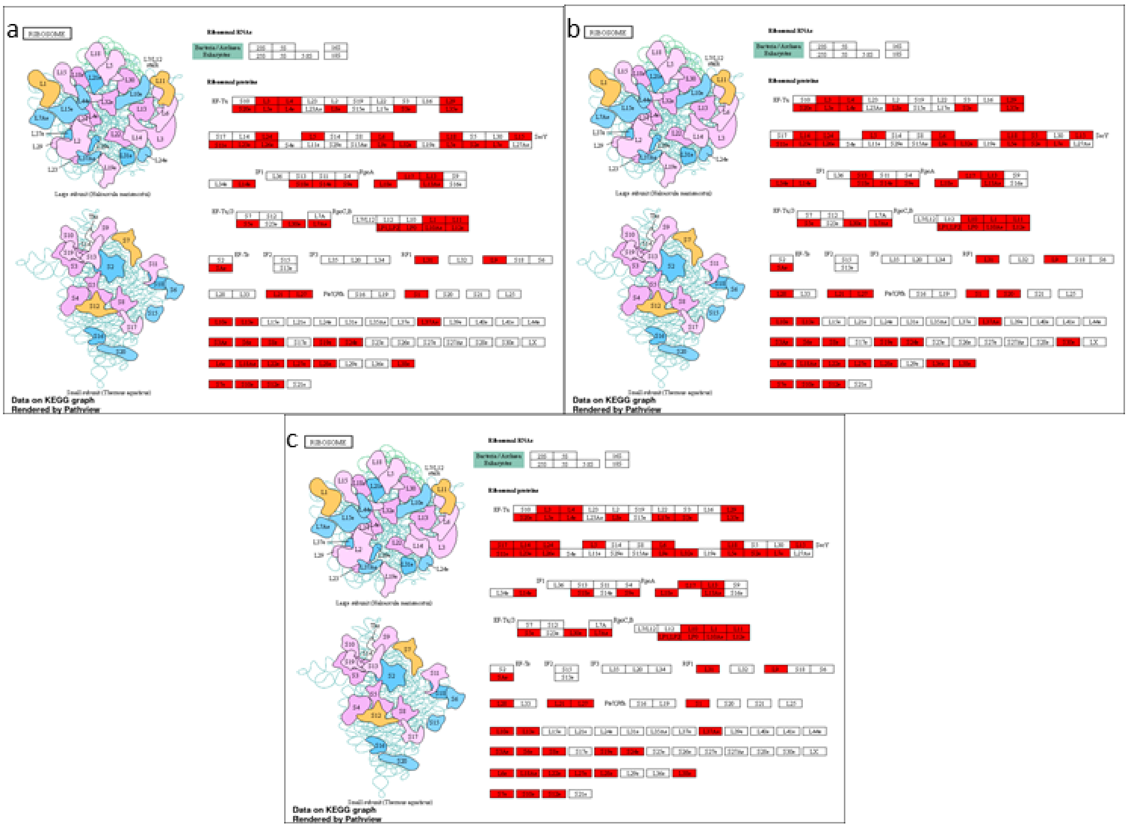

**Fig. S39:** The diagram of the enriched pathway Ribosome with the proteins found in the monosomal fraction of Cvi-0 freshly harvested (a), after ripened (b) and imbibed (c) seed highlighted in red.

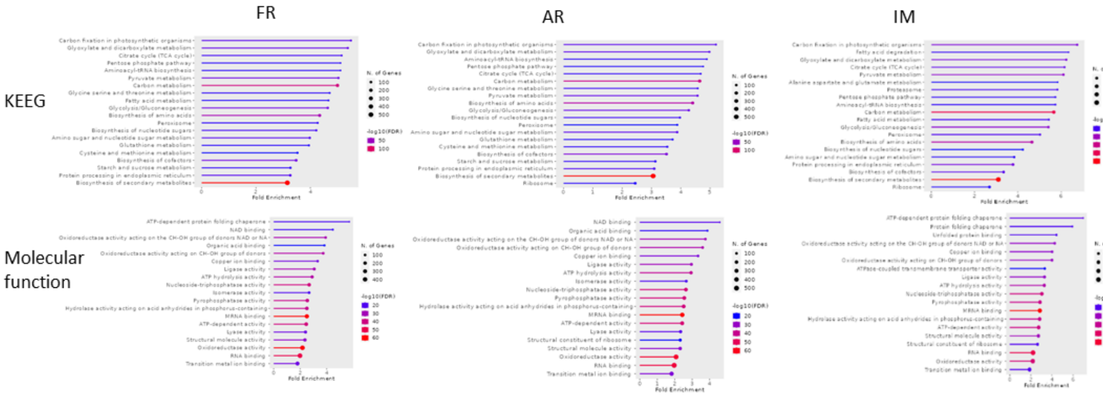

**Fig. S40:** KEEG and Molecular function GO term analysis of proteins contained in a polysomal fraction of freshly-harvested (FH), after-ripened (AR) and imbibed seeds (IM) of Cvi-0.

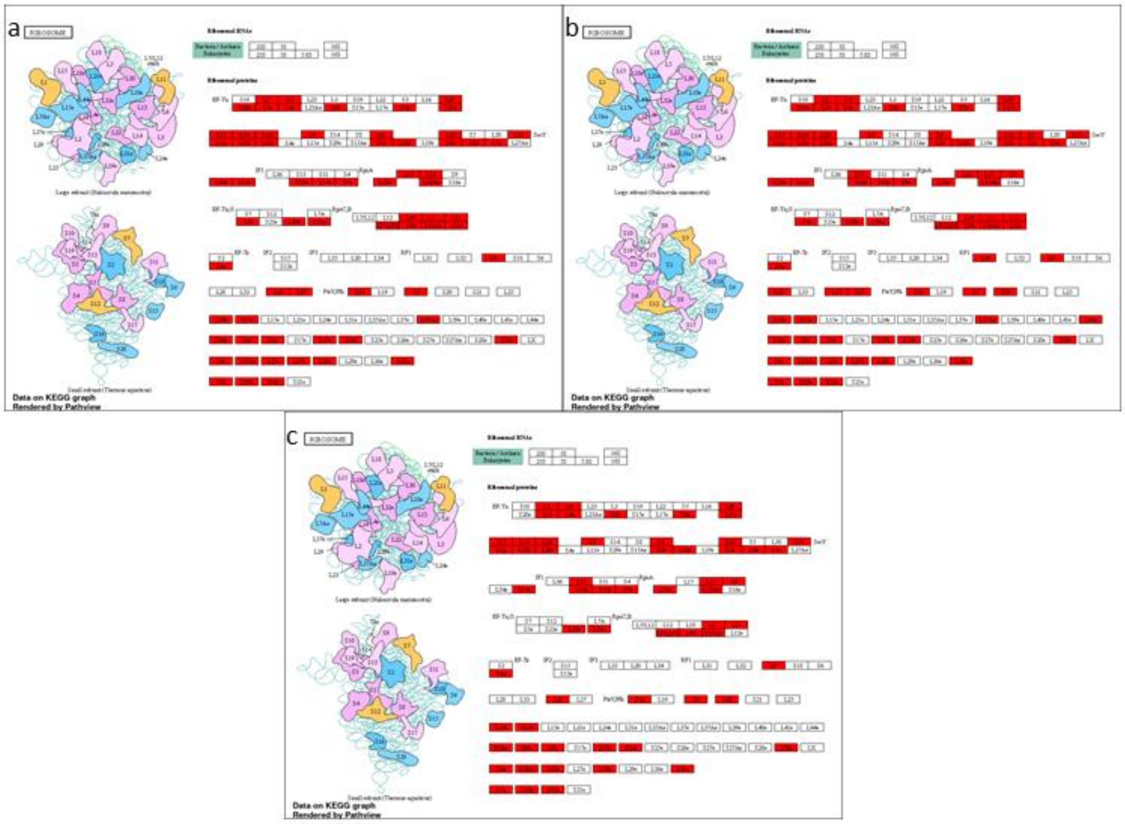

**Fig. S41:** The diagram of the enriched pathway Ribosome with the proteins found in the polysomal fraction of Cvi-0 freshly harvested (a), after ripened (b) and imbibed (c) seed highlighted in red.
